# A compact three-way junction oligonucleotide probe for widefield and super-resolution imaging of RNAs and proteins

**DOI:** 10.64898/2026.11.01.763471

**Authors:** Phuong Nguyen, Hongqiang Ma, Maomao Chen, Xuejiao Sun, Sihan Li, Adam C. Soloff, Da Yang, Yang Liu

**Author notes:** Corresponding author. Yang Liu.

## Abstract

Molecular imaging of RNAs and proteins requires methods that preserve nanoscale organization while enabling large-scale tissue analysis. However, weak or low-abundance targets often need signal amplification, which can increase labeling size, background, or signal spread and compromise super-resolution imaging. Here, we report a compact three-way junction (3WJ) oligonucleotide probe platform for multiscale RNA and protein imaging. The design couples paired target recognition with junction assembly and recruitment of preassembled amplifier complexes, while tunable amplification balances signal strength and labeling size. Using DNA origami and single-molecule localization microscopy, we show enhanced signal while retaining the ability to resolve nanoscale target patterns. The platform improves RNA detection and supports simultaneous RNA-protein super-resolution imaging in cells. It also enhances protein detection in formalin-fixed, paraffin-embedded tissues under low-numerical-aperture, large-field imaging conditions. Together, this programmable 3WJ platform enables sensitive RNA and protein detection across scales, supporting super-resolution analysis of molecular organization and large-field tissue mapping.

## INTRODUCTION

Spatially resolved molecular imaging increasingly requires RNA and protein probes that can operate across imaging scales, from nanoscale molecular organization to cellular phenotypes and large-field tissue architecture. Fluorescence in situ hybridization (FISH) and immunofluorescence imaging are essential tools for visualizing RNAs and proteins within their native spatial environments. However, imaging across these scales creates a fundamental probe-design challenge. Super-resolution microscopy requires compact probes that minimize linkage error and preserve spatial precision, whereas high-throughput tissue mapping and detection of low-abundance targets require strong signal amplification with minimal nonspecific binding. Although amplification can improve detectability, it can also increase effective probe size, apparent signal spreading, and nonspecific background, all of which can compromise super-resolution imaging. Thus, a central unmet need is a probe architecture that can operate across spatial scales: compact enough for molecular-scale super-resolution imaging, bright and specific enough for low-abundance RNA and protein detection, and scalable enough for large-field-of-view tissue mapping.

Single-molecule fluorescence in situ hybridization (smFISH) (*1, 2*) remains one of the most widely used approaches for RNA imaging due to its ability to detect individual transcripts using fluorescently labeled oligonucleotide probes. Conventional smFISH typically requires tens to hundreds of probes tiled along a target RNA transcript to generate sufficient signal intensity, particularly for low-abundance transcripts (*1, 3–5*). This requirement increases experimental cost and limits detection of short RNAs, partially degraded transcripts, or targets containing only a limited number of accessible binding sites. These limitations become particularly important for applications involving low-abundance and spatially confined RNA targets, as well as multiplexed RNA–protein imaging within crowded nuclear environments.

To improve sensitivity while reducing the number of target-binding probes, a variety of amplification strategies have been developed, including tyramide signal amplification (TSA) (*6–8*), enzyme-based rolling-circle amplification (RCA) (*9–13*), signal amplification by exchange reaction (SABER) (*14, 15*), hybridization chain reaction (HCR) (*16*), click chemistry-based amplification (*17, 18*), and branched DNA (bDNA) approaches (*19–21*). Among these, bDNA approaches offer a modular, enzyme-free framework for signal amplification, using successive layers of oligonucleotide hybridization to recruit multiple fluorescent reporters to each target. Target recognition and amplification can be designed separately, while the number of reporter-binding sites provides control over amplification capacity. However, amplifier sequences require careful optimization to minimize non-specific binding and inefficient assembly, and sequential recruitment of amplification layers adds hybridization and washing steps (*19*). These considerations motivate bDNA-based designs that simplify assembly within the specimen while maintaining specificity and controlling the spatial extent of labeling. More broadly, structurally defined DNA nanostructures (*22*) offer control over fluorophore stoichiometry and spatial organization, potentially enabling compact signal amplification strategies with tunable fluorescence output.

Specificity is equally important because amplification can magnify signals arising from non-specific probe binding, converting rare off-target events into detectable signals that may be difficult to distinguish from true target signal. In conventional smFISH, signals from multiple probes bound to the same transcript help distinguish true signals from background (*1*). Several strategies further improve specificity by requiring coordinated recognition of the intended target. Two-color colocalization approaches identify transcripts using independently labeled probe sets and distinguish true signals by their spatial coincidence (*1, 23, 24*). Split-FISH (*25*) and the double-Z design used in RNAscope (*26*) couple signal generation to adjacent binding of paired probes, reducing signals from isolated probe-binding events. The recently reported π-FISH (*21*) similarly uses paired probes, incorporating complementary base pairs between the probes to enhance hybridization stability. The paired probes jointly recruit a secondary amplifier, followed by sequential amplification and fluorescent readout. This work establishes a precedent for combining inter-probe base pairing with amplified split-probe detection. However, achieving strong amplification while maintaining specificity and limiting the size of the labeling assembly remains an important design challenge, particularly when sensitive detection must be combined with nanoscale imaging. This motivates probe designs that bring paired target recognition and stable, specific amplifier recruitment within a single compact assembly.

Three-way junction (3WJ) formation provides a compact framework for organizing multiple oligonucleotide interactions within a compact assembly. Structural studies have shown that 3WJs form compact branched, Y-like architectures, that brings three oligonucleotide arms together within a defined local geometry (*27, 28*). This architecture offers a basis for coupling target recognition with localized assembly, while its compact organization is attractive for nanoscale imaging. Building on this principle, we developed a compact 3WJ oligonucleotide probe with tunable signal amplification for multiscale imaging of RNAs and proteins. Our design combines short amplifier-binding regions with preassembled amplifier complexes. Amplification can be adjusted to target abundance and imaging requirements, supporting both sensitive widefield detection and super-resolution imaging.

We characterize probe architecture, amplification and specificity using DNA origami and single-molecule localization microscopy, then demonstrate detection of diverse RNAs, including low-abundance transcripts. We extend the platform to protein detection through 3WJ-enabled immunofluorescence (3WJ-IF) and combine RNA and protein labeling for two-color super-resolution imaging in cells. Finally, we demonstrate amplified protein detection in formalin-fixed paraffin-embedded (FFPE) tissues using large-field fluorescence imaging with low-NA optics. Together, these results establish a common probe platform for RNA and protein detection across scales, from nanoscale organization in cells to large-area tissue mapping.

## RESULTS

### Design principle of the 3WJ probe for target-dependent and tunable signal amplification

Our compact three-way junction (3WJ)-based probe system couples target recognition and structural assembly to tunable signal amplification, favoring amplifier recruitment when the split probes assemble on adjacent regions of the target (**Fig. 1 and Fig. S1**). The system consists of two split probes, each containing three functional domains: a target-recognition sequence (TRS), a junction-forming sequence (JFS), and a half-binding sequence (HBS) (**Fig. 1A**). The TRS domains recognize adjacent regions of the target, the JFS domains promote 3WJ formation, and the two HBS domains jointly provide a docking interface for recruitment of preassembled amplifier complexes (**Fig. 1B**). When the TRS domains of both split probes hybridize to adjacent target regions, their JFS domains are brought into proximity for complementary base pairing, favoring 3WJ formation. The resulting assembly positions the two HBS domains to jointly provide a docking interface for amplifier recruitment. In this way, target binding, 3WJ assembly, and amplifier recruitment are designed to work together to support efficient signal generation by target-bound probe pairs while limiting signals from isolated probe-binding events. However, improved target recognition and junction assembly do not necessarily eliminate false-positive signal generation when signal amplification is introduced. To limit amplifier recruitment by incomplete probe configurations, each split probe carries a short, 9-nucleotide (nt) HBS and the two segments together form the docking interface recognized by the primary amplifier. To limit potential secondary structure formation, the HBS sequences were designed using a restricted nucleotide composition, in which each position contains only A, T, or C nucleotides, excluding G (H = A/T/C). As shown in **Fig. S2**, H-HBS sequences produced stronger signals than N-HBS designs (in which each HBS position can contain A, T, C, or G) for both RNA (*GAPDH* in HCT116 cells) and protein detection (RNAPII in MCF7 cells).

**Fig. 1.**
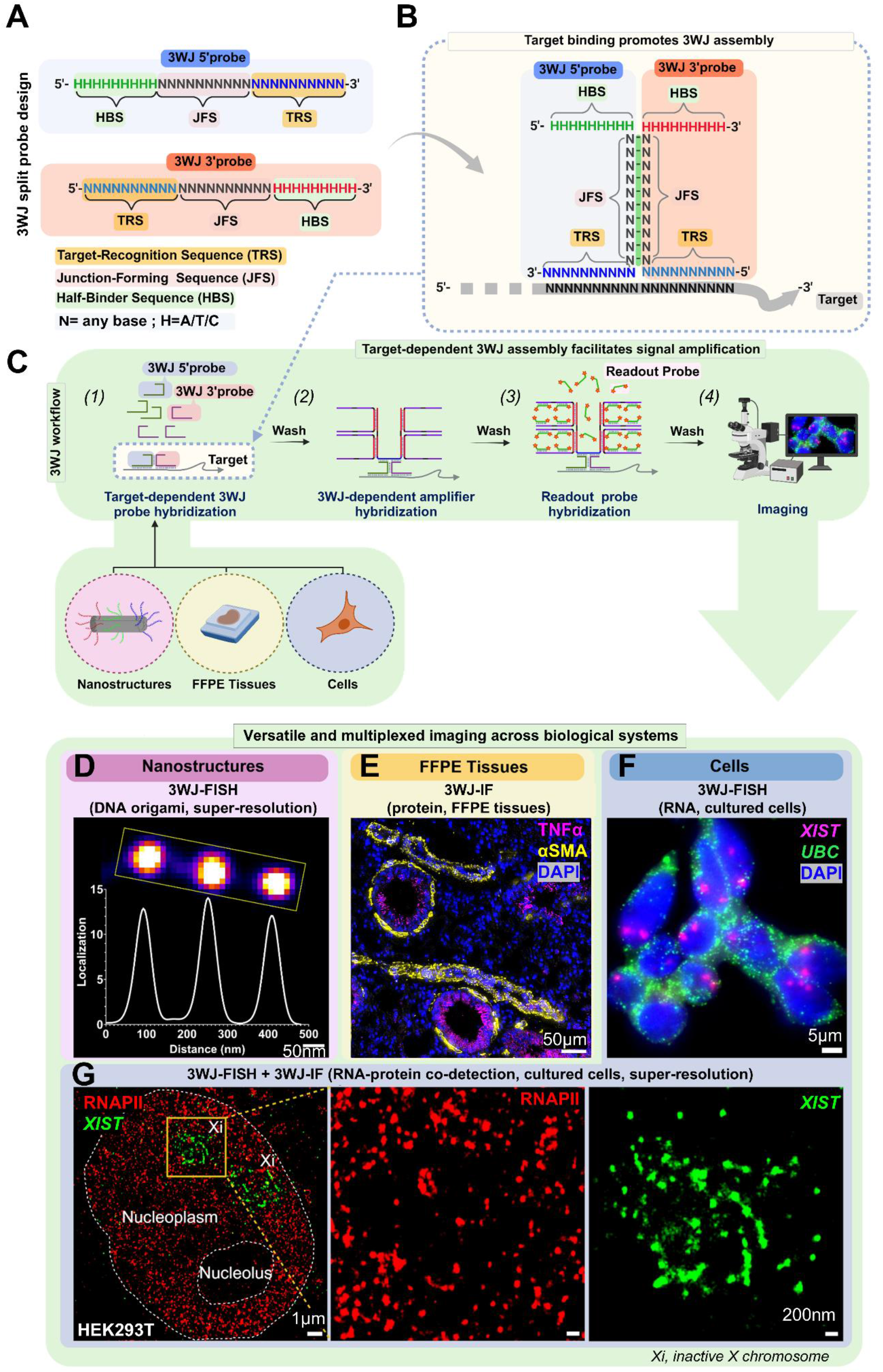
Design and workflow of the 3WJ-based split-probe amplification platform. (**A**) Schematic of the split 3WJ probe design. Each probe contains a target-recognition sequence (TRS), a junction-forming sequence (JFS), and a half-binding sequence (HBS). N denotes A/C/G/T and H denotes A/T/C. (**B**) Binding of the split probes to adjacent target regions brings their JFS domains into proximity, favoring 3WJ formation and positioning the two HBS domains for amplifier recruitment. (**C**) Workflow of the 3WJ imaging strategy. (1) Split probes first hybridize to adjacent regions of the target. (2) Target-dependent 3WJ formation positions the paired HBS docking interface for recruitment of preassembled primary-secondary amplifier complexes. (3) Fluorescent readout probes hybridize to the amplifier modules to generate signal. (4) The amplified signals are imaged by fluorescence microscopy. (**D-G**) Representative applications of the 3WJ platform. Examples include (D) super-resolution imaging of six-helix bundle (6HB) DNA origami nanostructures; (E) protein detection in FFPE tissues, including tumor necrosis factor alpha (TNFα) and alpha-smooth muscle actin (αSMA), with nuclei counterstained by 4′,6-diamidino-2-phenylindole (DAPI). Additional images, including a larger field of view, are shown in Supplementary Fig. S17. (F) RNA detection in cells, including X-inactive specific transcript (*XIST*) and ubiquitin C (*UBC*); and (G) two-color super-resolution imaging of *XIST* RNA and RNA polymerase II (RNAPII). Xi indicates the inactive X chromosome.

We next integrated the 3WJ probe design into a modular amplification workflow (**Fig. 1C**). In the first step, the two split probes hybridize to adjacent regions of the target and assemble into the 3WJ structure. In the second step, a preassembled amplifier complex is recruited through a paired HBS docking interface. Because the primary and secondary amplifiers are preassembled before target labeling (**Fig. S1B**), they are recruited together to the target-bound 3WJ probe in a single hybridization step (**Fig. S1C**). This design reduces the number of sequential hybridization and washing steps within the sample. In the third step, fluorescent readout probes hybridize to the amplifier scaffold to generate detectable signal, followed by standard washing and imaging (**Fig. 1C(*i–iv*)**).

For the standard 16× amplification configuration, both the primary and secondary (1° and 2°) amplifiers were designed as 100-nt oligonucleotides composed of multiple binding domains, each approximately 18-nt long, separated by spacer regions (**Fig. S1A-B**). To stabilize the interactions between the primary and secondary amplifiers, locked nucleic acids (LNA) were incorporated into their complementary binding domains (**Table S1**; **Fig. S3A**). The primary and secondary amplifiers were pre-assembled by controlled annealing before use and stored frozen until hybridization. In this assembled configuration (**Fig. S1B**), the single-stranded regions required for 3WJ docking and readout-probe binding remain available for hybridization. The 3WJ docking domain of the primary amplifier was designed to minimize internal secondary structure and facilitate hybridization with the paired HBS docking interface on the target-bound 3WJ probe (**Fig. S1C**). In RNAPII detection by 3WJ-IF, pre-assembled LNA-containing amplifiers produced stronger fluorescence signals than their LNA-free counterparts (**Fig. S3**), consistent with a beneficial contribution of LNA incorporation to signal amplification.

We demonstrated this approach across diverse sample types, labeling targets, and spatial scales, including DNA nanostructures, cultured cells, and FFPE tissue sections (**Fig. 1C–D**). At the nanoscale, 3WJ-FISH resolved programmed probe-binding sites on DNA origami nanostructures by direct Stochastic Optical Reconstruction Microscopy (dSTORM), demonstrating compatibility with super-resolution imaging (**Fig. 1D*(i)*** and **Fig. S5**). In FFPE tissue sections, 3WJ-based immunofluorescence (3WJ-IF) enabled multiplexed protein detection with distinct spatial patterns of tumor necrosis factor alpha (TNFα) and alpha-smooth muscle actin (αSMA) in inflamed lung tissue **(Fig. 1D*(ii)***). In cultured HEK293T cells, 3WJ-FISH enabled multiplexed RNA detection, including the long noncoding RNA *XIST* in the nucleus and the mRNA *UBC* in the cytoplasm, illustrating multiplexed RNA imaging across transcript classes (**Fig. 1D(*iii*)**). The same amplification strategy also supported RNA–protein co-detection by super-resolution microscopy, revealing nanoscale spatial organization of XIST RNA and RNAPII (**Fig. 1D(*iv*)**). Together, these results illustrate the multiscale imaging capabilities of the 3WJ platform, which are systematically validated in nanoscale imaging, RNA detection, protein detection, and RNA-protein spatial analysis, as detailed below.

### 3WJ probe architecture and pre-assembled amplification strategy enhance signal generation while supporting nanoscale imaging

We next investigated how probe architecture influences the performance of the 3WJ-based platform. In this system, the split-probe pair serves as the interface that couples target recognition to downstream signal amplification. An optimal architecture must support target-dependent assembly, efficiently recruit the amplifier, preserve a compact probe geometry for super-resolution imaging, and minimize signal from isolated or nonspecific probe-binding events.

To define the architectural features required for this balance, we systematically compared split-probe designs that differed in junction geometry and the number of complementary bases in the JFS. Our primary 3WJ design contains a 10-nt complementary JFS, which was designed to maintain a compact probe architecture while supporting sufficient stability for amplifier recruitment. As shown in **Fig. 2A**, we compared this 3WJ design with two JFS-modified variants within the same overall 3WJ framework: a 4C design, in which only four nucleotides in the JFS are complementary, and a 0C design, in which the JFS region is replaced with non-complementary poly(A) sequences. Thus, the 4C and 0C designs allowed us to assess how reducing JFS-mediated base pairing affects signal generation. We further compared the 3WJ design with the previously reported π-shaped (*21*) and circular split-probe geometries (*25*), which represent alternative proximity-dependent architecture for split-probe assembly, as shown in **Fig. S4**.

**Fig. 2.**
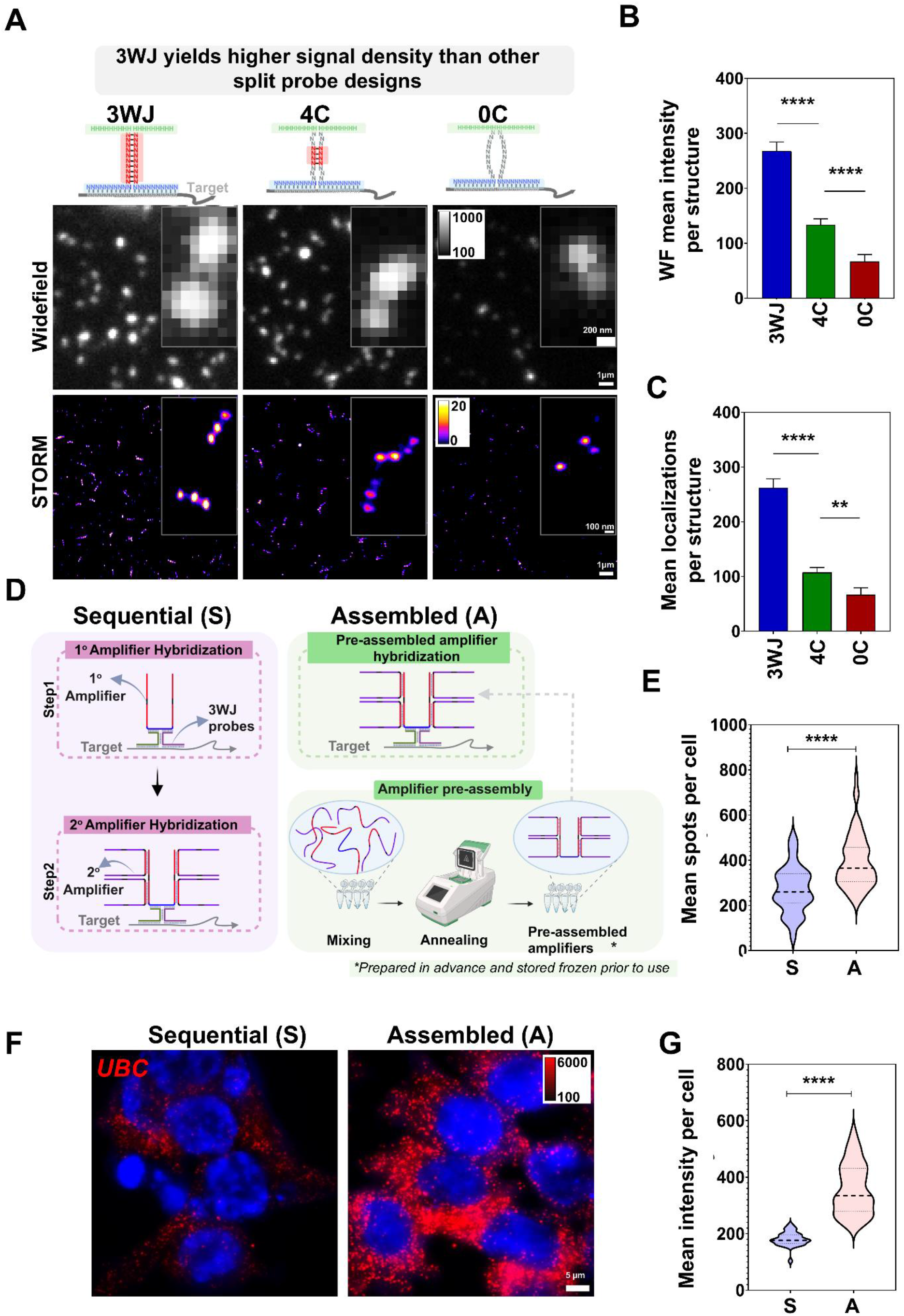
Optimization of split-probe architecture and amplification strategy. (**A**) Comparison of split-probe designs differing in the number of complementary bases of the junction-forming sequence (JFS), including the 10-nt complementary 3WJ design, four-connector (4C) design with four complementary bases, and zero-connector (0C) design with no designed complementary base pairing in the connector region. The 4C and 0C variants retain the same overall probe framework. DNA origami structures contain three spots separated by approximately 168 nm, with six target sites per spot. Representative widefield (WF) and super-resolution (STORM) images are shown with corresponding magnified views. Within each imaging modality, images were displayed using consistent intensity scaling to enable direct visual comparison. Intensity scales and scale bars are indicated in the images. (**B–C**) Quantification of signals for different connector designs, measured as mean WF intensity per structure (**B**) and mean STORM localization number per 3-spot DNA origami structure (**C**). Data are presented as mean ± SD. Statistical significance was determined by Welch’s one-way ANOVA with Dunnett’s T3 multiple comparisons test (****p < 0.0001). (**D**) Schematic illustration of amplification strategies. Sequential amplification (S) involves stepwise hybridization of LNA-free primary and secondary amplifiers, whereas assembled amplification (A) uses pre-assembled LNA-containing primary-secondary amplifier complexes to enable one-step hybridization to the target-bound 3WJ probe. (**E–G)** Detection of *UBC* RNA transcripts in HCT116 cells using sequential (S) and assembled (A) amplification strategies. (**F**) Representative images. Intensity scaling is consistent across conditions, and both intensity scales and scale bars are indicated in the images. (**G**) Violin plots with median and quartiles indicated showing the distribution of mean intensity per cell. (**E**) the number of detected RNA spots per cell. Statistical significance was determined by a two-tailed Welch’s t-test (****p < 0.0001).

To enable a quantitative and unbiased comparison, we used DNA origami structures with a predefined number and spatial organization of target sequences. Each 6-helix bundle (6HB) (*29*) structure contained a total of 18 single-stranded DNA target sites (6 sites per spot × 3 spots) with inter-spot spacing of ∼168 nm. **Figure 2A** shows both conventional diffraction-limited wide-field fluorescence images and corresponding STORM-based super-resolution images. In wide-field images, the three target spots appeared as a diffraction-limited and unresolved cluster, whereas super-resolution imaging resolved the nanoscale organization of the origami structure, with an estimated inter-spot distance of 153 ± 11 nm, compared with the designed spacing of approximately 168 nm (**Fig. S5**).

Among the JFS variants, the 3WJ design produced stronger fluorescence signals and more distinct spots in both wide-field and super-resolution images. In STORM images, the 3WJ probes resolved adjacent spots within the 3-spot origami structure. In contrast, the 4C design showed reduced signal intensity and partial loss of the expected spatial pattern, whereas the 0C design failed to resolve the expected spot organization (**Fig. 2A**). Similar trends were observed when 3WJ design was compared with π-shaped and circular split-probe designs, both of which showed reduced fluorescence signal and partial loss of the 3-spot DNA origami structure (**Fig. S4B**). Quantitative analysis further confirmed that the 3WJ architecture generated substantially higher signal, as measured by mean fluorescence intensity per diffraction-limited spot in widefield images and mean localization counts per resolved three-spot DNA origami structure in STORM images (**Fig. 2B–C**). The same trend was observed across all other tested split-probe geometries, which produced lower fluorescence intensity and localization number per structure relative to the 3WJ configuration (**Fig. S4C–D**). Importantly, all comparisons were performed under identical experimental conditions, including probe concentration, hybridization time, amplification components, imaging parameters, and analysis thresholds. Together, these results demonstrate that the 3WJ design with a 10-nt complementary JFS improves signal generation relative to other JFS variants while maintaining the ability to resolve the nanoscale DNA origami pattern.

Having established that the 3WJ JFS architecture improves signal generation on defined targets, we next examined whether the mode of amplification delivery affects detection performance. We compared a stepwise sequential amplification (S) using LNA-free amplifiers, in which primary and secondary amplifiers were introduced in separate hybridization steps, with an assembled amplification strategy (A) using LNA-containing amplifiers, in which the primary and secondary amplifiers were pre-assembled by controlled annealing before hybridization to the target-bound 3WJ probes (**Fig. 2D**). The assembled strategy was designed to reduce the number of on-sample hybridization steps and to promote formation of the intended amplifier complex before target binding. In this configuration, the pre-assembled complex is recruited to the target-bound 3WJ probe through the primary amplifier, while the secondary amplifier provides additional readout-binding sites for fluorescence signal generation.

We further evaluated whether this assembled amplification strategy improved RNA detection in cells. As shown in **Fig. 2F**, assembled amplification produced stronger signals for *UBC* RNA spots in HCT116 cells than sequential amplification. Quantitative analysis showed significant increases in both mean signal intensity and the number of detected *UBC* RNA spots per cell with the assembled approach (**Fig. 2E, G**). These results demonstrate improved detection performance with the pre-assembled, LNA-containing amplification strategy. By combining a target-dependent 3WJ with this amplification system, the platform supports enhanced signal generation within a simplified workflow while preserving compatibility with nanoscale imaging.

### 3WJ-FISH enables enhanced sensitivity and tunable signal amplification

We next evaluated the performance of 3WJ-FISH for RNA detection and its ability to provide quantitative and tunable signal output (**Fig. 3**). Compared with conventional smFISH, an established single-molecule RNA detection method, 3WJ-FISH produced substantially stronger signals and higher spot counts for *UBC* mRNA in HeLa cells (**Fig. 3A–C**), consistent with enhanced detection sensitivity. Notably, the 3WJ-FISH probe pairs were designed to target transcript regions overlapping those used for smFISH. The target-recognition sequences (TRS) in the split probes were selected from corresponding smFISH probe binding sites, allowing a more direct comparison of detection performance between the two approaches. Although the smFISH probe set included additional binding sites, all measurements were performed under matched imaging and analysis conditions, ensuring a direct comparison between the two approaches.

**Fig. 3.**
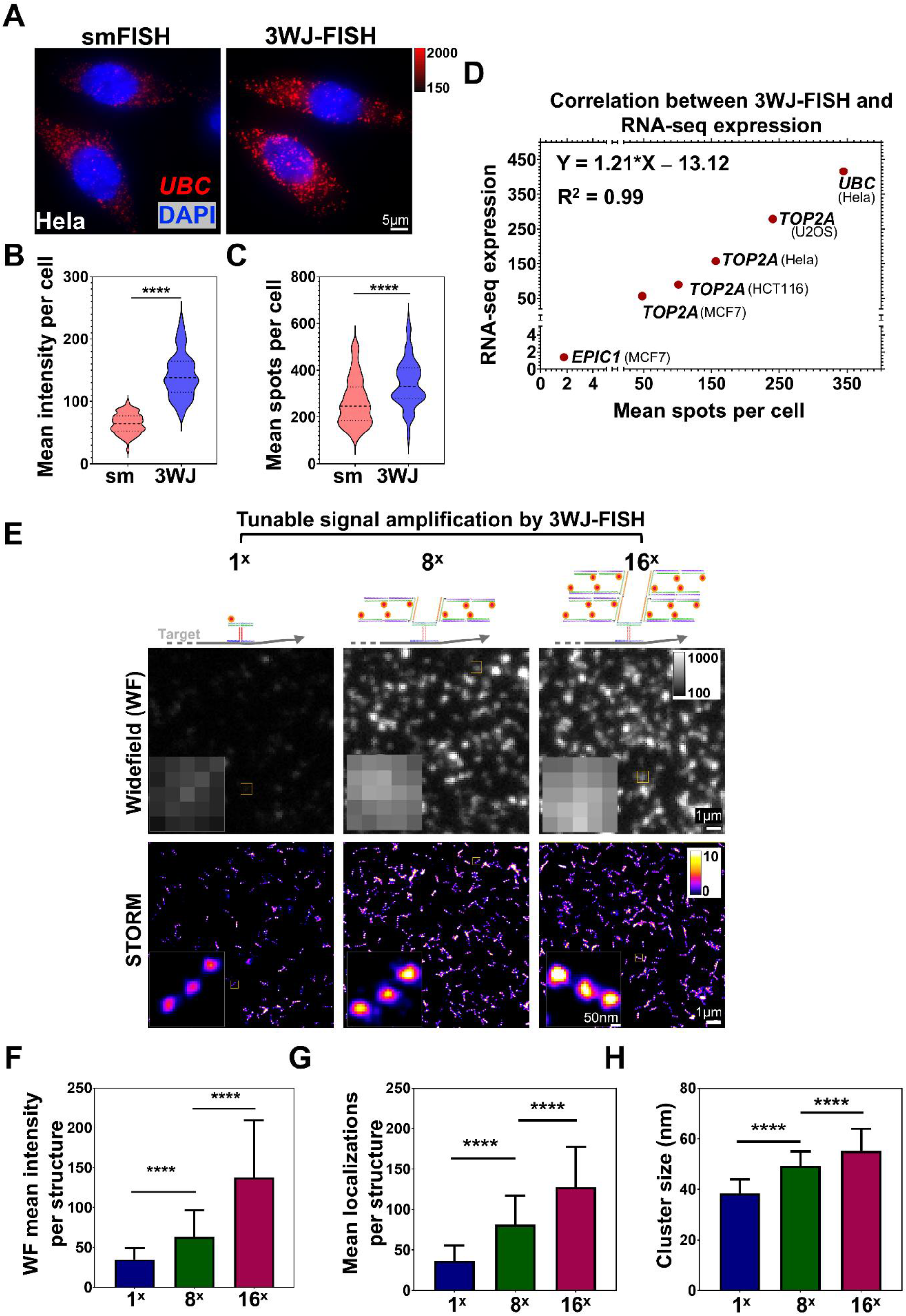
Enhanced RNA detection and tunable signal amplification by 3WJ-FISH. (**A**) Representative images of *UBC* mRNA detection in HeLa cells using smFISH and 3WJ-FISH (with 16× amplification). Nuclei are counterstained with DAPI. Scale bar, 5 μm. (**B–C**) Quantification of *UBC* mRNA detection by smFISH and 3WJ-FISH. Distribution of mean fluorescence intensity per cell (**B**) and mean RNA spots per cell (C) for smFISH (sm) and 3WJ-FISH (3WJ), shown as violin plots with median and quartiles indicated. Statistical significance was determined using a two-tailed Welch’s t-test (\*\*\*\**P* < 0.0001). (**D**) Correlation between 3WJ-FISH measurements (mean spots per cell) and RNA-seq expression levels (transcripts per million or TPM) across multiple genes and cell lines. Linear regression analysis shows strong agreement (R² = 0.99). Mean RNA spot counts per cell used for correlation analysis are summarized in **Table S13**, and the corresponding single-cell distributions are provided in **Fig. S12**. Comparison with smFISH and representative images are shown in **Fig. S7-11**. (**E**) Effect of 3WJ-FISH signal amplification on the super-resolved nanostructure using increasing amplification levels of 1×, 8×, and 16× on 6HB DNA origami structures. Representative widefield (top) and STORM (bottom) images with corresponding magnified views are shown. Images were acquired under identical conditions and displayed with consistent intensity scaling to enable direct comparison across amplification levels. Scale bars, 1 μm (widefield) and 1 μm or 50 nm (STORM, as indicated). (**F–H**) Quantification of mean fluorescence intensity (**F**), mean number of localizations (**G**), and cluster (spot) size (**H**) as a function of amplification level. Data are presented as mean ± SD. Statistical significance was assessed using Welch’s ANOVA followed by Dunnett’s T3 multiple comparisons test (****P < 0.0001).

We next compared 3WJ-FISH with amplified split-probe designs lacking JFS by evaluating circular (CIR-FISH) and double-Z (Z-FISH) architectures for *NEAT1* detection in cells using the same 16× amplification and preassembly procedure. Under these conditions, 3WJ-FISH produced the highest fluorescence intensity per nuclear *NEAT1* spot, while CIR-FISH generated stronger signals than Z-FISH (**Fig. S6**), consistent with the previously reported comparison of circular and double-Z designs (*25*). These results support improved signal generation by the 3WJ architecture within a shared amplification workflow.

To assess quantitative performance, we compared 3WJ-FISH measurements with corresponding RNA-seq expression values across multiple genes and cell lines. RNA-seq expression values were obtained from publicly available transcriptomic datasets for the corresponding genes and cell lines (*30*). The mean RNA spot counts per cell showed a strong correlation with RNA expression levels (R² = 0.99), supporting the quantitative performance of 3WJ-FISH (**Fig. 3D**). The selected targets spanned different RNA classes and abundance ranges, including the lncRNA *EPIC1* (*31*), as well as protein-coding mRNAs such as *TOP2A* and *UBC*. For comparison, conventional smFISH detected fewer *UBC* and *TOP2A* RNA spots than 3WJ-FISH (**Fig. 3A-C**; **Figs. S7-11**). In particular, 3WJ-FISH detected the low-abundance lncRNA *EPIC1* as discrete spots predominantly localized within the nuclei, whereas conventional smFISH produced numerous cytoplasmic signals inconsistent with the low *EPIC1* expression level indicated by RNA-seq data (**Fig. S8; Table S13**). The corresponding single-cell distributions of RNA spot counts and intensities across genes and cell lines from 3WJ-FISH are provided in **Fig. S12**, multiplexed RNA labeling is shown in **Fig. S13**, and summary statistics used for correlation analysis are reported in **Table S13**.

We further examined whether the fluorescence signal could be tuned by varying the level of amplification. This capability is important for multiscale imaging because different imaging contexts impose different signal requirements. Low-abundance targets require stronger amplification to improve detectability, and large-field-of-view imaging can benefit from stronger amplification to compensate for reduced photon collection with lower-NA objectives. In contrast, super-resolution imaging requires sufficient signal without excessive expansion of the apparent target size. We therefore used 6HB DNA origami as a structurally defined nanoscale reference to evaluate how different amplification levels affect both signal strength and the apparent size and separation of labeled target regions.

Increasing the amplifier level from 1× to 8× to 16× produced progressive increases in fluorescence intensity and the number of localizations per structure in both wide-field and super-resolution imaging (**Fig. 3E–G**). Representative STORM images (**Fig. 3E**) of individual 6HB structures showed that the three linearly arranged spots can be resolved, demonstrating that 3WJ-FISH supports compact nanoscale structures while providing tunable signal amplification. At higher amplification levels, the spot size increased (**Fig. 3H**), consistent with greater accumulation of amplified signal at each target site. These results also illustrate an important trade-off: higher amplification improves the signal, whereas lower or intermediate amplification better preserves compact nanoscale structures. Thus, 3WJ-FISH allows amplification conditions to be adjusted according to the imaging goal, from sensitive detection in wide-field imaging to spatially resolved molecular structure via super-resolution imaging. Additional super-resolution imaging of *NEAT1* RNA under intermediate (8×) amplification further supported the nanoscale resolving capability of 3WJ-FISH in cells (**Fig. S14**), revealing partially hollow *NEAT1* localization patterns consistent with reported paraspeckle substructures (*32*). Together, these results demonstrate that 3WJ-FISH enables sensitive, quantitative, and tunable detection across both diffraction-limited wide-field fluorescence and super-resolution imaging modalities.

### Validation of 3WJ probe specificity and suppression of nonspecific amplification

After establishing that 3WJ-FISH supports tunable signal amplification, we next evaluated probe specificity under the intermediate 8× amplification condition. This condition provides sufficient signal for RNA detection while maintaining relatively compact labeling in super-resolution imaging and was used consistently for the specificity experiments (**Fig. 4**). Because amplification can convert rare off-target binding events into detectable signals, minimizing nonspecific amplifier recruitment is essential for accurate RNA detection.

**Fig. 4.**
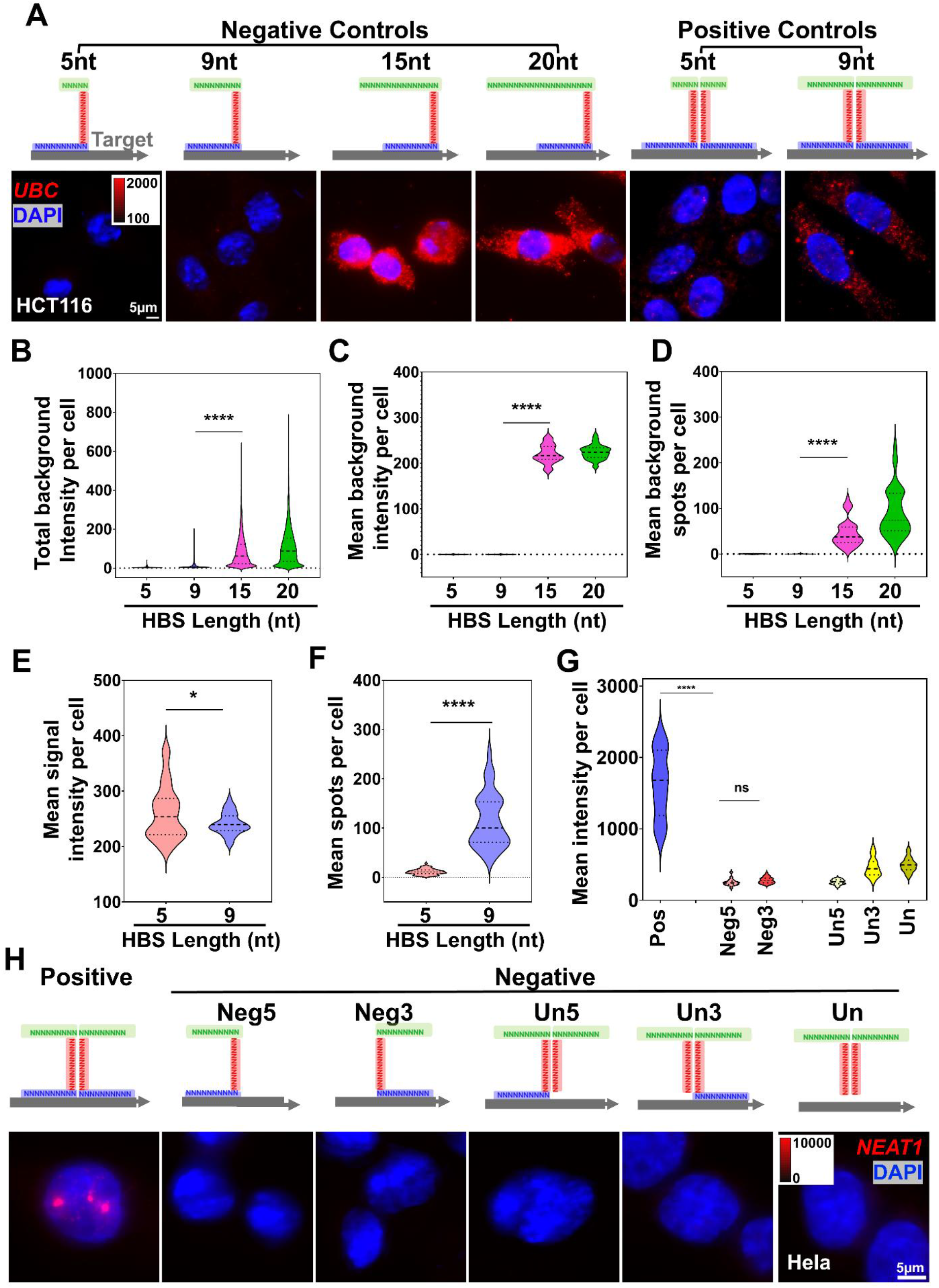
Optimization of half-binding sequences (HBS) length for specific 3WJ-FISH amplification and validation of target-dependent 3WJ-FISH amplification. (**A**) Representative images of UBC-targeted RNA labeling (8× amplification) in HCT116 cells using 3WJ probes with varying HBS lengths (5, 9, 15, and 20 nt), comparing unpaired negative-control probes (left) and fully matched positive-control probes (right). (**B–D**) Background signal across HBS lengths in unpaired negative controls, measured as total fluorescence signal per cell (B), mean fluorescence signal per cell (C), and detected RNA spots per cell (D). Data are shown as violin plots with median and quartiles indicated. Statistical significance was assessed using the Kruskal–Wallis test followed by Dunn’s multiple comparisons test (****P < 0.0001). (**E–F**) Positive signal output for shorter HBS lengths (5 and 9 nt), measured as mean fluorescence signal per cell (E) and mean RNA spots per cell (F), with median and quartiles indicated. Statistical significance was determined using a two-tailed Welch’s t-test (E, \**P* < 0.05; F, ****P < 0.0001). (**G**) Quantification of *NEAT1* RNA detection in HeLa cells across probe configurations, including fully matched positive probes (Pos), unpaired negative controls (Neg5, Neg3), and unbound controls lacking one (Un5, Un3) or both (Un) target-recognition sequence (TRS) domains. The unbound controls retain the complementary 10-nt junction-forming sequences (JFS) and HBS domains. Data are shown as violin plots with median and quartiles indicated. Statistical significance was assessed using Welch’s ANOVA followed by Dunnett’s T3 multiple comparisons test (**P < 0.01; ****P < 0.0001). (**H**) Representative images of *NEAT1* RNA detection in HeLa cells under different control configurations, showing substantially reduced signals in the unpaired and TRS-deletion controls. Identical display settings were used within each experimental set; corresponding intensity scales and scale bars are shown in the images.

We first investigated how the length of the half-binding sequence (HBS), which contributes to the paired docking interface for amplifier recruitment after target-dependent 3WJ assembly, affects non-specific amplification. To assess the contribution of HBS length to background, we used negative control conditions in which only one unpaired 3WJ probe was applied. In these probes, the TRS and JFS were retained, while the HBS length was varied among 5, 9, 15, and 20 nt using sequences composed only of A/T/C – only “H” nucleotides. The design preserved one amplifier-binding segment but prevented assembly of the complete split-probe pair, allowing us to directly assess whether HBS length alone could drive nonspecific amplifier recruitment.

The selected HBS lengths were chosen to test how the length of an individual amplifier-binding segments influences background. Shorter HBS lengths were inspired by prior split-probe designs, where bridge-probe interactions of ∼9 nt or fewer nucleotides suppress nonspecific background from unpaired probes (*25*). Longer HBS lengths (15 and 20 nt) were included to test whether extending the amplifier-binding segment increases background, whereas the 5 nt HBS was used to assess a shorter alternative to the 9-nt design for amplifier recruitment within the 3WJ framework. As shown in **Figs. 4A-D**, in the negative controls, short HBS lengths (5 and 9 nt) resulted in minimal background signal, whereas longer HBS lengths (15 and 20 nt) led to substantially increased nonspecific signal, as measured by total fluorescence intensity, mean intensity, and detected background spot counts per cell. These results are consistent with longer HBS segments allowing amplifier recruitment by individual probes, even in the absence of probe-pair assembly. In contrast, the 9-nt HBS effectively suppressed nonspecific amplification under unpaired probe conditions while retaining sufficient binding capacity for target-dependent amplification. Accordingly, we selected the 9-nt HBS for subsequent probe design.

We next tested whether the 9-nt HBS design suppresses nonspecific amplification when components required for paired target recognition are absent, using the long noncoding RNA *NEAT1* as a model target. Dual-color colocalization of independently labeled probe subsets has been used to validate lncRNA-FISH specificity in a study that included *NEAT1* (*24*). The 3WJ design is intended to favor amplifier recruitment when both split probes bind adjacent target sites and assemble the paired HBS docking interface. To test this requirement, we evaluated a series of control configurations in which individual probes or target-recognition domains were removed. In the unpaired negative controls (Neg5, Neg3), only one probe of the pair was present, preventing complete 3WJ formation. In the unbound configurations (Un5, Un3, Un), one or both probes lacked the TRS domain required for target binding, disrupting paired target recognition while retaining the complementary 10-nt JFS and the HBS domains.

Compared with the fully matched 3WJ probe pair, all control configurations produced negligible signals, including both the unpaired and unbound controls (**Figs. 4G–H**). These results show that retaining either a single unpaired probe or the complementary 10-nt JFS and HBS domains without paired target recognition was insufficient to generate appreciable signal under the tested hybridization and washing conditions. These findings suggest that adjacent target binding brings both probe strands into proximity and helps stabilize the assembled amplifier-docking interface. Loss of either TRS domain removes this contribution from paired target binding, potentially limiting amplifier recruitment or retention despite the presence of complementary JFS and HBS domains. Together, these findings support the contribution of two coordinated design features to background suppression: a short HBS that limits recruitment by unpaired probes, and paired target recognition that brings the junction-forming and amplifier-docking domains together. The 9-nt HBS supported signal generation with the complete target-binding probe pair, while maintaining low background in the unpaired and TRS-deletion controls.

### 3WJ-IF enables oligonucleotide-based amplification for sensitive in situ protein imaging in cells and tissues

Building on the high sensitivity and specificity of 3WJ probes demonstrated in 3WJ-FISH, we next extended this strategy to protein detection using immunofluorescence, termed 3WJ-IF (**Fig. 5A**). In 3WJ-IF, oligonucleotide-conjugated secondary antibodies recruit the 3WJ amplification system after antibody binding to the target protein, replacing conventional fluorophore-labeled secondary antibodies with a programmable oligonucleotide-based amplification strategy. We evaluated this approach using both high-NA microscopy and under ultra-large-field-of-view imaging conditions using an NA = 0.5 objective (*33*). The latter imaging configuration enables broad tissue mapping but has lower photon collection efficiency than high-NA microscopy, making weak protein signals more difficult to detect. Thus, we tested whether 3WJ-based amplification improve protein detection in cells and tissue across these imaging configurations.

**Fig. 5.**
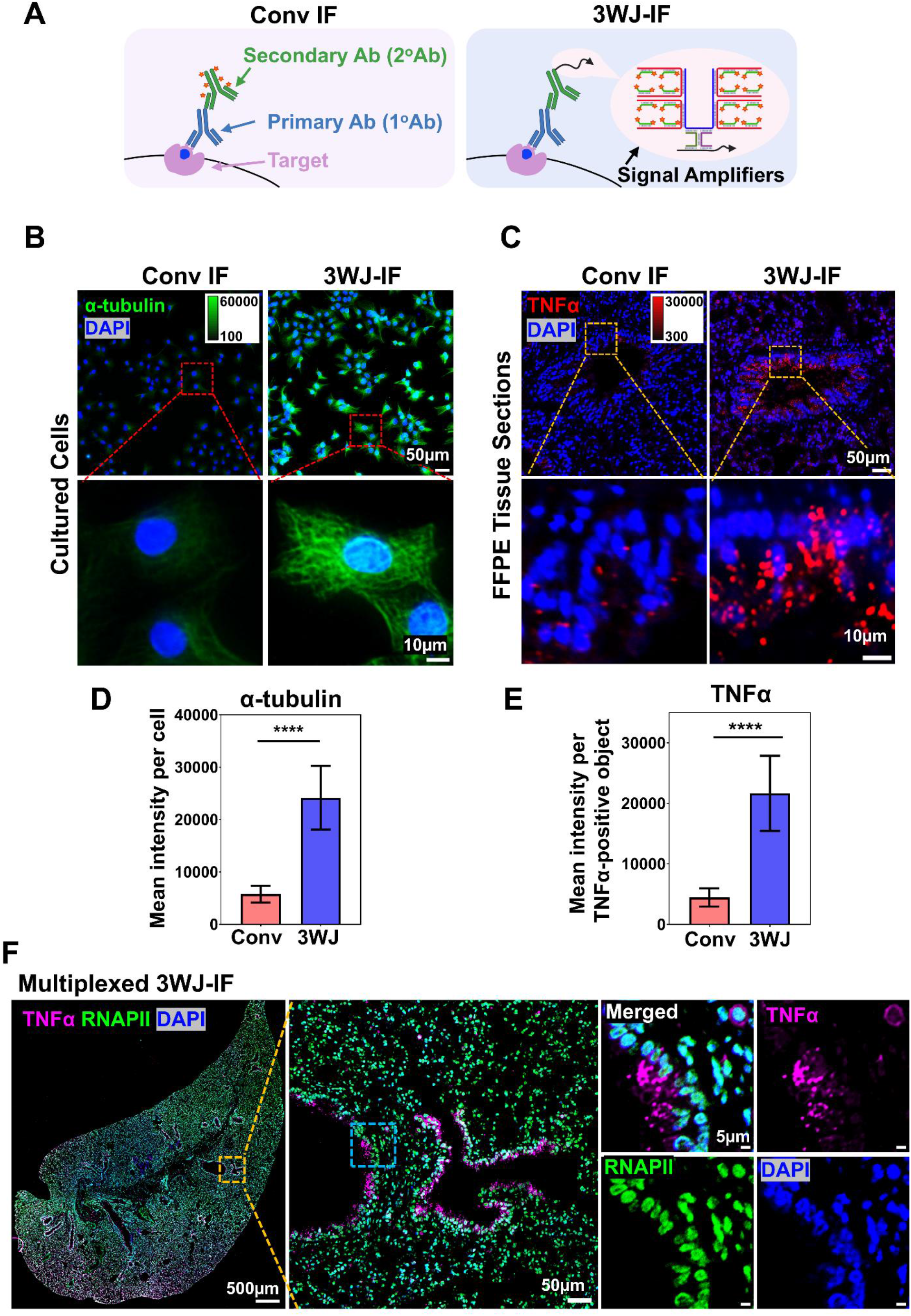
Application of 3WJ-based signal amplification for immunofluorescence (3WJ-IF). (**A**) Schematic comparison of conventional immunofluorescence (Conv IF) and 3WJ-based immunofluorescence (3WJ-IF). In Conv IF, fluorescence signal is generated by fluorophore-conjugated secondary antibodies, whereas in 3WJ-IF, oligonucleotide-conjugated secondary antibodies provide target sequences for 3WJ probe assembly at antibody-bound protein targets. The assembled 3WJ probe recruits oligonucleotide amplifiers, followed by fluorescent readout-probe hybridization for signal generation. (**B-C**) Representative images of α-tubulin staining in cultured cells (B) and TNFα detection in FFPE tissue sections (C) using Conv IF and 3WJ-IF. Images were acquired on a large-FOV mesoscope system with NA = 0.5, providing a lower photon-collection efficiency imaging condition. Corresponding magnified views are shown. (**D– E**) Quantification of fluorescence signals showing increased sensitivity of 3WJ-IF compared to Conv IF, including mean signal per cell for α-tubulin (D) and mean intensity per TNFα-positive object in inflamed mouse lung tissue (E). Data are presented as mean ± SD. Statistical significance was determined using a two-tailed Welch’s t-test (****P < 0.0001). (**F**) Multiplexed 3WJ-IF whole-slide imaging of an inflamed mouse lung tissue, showing simultaneous detection of TNFα and RNAPII in FFPE tissue sections. For all image panels, identical display settings were used within each experimental comparison to enable direct comparison across conditions; corresponding intensity scales and scale bars are shown in the images.

We first evaluated whether 3WJ-IF improves protein detection sensitivity in cultured cells under low-NA imaging conditions. For α-tubulin staining, 3WJ-IF produced stronger and more clearly defined cytoskeletal signals than conventional immunofluorescence (Conv IF) (**Fig. 5B**) using matched acquisition settings. Quantitative analysis confirmed a significant increase in mean signal intensity per cell, with 3WJ-IF producing more than a three-fold enhancement over Conv IF (**Fig. 5D**). We next tested 3WJ-IF in formalin-fixed, paraffin-embedded (FFPE) tissue sections of the inflamed mouse lung using TNFα as a low-signal target. Under the same large-FOV imaging conditions, TNFα signals were barely detectable by Conv IF (**Fig. 5C**), whereas 3WJ-IF generated markedly stronger staining, corresponding to an approximately 3.6-fold increase in signal intensity (**Fig. 5E**). These results demonstrate that 3WJ-IF enhances signal intensity and detectability in both cultured cells and FFPE tissue specimens, including large-FOV imaging conditions where lower photon collection can limit detection of weak targets.

To compare protein labeling with an alternative split-probe architecture, we evaluated 3WJ-IF and circular split-probe IF (CIR-IF) for EZH2 detection using the same 16× amplification and preassembly procedure. Under these matched amplification conditions, 3WJ-IF generated higher fluorescence signals than CIR-IF (**Fig. S15**), consistent with the signal enhancement observed in the RNA-FISH and DNA origami comparisons.

To determine whether the amplified signal depends on the presence of both split probes, we evaluated unpaired control configurations. In the unpaired negative controls (Neg5 and Neg3), only one component of the 3WJ probe pair was present, preventing complete assembly of the amplifier-docking interface (**Fig. S16**). Compared with the fully matched 3WJ configuration, these unpaired controls produced minimal signal (**Figs. S16C-D**), supporting suppression of amplified labeling from isolated probe components. In contrast, robust nuclear RNAPII signals (red) were observed only under fully matched conditions, while negligible signals were detected in negative controls. The observed signal patterns were consistent with the expected nuclear localization of RNAPII within DAPI-defined nuclear regions (**Fig. S16C)**.

We then evaluated whether 3WJ-IF supports multiplexed protein detection in FFPE tissue using the same large-FOV imaging system. Two-color 3WJ-IF was performed for whole-slide imaging of TNFα (magenta) and RNAPII (green) (**Fig. 5F**) in inflamed mouse lung tissue. TNFα staining was observed in airway-associated regions, while RNAPII showed predominantly nuclear localization, consistent with the expected spatial distributions of these markers. Magnified views of the dashed region further reveal distinct staining patterns, with TNFα enriched in airway-associated regions and RNAPII localized with DAPI-stained nuclei. Separate visualization of individual channels alongside the merged image illustrates the distinct distributions of the two markers within the same tissue section. An additional example of two-color 3WJ-IF detection of TNFα and αSMA together with DAPI is shown in **Fig. S17**. Together, these results establish 3WJ-IF as a programmable oligonucleotide-based amplification strategy for in-situ protein imaging, providing enhanced signal generation in cells and tissues, low background in unpaired probe controls and multiplexed detection in FFPE tissue samples. Signal enhancement under NA = 0.5 ultra-large-FOV imaging conditions supports the utility of 3WJ-IF for multiscale imaging, where sensitive molecular detection must be maintained across large tissue areas.

### 3WJ amplification enables sensitive RNA-protein co-detection for super-resolution imaging

We next investigated whether 3WJ-based amplification enables robust simultaneous detection of RNA and protein targets across imaging modalities (**Fig. 6**). RNA-protein co-detection is technically challenging because RNA-FISH typically involves formamide-containing hybridization buffers, increased temperatures, and stringent washes, which can weaken antibody-based immunofluorescence signals. This signal loss can limit the detection of low-abundance or weakly labeled proteins in combined FISH-IF workflow. Because 3WJ-IF provides oligonucleotide-based signal amplification, we reasoned that it could improve protein detection after RNA-FISH processing.

**Fig. 6.**
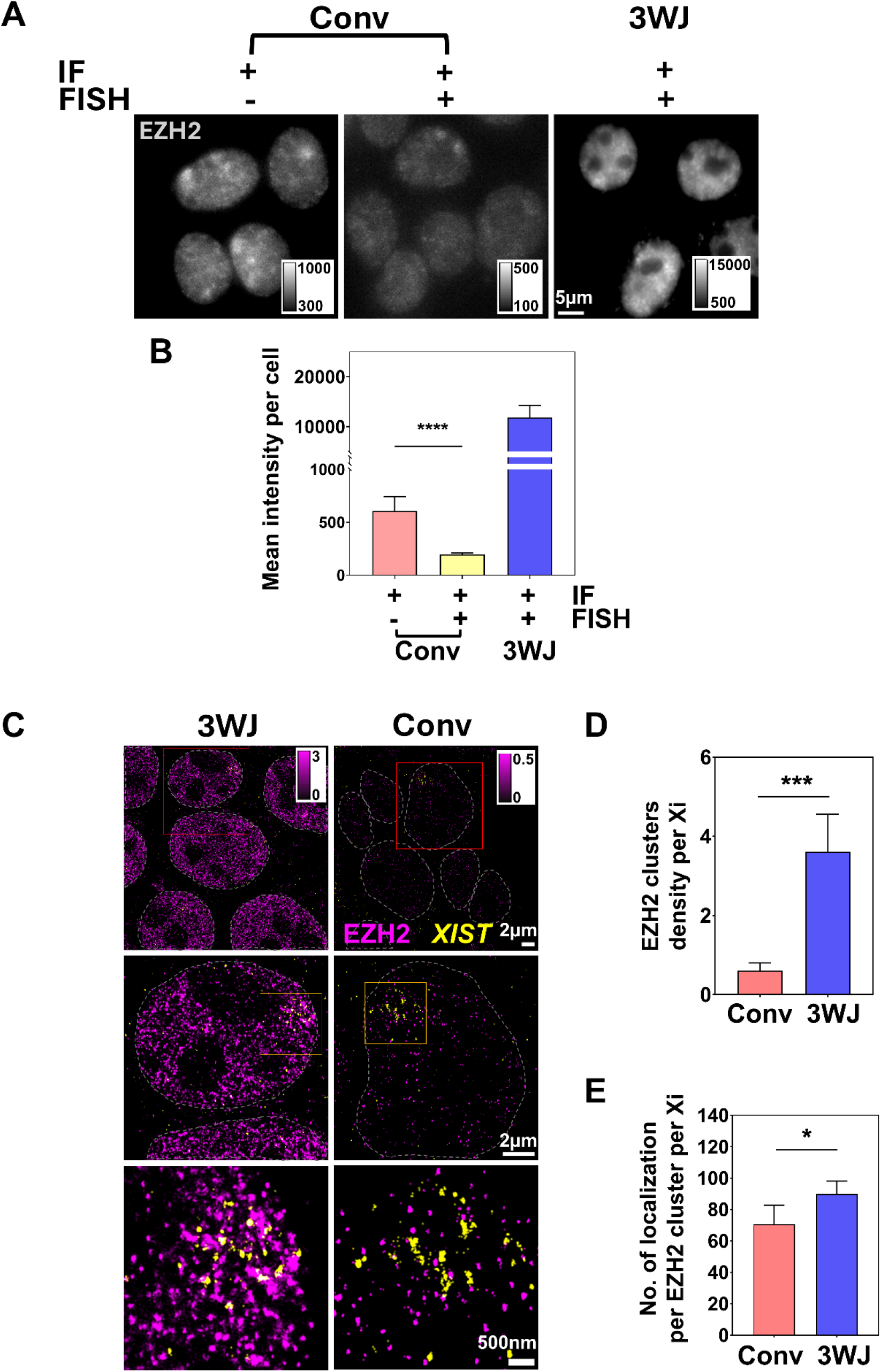
3WJ amplification enables RNA-protein co-detection across widefield and super-resolution imaging. (**A**) Representative widefield (WF) images showing EZH2 immunofluorescence (IF) signal with or without RNA-FISH, comparing conventional IF (Conv) and 3WJ-IF. (**B**) Quantification of mean EZH2 signal intensity per cell under the indicated IF and FISH conditions. Data are presented as mean ± SD. Statistical significance was assessed using Welch’s ANOVA followed by Dunnett’s T3 multiple comparisons test (****P < 0.0001). (**C**) Super-resolution images showing co-detection of EZH2 protein (magenta) and *XIST* RNA (yellow) using 3WJ-FISH and 3WJ-IF, compared to conventional conditions in cultured HEK 293T cells. Insets highlight regions of interest, with corresponding magnified views. (**D–E**) Quantification of EZH2 organization within *XIST*-associated inactive X chromosome regions. Plots show EZH2 cluster density (**D**) and the number of EZH2 localizations within inactive X chromosome (Xi) regions (**E**). Data are presented as mean ± SD. Statistical significance was determined using two-tailed Welch’s t-tests (*P = 0.0100 for E; ***P < 0.0001 for D). All comparisons were performed under matched experimental conditions, including probe concentrations, antibody dilution, incubation procedures, and imaging settings. Display ranges were adjusted independently where needed to visualize both weak and strong signals, with intensity scales and scale bars indicated in the images.

**Figures 6A-B** show that conventional IF produced relatively weak EZH2 signals under the tested conditions, even without RNA-FISH processing. When conventional IF was combined with RNA-FISH, the EZH2 signal was further reduced. In contrast, 3WJ-IF significantly enhanced EZH2 protein signal intensity under the same RNA-protein co-detection conditions. Enhanced protein detection was also observed for RNAPII during co-detection with *XIST* lncRNAs (**Fig. S18**). These results demonstrate that 3WJ-IF improves protein detection in RNA-protein co-detection workflow, especially for targets with weak IF signals that are further compromised by FISH processing.

We next evaluated whether 3WJ-based RNA-protein co-detection is compatible with super-resolution imaging. As shown in **Fig. 6C**, conventional IF combined with RNA-FISH produced insufficient EZH2 signal for reliable STORM reconstruction. In contrast, 3WJ-enabled co-detection of EZH2 protein (magenta) and *XIST* lncRNA (yellow) produced sufficient localization events to reconstruct their nanoscale patterns. Quantitative analysis revealed a significant increase in detected EZH2 cluster density and the number of localizations under 3WJ conditions compared with conventional IF-based co-detection (**Fig. 6D–E**), enabling improved visualization of protein organization relative to *XIST* RNA domains. The super-resolution images revealed that EZH2 signal was enriched within *XIST*-associated inactive X chromosome (Xi) regions (**Fig. 6C**), while RNAPII clusters were largely excluded from *XIST*-associated domains (**Fig. 1D(iv); Fig. S19**). These complementary spatial patterns are consistent with the transcriptionally repressive nature of Xi. The enrichment of EZH2 within *XIST*-associated regions and the exclusion of RNAPII from these domains reveal complementary spatial relationships, consistent with their known functional roles in transcriptional repression and active transcription, respectively. These observations agree with previous reports (*32, 34–36*) and highlight the ability of 3WJ-enabled co-detection to visualize nanoscale RNA-protein organization.

Together, our results demonstrate that 3WJ-based amplification supports simultaneous RNA and protein detection. By combining 3WJ-FISH and 3WJ-IF, the platform supports sensitive, specific, and multiscale analysis of RNA–protein organization, from diffraction-limited imaging to super-resolution localization within cells.

## DISCUSSION

In this study, we developed a compact 3WJ-based oligonucleotide amplification platform for multiscale imaging of RNA and protein targets, enabling target-specific signal amplification while preserving nanoscale spatial fidelity for super-resolution imaging. Although super-resolution microscopy can resolve nanoscale structures, reliable reconstruction often requires sufficient photons and localization events from the target. This becomes challenging for weak or low-abundance targets and RNA-protein co-detection workflows in which FISH processing can reduce immunofluorescence signals. Conventional amplification can improve detectability, but excessive probe size or signal spreading can compromise nanoscale structural information. The 3WJ platform addresses this trade-off by coupling conditional split-probe assembly with a compact 3WJ junction architecture and a pre-assembled amplifier complex, enabling practical signal enhancement with adjustable amplification levels for diffraction-limited and STORM imaging without excessive expansion of the effective probe size. The same amplification principle also improves detection of weak protein targets under large-FOV imaging conditions, where low-NA optics reduce photon collection efficiency. Using DNA origami standards, cultured cells, FFPE tissues and RNA–protein co-detection, we show that this design supports sensitive detection and STORM-based nanoscale structural imaging.

A key design principle of the 3WJ platform is the coupling of paired target recognition, junction assembly and amplifier recruitment. This coupling supports efficient signal generation while limiting amplification from incomplete probe configurations, as demonstrated by the low background in a single unpaired probe and those lacking one or both TRS domains, even when the complementary 10-nt JFS and 9-nt HBS domains are retained. These findings suggest that binding to adjacent target sites brings the probes together and helps stabilize the amplifier-docking assembly, making target recognition an integral part of amplification control.

The local junction configuration contributes to the efficiency of this coupled assembly. The 3WJ design with a 10-nt complementary JFS provides local base pairing and stability to support junction formation and amplifier recruitment, while maintaining a compact probe geometry compatible with nanoscale imaging. Reducing the number of complementary base pairs in the JFS domains, as in the 4C and 0C designs, weakened fluorescence signal and compromised the reconstruction of the nanoscale DNA origami pattern, consistent with fewer detected single molecule localization events. Paired recognition has precedents in existing split-probe methods, such as double Z and circular designs (*25*), while π-FISH also incorporates inter-probe base pairing (*21*). The contribution of the 3WJ design lies in integrating junction formation with short, paired amplifier-binding domains and preassembled amplification. The advantage of this integration is also supported by stronger signals in RNA and protein labeling, including higher fluorescence intensity per RNA spot in *NEAT1* than circular and double-Z designs and enhanced EZH2 detection by 3WJ-IF compared with circular probe conjugated IF. Together, these findings support the junction configuration as an important design parameter for translating target recognition into efficient amplified signal generation within a compact labeling architecture.

The pre-assembled amplification strategy also simplifies delivery compared with sequential branched amplification approaches. Many branched amplification methods rely on sequential hybridization of oligonucleotides containing repetitive domains, which can require extended incubation, denaturation, or carefully optimized assembly conditions (*19, 21*). These stepwise processes may increase variability and expose long single-stranded regions that contribute to nonspecific background. In contrast, the 3WJ platform preassembles the primary-secondary amplifier complex through controlled annealing prior to target binding, conceptually similar to pre-folding a DNA nanostructure before use. This separates amplifier assembly from target labeling and reduces the number of on-sample hybridization steps. LNA-modified binding domains are intended to stabilize the primary-secondary amplifier complex, consistent with the higher signals observed for LNA-containing amplifiers. Together, junction-dependent docking and amplifier preassembly provide a modular approach to improving signal generation while simplifying the labeling workflow.

Amplification can also be adjusted to balance detectability and spatial compactness, as illustrated by the increased signal and apparent labeling size in the DNA origami experiments. This flexibility allows amplification strength to be selected according to the imaging goal, from weak target detection under low-NA optics to super-resolution nanoscale imaging. This compact amplification strategy also addresses a practical barrier to super-resolution imaging of RNA-protein organization. RNA–protein co-detection is challenging because RNA-FISH processing can weaken conventional immunofluorescence signals, which are often substantially reduced following RNA-FISH processing. By combining 3WJ-FISH and 3WJ-IF, we obtained sufficient signal to analyze nanoscale spatial relationships within *XIST*-associated inactive X chromosome regions. The observed EZH2 enrichment and exclusion of RNAPII from *XIST*-associated domains are consistent with the transcriptionally repressive state of Xi (*34–36*).

Beyond overcoming weak protein signals, compact amplification may benefit RNA targets with limited accessible probe-binding sites and the imaging of spatially confined RNA–protein assemblies. The same design is particularly relevant for imaging short or spatially confined RNA species, such as enhancer RNAs (eRNAs) and nascent transcripts, where only a limited number of probe-binding sites may be available. For example, the MYC-bound eRNA MERG1 is ∼500 nt in length (*37*), requiring nearly the entire transcript sequence to design 11 targetable 3WJ probe pairs. Such targets could be challenging for conventional smFISH approaches, which often require at least tens of individual probes to generate sufficient signal intensity. In prior super-resolution imaging studies of *MERG1* eRNA together with MYC protein and nascent *GREB1* transcripts, compact probe size was important for preserving spatial resolution and enabling multiplexed imaging within crowded nuclear environments (*37*). These considerations highlight the potential utility of compact, conditional amplification strategies for studying spatially organized regulatory interactions at the nanoscale.

Beyond super-resolution imaging, the 3WJ amplification strategy also supports sensitive protein detection at the tissue scale. In large-FOV high-throughput imaging, lower-NA optics reduce photon collection efficiency and make weak protein signals more difficult to detect. The improved detection of weak protein targets in FFPE tissue sections with 3WJ-IF illustrates how signal amplification can help address this sensitivity constraint while retaining broad spatial coverage. This extends the same amplification principle from nanoscale RNA–protein localization to large-area tissue mapping.

The current work has several limitations. The current 3WJ platform was evaluated for RNA and protein detection but not for genomic DNA-FISH. Extending the platform to genomic DNA will require validation under DNA denaturation and hybridization conditions, including assessment of target accessibility, probe assembly and background amplification. Multiplexing was demonstrated only for two targets; performance in highly multiplexed or cyclic imaging remains to be established. Such applications will require orthogonal probe and amplifier sets with minimal cross-reactivity and consistent detection across targets.

The modular nature of the 3WJ system provides opportunities for future expansion to address these multiplexing challenges. One potential direction is to adapt the platform for cyclic imaging strategies (*19, 38*) by introducing cleavable elements, such as uracil-containing spacer regions into the amplifiers. This design could allow selective enzymatic destabilization and removal of amplification components after each imaging round, while preserving the target-bound 3WJ probe pairs for subsequent cycles. Efficient amplifier removal and the retention of the target-bound probes would need to be validated over repeated cycles. In parallel, integration of the 3WJ framework with oligonucleotide-conjugated nanobodies or preassembled nanobody-DNA complexes could enable more compact protein labeling. Because nanobodies are smaller than conventional secondary antibodies, this approach may improve tissue accessibility and reduce steric constraints. Together, these future developments could extend the 3WJ platform from sensitive low-plex RNA and protein detection toward scalable, multiplexed molecular imaging across cells and tissues.

Together, these findings establish the 3WJ platform as a compact and programmable amplification strategy for spatially resolved molecular imaging. By coupling paired target recognition and junction assembly with recruitment of preassembled amplifiers, the platform enhances RNA and protein signals while limiting amplification from incomplete probe configuration. Its tunable amplification supports a common framework for imaging across scales, from nanoscale RNA–protein organization within cells to large-FOV tissue mapping.

## MATERIALS AND METHODS

### 3WJ probe design and sequence selection

The 3WJ probe consists of a pair of single-stranded oligonucleotides that assemble into a three-way junction (3WJ) structure when hybridized to adjacent regions of the target. Each probe strand comprises three functional domains: (i) a target-recognition sequence (TRS), (ii) a junction-forming sequence (JFS) that supports 3WJ assembly, and (iii) a half-binding sequence (HBS) that jointly provides an amplifier-docking interface with the HBS of the paired probe (**Fig. 1A-B**).

For RNA detection, candidate 20-nt target-specific sequences were initially generated using Stellaris Probe Designer (*39*) with parameters optimized for specificity (masking level 4, maximum 96 probes, a total of 20-nt probe length for the target sequence). Each candidate region was then extended by 10 nt on both sides to generate a 40-nt target-binding region. The extended sequences were further screened by BLAST against human genome and transcriptome to exclude sequences with potential off-target homology. Selected target-binding regions were divided into two adjacent target-recognition sequences (TRSs), typically 20 nt each, and the corresponding JFS and HBS domains were appended to generate the complete 3WJ probe pair. For DNA origami imaging and 3WJ-based immunofluorescence (3WJ-IF), each probe contained a shorter 10-nt TRS complementary to adjacent regions of the corresponding DNA target handle. Unless otherwise specified, each 3WJ probe strand contained a 10-nt JFS and a 9-nt HBS, with HBS lengths varied from 5 to 20 nt for optimization experiments (**Fig. 4A–F**; **Table S3**).

The HBS and JFS sequences (*14, 40, 41*) were adapted from previously reported orthogonal DNA systems and further selected to minimize predicted secondary structure formation and nonspecific interactions. Unless otherwise specified, HBS sequences were restricted to A, T, and C nucleotides (H = A/T/C) to reduce potential G–C base pairing within the half-binding region. N-HBS designs with unrestricted nucleotide composition were used for comparison (**Fig. S2**). Detailed sequences of the 3WJ probe sets are provided in **Tables S2-S4**.

### Design and assembly of tunable 3WJ amplifiers

Amplifiers were designed as a modular two-stage amplification system (**Fig. S1A**). Upon target-dependent 3WJ assembly, the two half-binding sequences (HBSs) are brought together to form a paired docking interface for primary amplifier recruitment. The primary (1°) amplifier is a single-stranded oligonucleotide that serves as the first amplification backbone. It contains a central 3WJ docking domain that hybridizes to the docking interface formed by the paired HBSs, flanked by multiple secondary (2°) amplifier-binding domains (**Fig. S1B–C**).

The secondary (2°) amplifier was designed as a second amplification backbone containing a central primary-amplifier binding domain that hybridizes to the corresponding binding domains on the primary (1°) amplifier, as well as multiple readout-binding domains for fluorescent readout probe hybridization. Spacer sequences were incorporated between adjacent domains to reduce steric hindrance and limit undesired secondary structure formation. The 16× configuration used 100-nt primary and secondary amplifiers, whereas the 8× configuration used a 58-nt primary amplifier with the same 100-nt secondary amplifier. In the 1× configuration, a fluorescent readout oligonucleotide hybridized directly to the paired HBS docking interface without additional amplification layers. Two orthogonal amplifier/readout systems (Sets 1 and 2) were designed with variants for different amplification configurations and experimental applications. Fluorescent readout probes were labeled at either the 5′ end or both the 5′ and 3′ ends, depending on the application. Detailed sequences, lengths, modifications, and experimental applications of all amplifier and readout variants are provided in **Table S1**.

The domains involved in primary-secondary amplifier binding were constructed using locked nucleic acid (LNA)/DNA mixmers. Each of these binding domains contained five (Set 1) or four (Set 2) LNA-modified nucleotides. LNA-modified nucleotides were incorporated into interacting binding domains and arranged in complementary pairs between the 1° and 2° amplifiers to stabilize inter-amplifier hybridization under hybridization conditions. At the sequence level, the 1° amplifier is designed to hybridize efficiently to the 3WJ probe when the two HBSs are correctly assembled on the target to form the complete docking interface (**Fig. S1C**). Sequences of the 1° and 2° amplifiers are provided in **Table S1**.

To preassemble the amplifier complex before target labeling, 1° and 2° amplifiers were mixed at a 1:4 molar ratio in folding buffer consisting of 1× TAE supplemented with 12.5 mM MgCl_2_. The mixture was subjected to thermal annealing using a controlled cooling ramp from 60°C to 4°C (3 min 12 s per °C). This pre-assembly step allows the 2° amplifiers to hybridize to the 1° amplifier backbone in solution before recruitment to the target-bound 3WJ probe (**Fig. S1B**). Pre-assembly reduces the number of hybridization steps required at the target site and limits exposed single-stranded regions within the amplifier complex. Assembled amplifiers were stored at −20°C until use.

### Oligonucleotide-antibody conjugation

Secondary antibodies (**Table S10**) were buffer-exchanged and concentrated using centrifugal filters (50 kDa MWCO) to remove preservatives and adjust antibody concentrations. Antibody concentrations were determined spectrophotometrically (Nanodrop 2000, ThermoFisher) and adjusted to 2 mg/ml. Purified antibodies were reacted with DBCO-sulfo-NHS ester crosslinkers (10 mg/ml in DMF), using a 10-fold molar excess of crosslinker relative to antibody, for 1.5–2 h at room temperature with gentle mixing, protected from light. Excess crosslinker was removed using desalting columns (7 kDa MWCO).

Azide-modified oligonucleotides (**Table S9**) were then added to the DBCO-modified antibodies at a 3:1 molar ratio of oligonucleotide to antibody and incubated overnight at 4°C to allow copper-free click conjugation. Unreacted oligonucleotides were removed by repeated filtration using 50 kDa MWCO centrifugal filters. The resulting oligonucleotide–antibody conjugates were further purified by ion-exchange high-performance liquid chromatography (IEX-HPLC) available at University of Pittsburgh Small Molecule Biomarker Core to separate unconjugated antibodies, free oligonucleotides, and conjugated species (**Fig. S20**). Purified conjugates were stored at −20°C in storage buffer containing 50% glycerol, 100 µg/ml BSA, and 4 mM EDTA (pH 8.0).

### Fluorophore-antibody conjugation

Secondary antibodies (**Table S10**) were conjugated to fluorophores using a standard NHS-ester labeling protocol (Thermo Fisher Scientific). Briefly, antibodies in 1× PBS were adjusted to 0.1 M sodium bicarbonate (NaHCO₃, pH 8.3) and reacted with fluorophore NHS esters dissolved in DMSO at a dye-to-antibody molar ratio of approximately 5:1. The reaction was incubated for 30 min at room temperature (RT), protected from light. Excess unreacted fluorophores were removed by size-exclusion chromatography, and the resulting conjugates were collected in PBS containing 0.05% sodium azide and 1 mg/ml BSA. For long-term storage, purified conjugates were supplemented with 50% glycerol, aliquoted, and stored at −20°C. The degree of labeling (DOL) was determined spectrophotometrically using a NanoDrop 2000 spectrophotometer (Thermo Fisher Scientific) according to the manufacturer’s instructions.

### Cell and tissue sample preparation

HCT116, HeLa, MCF7, U2OS, and HEK293T cells were cultured in Dulbecco’s modified Eagle’s medium supplemented with 10% fetal bovine serum (Gibco, Invitrogen) and 100 U/ml penicillin–streptomycin (Gibco, 15140148). Cells were maintained at 37°C in a humidified incubator containing 5% CO_2_ until the desired confluency was reached. Cells were fixed with 4% (v/v) paraformaldehyde for 15 min at room temperature (RT) and washed with 1× PBS. For RNA-FISH experiments, fixed cells were stored in 70% (v/v) ethanol at −20°C for at least 12 h prior to permeabilization. For immunofluorescence-based applications (including 3WJ-IF and RNA–protein co-staining), cells were processed immediately after fixation without ethanol storage. Subsequent permeabilization and staining steps were performed as described in the 3WJ-FISH and 3WJ-IF staining section.

Formalin-fixed paraffin-embedded (FFPE) tissue blocks of an inflamed mouse lung were sectioned at 3–5 µm thickness and mounted on gelatin-coated coverslips. Tissue sections were baked at 60°C for 1 h, followed by deparaffinization in xylene and rehydration through a graded ethanol series (100%, 95%, 70%, and 50%). Heat-induced antigen retrieval was then performed in citrate buffer (pH 6.0). To reduce tissue autofluorescence, sections were treated with quenching solution (4.5% H_2_O_2_ and 24 mM NaOH in 1× PBS) under LED illumination for 30 min, repeated 2–3 times as needed. Sections were then rinsed in 1× PBS and stored at 4°C until use.

### Origami preparation

DNA origami structures were designed based on a previously reported 6-helix bundle (6HB) architecture (*29*). Backbone, extension, and biotinylated staple strands were purchased from IDT. Origami structures were assembled in a one-pot reaction in 1× TE buffer supplemented with 12.5 mM MgCl_2_, containing 10 nM M13mp18 single-stranded DNA scaffold (New England Biolabs, N4040S) and 250 nM staple strands, including extension staples carrying H3 target sequences (**Table S7**). The mixture was heated to 80°C for 5 min, held at 65°C for 15 min, and then gradually annealed from 60°C to 4°C at a rate of 3 min 12 s per °C using a thermocycler (Bio-Rad T100). Assembled DNA origami structures were purified using centrifugal filters (50 kDa MWCO, Amicon Ultra 0.5 ml, Merck) to remove excess staple strands. Samples were centrifuged three times at 10,000 ×g for 5 min in FoB5 buffer containing 5 mM Tris base, 1 mM EDTA, 5 mM MgCl_2_, and 5 mM NaCl. Purified origami structures were collected and stored at −20°C.

For imaging, glass coverslips (Fisher Scientific, No. 1.5, 22 × 22 mm, 12541B) were rinsed with 100% ethanol and air-dried prior to chamber assembly. Sample chambers were assembled by mounting SecureSeal™ hybridization chambers (Grace Bio-Labs, SA8R-0.5, 621505) onto the cleaned coverslips. Individual wells were excised from the multi-well gasket and applied prior to use. Chambers were washed three times with ultrapure water, then incubated with biotinylated bovine serum albumin (1 mg/ml in buffer A: 10 mM Tris-HCl, 100 mM NaCl, 0.05% (v/v) Tween 20, pH 8.0) for 5 min. After washing with buffer A, chambers were incubated with streptavidin (0.5 mg/ml in buffer A) for 5 min. After washing with buffer A, chambers were equilibrated with buffer B (5 mM Tris-HCl, 10 mM MgCl₂, 1 mM EDTA, 0.05% (v/v) Tween 20, pH 8.0). Purified DNA origami structures were introduced into the chamber and incubated for 60 min, to allow immobilization through biotin–streptavidin binding. Chambers were then washed with buffer B to remove unbound origami structures. Immobilized DNA origami samples were subsequently used for 3WJ-FISH amplification and imaging.

### 3WJ RNA in situ hybridization (3WJ-FISH)

Fixed and permeabilized cells or tissue sections on coverslips were dehydrated through an ethanol series (70%, 84%, and 100% (v/v)) and air-dried prior to mounting with hybridization chambers. Samples were rehydrated in 1× PBS, permeabilized with 0.5% Triton X-100 for 15 min, treated with 0.1 N HCl for 5 min, and washed with 1x PBS. Samples were blocked in hybridization buffer (5× SSC, 9 mM citric acid, 0.1% (v/v) Tween 20, 0.05 mg/ml heparin, 1× Denhardt’s solution, and 10% dextran sulfate), supplemented with 20% formamide, 10 mM ribonucleoside vanadyl complexes (RVC), 0.5 mg/ml tRNA, and 0.4 µg/ml BSA by incubation at 50°C for 10 min followed by 37°C for 30–60 min. Samples were then washed three times (5 min each) at 37°C with washing buffer (40% formamide in 2× SSC).

3WJ probe sets (50 nM each; **Table S2**) were diluted in hybridization buffer supplemented with 20% formamide and 0.8 U/µl murine RNase inhibitor and applied to the samples. Samples were incubated at 50°C for 20 min for denaturation, followed by hybridization at 37°C overnight. After hybridization, samples were washed three times (5 min each) at 37°C with washing buffer (20% formamide in 2× SSC), followed by brief washes in 2× SSC at room temperature (RT). Pre-assembled amplifiers (50 nM; **Table S1**) were applied in hybridization buffer containing 20% formamide and incubated for 3 h at RT. Samples were then washed three times (5 min each) at RT with washing buffer (20% formamide in 2× SSC), followed by washes in 2× SSC at RT. Fluorescently labeled readout probes (20 nM; **Table S1**) were applied in hybridization buffer supplemented with 20% formamide and RNase inhibitor and incubated for 30–60 min at RT. DAPI was included where indicated. Samples were washed three times (5 min each) at RT with washing buffer (10% formamide in 2× SSC), followed by washes in 6× SSC, and stored in 6× SSC supplemented with 5 mM RVC before imaging. Imaging was performed in buffer containing 10% (w/v) glucose, 0.56 mg/ml glucose oxidase, 0.17 mg/ml catalase, 10 mM NaCl, and 50 mM Tris (pH 8.0).

### Comparison of sequential and pre-assembled amplification

Sequential and pre-assembled amplification workflows were compared using UBC-targeting 3WJ probes (**Fig. 2D–G**). For sequential amplification, LNA-free primary and secondary amplifiers (**Table S1**) were applied in two successive hybridization steps without prior assembly. Each amplifier was hybridized at 50 nM for 3 h at room temperature, using the same hybridization buffer and washing conditions described for 3WJ-FISH. For pre-assembled amplification, LNA-containing primary and secondary amplifiers were annealed as described above before application and hybridized under the same conditions in a single step. Subsequent readout probe hybridization, washing, and imaging were performed as described for 3WJ-FISH.

To evaluate the effect of LNA incorporation on pre-assembled amplification, RNAPII was detected by 3WJ-IF using LNA-containing or LNA-free primary and secondary amplifiers (**Fig. S3**; **Table S1**). Both amplifier complexes were pre-assembled under identical annealing conditions and applied using the same 3WJ-IF workflow. Fluorescence intensity was quantified to compare signal amplification between the two conditions.

### HBS-length optimization

To evaluate the effect of HBS length on nonspecific probe retention and signal amplification, UBC-targeting split probes with HBS lengths of 5, 9, 15, and 20 nt were designed (**Table S3**; **Fig. 4A–F**). The 5- and 9-nt HBS designs were evaluated as functional split probes, whereas the 15- and 20-nt designs served as controls for nonspecific probe retention in the absence of productive 3WJ assembly. All experiments used the 8× amplification configuration with a primary amplifier containing an extended HBS-docking domain (P-Amp-1b-HBSopt; **Table S1**). Probe hybridization, signal amplification, and imaging were performed as described for 3WJ-FISH.

### Comparison of H-HBS and N-HBS designs

The effects of HBS nucleotide composition on signal amplification were evaluated by comparing N-HBS (unrestricted nucleotide composition) and H-HBS (restricted to A, T, and C) designs (**Fig. S2**). *GAPDH* RNA in HCT116 cells and RNAPII Ser2P in MCF7 cells were detected using the corresponding 3WJ probe sets (**Tables S2** and **S4)**. N-HBS and H-HBS probes were paired with their respective primary amplifiers (P-Amp-1c-16x and P-Amp-1a-16x; **Table S1**). Probe hybridization, signal amplification, and imaging were performed as described for 3WJ-FISH and 3WJ-IF, respectively.

### Evaluation of 3WJ probe specificity

The specificity of target-dependent 3WJ assembly was evaluated using *NEAT1*-targeting probe pairs and a series of negative controls (**Fig. 4G–H**; **Table S2**). Negative controls included individual 5′ or 3′ probes (Neg5 and Neg3), combinations of a target-binding probe with a corresponding probe lacking the target-recognition sequence (Un5 and Un3), and probe pairs lacking target-recognition sequences on both strands (Un). All conditions were evaluated using 16× signal amplification (**Table S1**), with probe hybridization, amplification, and imaging performed as described for 3WJ-FISH.

### Single-molecule fluorescence in situ hybridization (smFISH)

smFISH was performed as a reference method for comparison with 3WJ-FISH. Sample preparation, including fixation, dehydration, rehydration, permeabilization, HCl treatment, and blocking, was carried out as described for 3WJ-FISH. smFISH probe sets (50 nM; **Table S8**) were diluted in hybridization buffer containing 20% formamide and 0.8 U/µl murine RNase inhibitor and applied to the samples. Samples were incubated at 50°C for 20 min for denaturation, followed by hybridization at 37°C overnight. After hybridization, samples were washed three times (10 min each) at 37°C with washing buffer (10% formamide in 2× SSC), followed by brief washes in 2× SSC at room temperature (RT). Fluorescently labeled readout probes (20 nM; **Table S1**) were applied and incubated at RT as described for 3WJ-FISH. Samples were then washed three times (5 min each) at RT with washing buffer (10% formamide in 2× SSC), followed by washes in 6× SSC, and stored in the dark at 4°C prior to imaging. Imaging was performed as described for 3WJ-FISH.

### Comparison of 3WJ-FISH and conventional smFISH

The detection performance of 3WJ-FISH and conventional smFISH was compared across RNA targets with different expression levels, including *UBC*, *TOP2A*, and *EPIC1* (**Fig. 3A–D** and **Figs. S7–S11).** To ensure comparable target recognition, each smFISH probe was designed using the TRS of one strand from the corresponding 3WJ probe pair, appended with an smFISH readout-binding domain. Thus, both methods targeted the same number of RNA sites, with 3WJ-FISH using two adjacent TRSs per site and smFISH using one of the corresponding TRSs. Probe sequences are provided in **Tables S2** and **S8**. Both methods were performed using their respective hybridization and detection protocols described above. Detection performance was evaluated by comparing mean fluorescence intensity and RNA spot counts per cell where individual spots could be reliably resolved. The relationship between RNA spot counts and RNA-seq expression was analyzed as described under Correlation of RNA spot counts with RNA-seq expression.

### Comparison of probe architecture for RNA detection

Z-FISH, CIR-FISH, and 3WJ-FISH were compared using probe pairs targeting *NEAT1*. All three probe architectures shared identical target recognition sequences (TRS) but differed in their functional domains, forming Z-shaped, circular, and three-way junction structures, respectively (**Fig. S6A**). Probe sequences are provided in **Tables S2**, **S5**, and **S6**. All three methods used the same pre-assembled amplifiers and fluorescent readout probes for 16× signal amplification (**Table S1**). Sample preparation, probe hybridization, washing, signal amplification, and imaging were performed as described for 3WJ-FISH. Detection performance was evaluated by quantifying the number of NEAT1 spots per nucleus and the mean fluorescence intensity of detected spots.

### Multiplexed RNA detection

For multiplexed RNA detection, target-specific 3WJ probe sets corresponding to two orthogonal amplifier systems (Sets 1 and 2; **Tables S1** and **S2**) were used to detect distinct RNA targets in the same sample (**Fig. 1D(*iii*)** and **Fig. S13**). Following probe hybridization, the corresponding pre-assembled amplifiers and fluorescent readout probes were applied using the procedures described for 3WJ-FISH. The two targets were distinguished using spectrally distinct fluorophores (Alexa Fluor 647 and CF568).

### 3WJ-based immunofluorescence staining (3WJ-IF)

Fixed cultured cells were permeabilized with 0.5% (v/v) Triton X-100 in 1× PBS for 15 min and blocked in antibody blocking buffer (3% (w/v) bovine serum albumin (BSA) and 0.02% (v/v) Triton X-100 in 1× PBS) for 1 h at room temperature (RT). Samples were incubated with primary antibodies (**Table S10**) in blocking buffer overnight at 4°C, followed by three washes (5 min each) with antibody washing buffer (0.3% (w/v) BSA and 0.02% (v/v) Triton X-100 in 1× PBS).

Oligonucleotide-conjugated secondary antibodies were applied in blocking buffer and incubated for 2 h at RT, followed by washing as described above. To stabilize antibody labeling before oligonucleotide hybridization, samples were then post-fixed with 4% (w/v) paraformaldehyde (PFA) in 1× PBS for 20 min at RT. To quench residual aldehyde groups, samples were incubated with 100 mM NH_4_Cl in 1× PBS for 20 min at RT, followed by three washes with 1× PBS.

Samples were equilibrated in hybridization buffer and blocked with 0.5 mg/ml tRNA for 1 h. 3WJ probe sets (**Table S4**) were then applied in hybridization buffer supplemented with 20% formamide for 6–12 h at RT. After hybridization, samples were washed three times (5 min each) at RT with washing buffer (20% formamide in 2× SSC), followed by washes in 2× SSC. Subsequent amplifier hybridization, signal development, and imaging were performed using amplifier and readout probes (**Table S1**) as described for 3WJ-FISH.

For FFPE tissue sections, 3WJ-IF was performed following deparaffinization, antigen retrieval, and autofluorescence quenching as described under Cell and tissue sample preparation. Subsequent antibody labeling, 3WJ probe hybridization, signal amplification, and imaging followed the procedures described above.

### Immunofluorescence staining (IF)

Conventional immunofluorescence staining was performed as a reference method for comparison with 3WJ-IF. Permeabilization, blocking, and primary antibody incubation were performed as described for 3WJ-based immunofluorescence staining (3WJ-IF). Fluorophore-conjugated secondary antibodies were used in place of oligonucleotide-conjugated secondary antibodies and incubated under the same conditions. After secondary antibody incubation, samples were washed and post-fixed with 4% (w/v) PFA in 1× PBS for 20 min at room temperature (RT). Residual aldehydes were quenched by incubation with 100 mM NH_4_Cl in 1× PBS for 20 min, followed by washing with PBS. Samples were then imaged using the same imaging conditions as those used for the corresponding 3WJ-IF experiments.

### Circular probe-based immunofluorescence staining (CIR-IF)

CIR-IF was performed using the same staining and hybridization procedures as described for 3WJ-IF, except that the 3WJ probe pair was replaced with the corresponding circular (CIR) probe pair (**Table S4**). The CIR probes were designed to form a circular hybridization architecture rather than a three-way junction, while retaining the same antibody-associated target recognition domains. Both CIR-IF and 3WJ-IF were performed using 16× signal amplification with the same amplifier and readout probes (**Table S1**). Nuclear EZH2 fluorescence intensity was quantified to compare the detection performance of the two probe architectures (**Fig. S15**).

### Co-staining RNAs and proteins by 3WJ probes

For simultaneous detection of RNAs and proteins, fixed cultured cells were processed using the 3WJ-based immunofluorescence (3WJ-IF) protocol, including permeabilization, blocking, primary antibody incubation, oligonucleotide-conjugated secondary antibody incubation, post-fixation, and equilibration in hybridization buffer.

To preserve RNA integrity during the immunofluorescence steps, 5 mM Ribonucleoside Vanadyl Complexes (RVC) were included in all buffers used during the 3WJ-IF procedure. After antibody labeling and post-fixation, samples were subjected to the 3WJ-FISH protocol to detect target RNAs and the oligonucleotide-conjugated secondary antibodies. This included 3WJ probe hybridization, amplifier hybridization, fluorescent readout-probe hybridization, washing, and imaging steps as described above.

### 3WJ-FISH-based signal amplification on DNA origami

Application of the 3WJ-FISH protocol to DNA origami samples followed the same general workflow as described for cells and tissue samples, with modifications to preserve origami integrity. DNA origami structures, assembled from single-stranded DNA (ssDNA) scaffolds and staple strands through programmed base pairing, are sensitive to denaturing conditions. Accordingly, denaturing reagents and elevated temperatures were avoided. All hybridization and washing buffers were prepared without formamide, and all incubation steps were performed at room temperature (RT). RNase inhibitors were not included in the buffers.

After surface immobilization of DNA origami structures, 3WJ probe hybridization, amplifier hybridization, washing, and imaging steps were performed as described above, except under non-denaturing room-temperature conditions. Fluorescent microspheres (0.1 μm diameter, F8803, Invitrogen), diluted 1:10,000 in 2× SSC, were added to the chamber and incubated for 30 min at RT as fiducial markers for drift correction during super-resolution imaging. The bead solution was removed and replaced with 2× SSC prior to imaging. Samples were stored in 2× SSC and exchanged into imaging buffer immediately before STORM imaging.

### Comparison of split-probe architectures on DNA origami

Five split-probe architectures (3WJ, 4C, π, 0C, and CIR) were compared using DNA origami structures carrying identical target-recognition sequences (**Table S4**; **Fig. 2A**; **Fig. S4A**). All architectures were evaluated using 16× signal amplification (**Table S1**). Probe hybridization, signal amplification, and imaging were performed as described above. Detection performance was evaluated by quantifying mean widefield fluorescence intensity and the number of STORM localizations per origami structure (**Fig. 2B-C**; **Fig. S4B–D**).

### Image acquisition and processing

Fluorescence imaging was performed using two custom-built microscopy platforms, with acquisition conditions summarized in **Table S11**. Super-resolution (STORM) and diffraction-limited widefield imaging were conducted on a custom-built STORM microscope based on an Olympus IX71 inverted microscope equipped with an oil immersion objective (UPLSAPO 100×, NA 1.4; Olympus), as described previously (*42*). Large-FOV mesoscopic widefield imaging was performed on a separate custom-built mesoscope system (*33, 43*).

For each target within a given experiment, imaging parameters including exposure time, gain, and illumination intensity were kept constant across all comparative conditions (e.g., HBS length optimization, specificity assays, and sensitivity analyses) to enable direct and quantitative comparison of fluorescence signals. Imaging settings were adjusted between different targets and fluorophores as needed to accommodate differences in signal intensity and dynamic range. Unless otherwise stated, all comparative imaging experiments within the same dataset were performed using identical sample preparation procedures, probe concentrations, incubation conditions, microscope acquisition settings, and image processing workflows. For visual comparison, images within the same experimental set were displayed using matched intensity scales whenever possible. When signal magnitudes differed substantially between conditions, display ranges were adjusted independently to visualize both weak and strong signals, and the corresponding intensity scales were indicated in each panel.

For STORM imaging, samples were excited using a 642 nm laser (VFL-P-1000-642-OEM3; MPB Communications, Quebec, Canada) under oblique illumination at a power density of ∼2.5 kW·cm⁻². Images were acquired with an exposure time of 20 ms for a total of ∼20,000 frames. Two-color STORM imaging was performed sequentially, with Alexa Fluor 647 imaged first, followed by CF568 using a 561 nm laser (VFL-P-200-560-OEM1; MPB Communications) at ∼0.8 kW·cm⁻². Drift correction was performed every 200 frames (∼4 s) using fluorescent nanospheres excited with a 488 nm laser (DL488-150, CrystaLaser, Reno, NV) as fiducial markers.

High-resolution widefield images were acquired on the STORM microscope (*42*) and large-FOV mesoscopic widefield images were acquired on the Mesoscope, processed using their respective previously published analysis pipelines (*33, 43*), including image reconstruction, flat-field correction, stitching, registration, and segmentation implemented in custom MATLAB-based workflows.

### Quantitative image analysis

RNA spot counting was performed using custom MATLAB scripts. Fluorescence images were background-corrected, and RNA spots were identified as local intensity maxima above manually selected thresholds. Thresholds were kept constant across comparative conditions within the same experiment. The number of RNA spots per cell and spot intensity were quantified within manually defined cell regions of interest (ROIs).

For immunofluorescence intensity analysis, images were processed in ImageJ. Signal regions were defined using Otsu thresholding on fluorescence images. The resulting binary masks were used to generate ROIs, which were applied to the original fluorescence images to calculate mean pixel intensity within each ROI.

For DNA origami quantification, individual 6-helix bundle (6HB) structures were manually selected based on their characteristic three-spot linear pattern, corresponding to the designed arrangement of DNA handles. This selection criterion was used to distinguish properly assembled structures from background signals and aggregates. ROIs corresponding to individual 6HB structures were used to quantify signal intensity and localization features using ImageJ on reconstructed STORM images and corresponding widefield images. Cluster size was obtained using custom MATLAB scripts as described previously (*42*). Acquisition and analysis thresholds were kept constant across comparative conditions within the same experiment to ensure accurate and unbiased comparisons.

For distance validation (**Fig. S5**), distances between adjacent fluorescence spots (d1 and d2) within individual 6HB DNA origami structures were measured from reconstructed STORM images using ImageJ, with pixel distances converted to nanometers based on image calibration. Measurements from 28 structures were compiled in Excel to calculate the mean and standard deviation for each distance. Distance distributions were plotted using GraphPad Prism.

For EZH2–*Xist* co-localization analysis, *Xist*-positive inactive X chromosome (Xi) regions were manually defined as ROIs. Custom MATLAB scripts published in a previous study (*42*) were used to quantify EZH2 cluster density and the number of EZH2 localizations within each Xi ROI from reconstructed STORM images. Quantitative analyses were performed using identical thresholding and analysis parameters across comparative conditions within the same experiment to ensure unbiased signal quantification.

### Correlation of RNA spot counts with RNA-seq expression

Mean RNA spot counts per cell obtained using 3WJ-FISH and conventional smFISH were compared with RNA-seq expression data obtained from the Cancer Cell Line Encyclopedia (CCLE; DepMap Public 22Q1) (*30*). Simple linear regression was performed using GraphPad Prism to evaluate the relationship between RNA spot counts and RNA-seq expression levels across different genes and cell lines (**Fig. 3D** and **Fig. S7**). Single-cell distributions of RNA spot counts and fluorescence intensities obtained by 3WJ-FISH are presented in **Fig. S12**. The corresponding mean RNA spot counts and RNA-seq expression values are summarized in **Table S13**.

### Statistical analysis

Statistical analyses were performed using GraphPad Prism. Details of statistical comparisons, sample sizes (n), mean ± standard deviation (SD), statistical tests, and corresponding P values are summarized in **Table S12**. For comparisons involving more than two groups, Welch’s ANOVA followed by Dunnett’s T3 multiple comparisons test or the Kruskal–Wallis test followed by Dunn’s multiple comparisons test was used, as appropriate. For pairwise comparisons, two-tailed unpaired Welch’s t-tests or Mann–Whitney U tests were used, as appropriate. Parametric or nonparametric tests were selected based on the distribution and characteristics of the data. All statistical analyses were performed using datasets acquired and processed under matched experimental and analysis conditions.

## Reagents and materials

A complete list of reagents and materials used in this study, including suppliers and catalog numbers, is provided in **Table S14**.

## Author contributions

Conceptualization: PN, YL

Methodology: PN, YL

Investigation: PN, XS

Formal analysis: PN, HM, MC, SL, DY

Software: HM

Resources: HM, AS

Visualization: PN

Funding acquisition: YL

Project administration: YL

Supervision: YL

Writing – original draft: PN

Writing – review & editing: PN, YL

## Data, Code, and Materials Availability

All data needed to evaluate the results in this paper are provided in the manuscript, Supplementary Materials, and source data files. This study did not generate new materials. The images and numerical data underlying the main figures are publicly available on Zenodo: https://doi.org/10.5281/zenodo.23792866. Code for RNA spot counting and example RNA-FISH data are available at https://doi.org/10.5281/zenodo.23377747. All oligonucleotide probe and amplifier sequences designed for this study are provided in Supplementary Tables.

