## Supplementary Materials for "A compact three-way junction oligonucleotide probe for widefield and super-resolution imaging of RNAs and proteins"

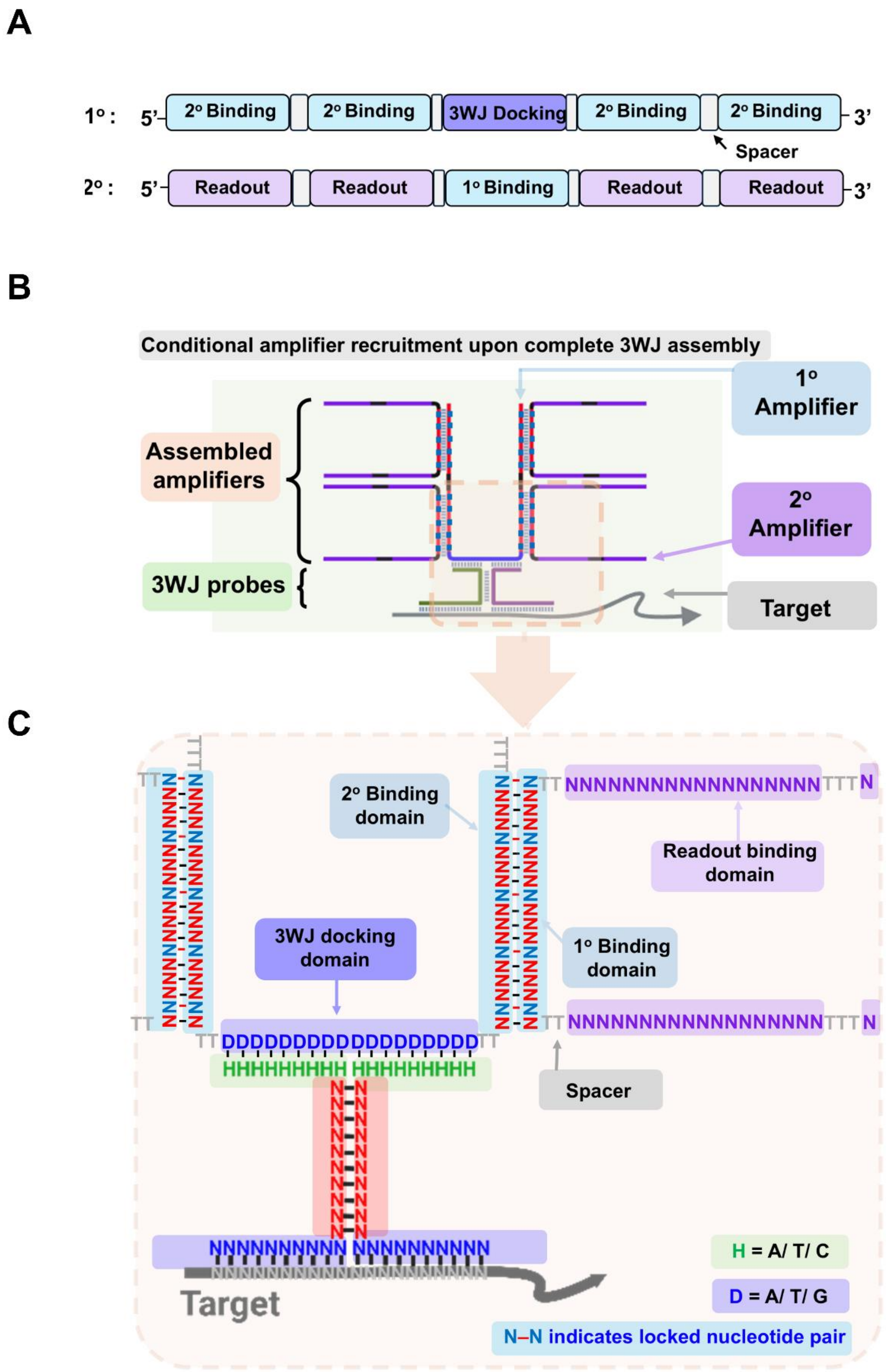
A
1°:
5'
2° Binding
2° Binding
3WJ Docking
2° Binding
2° Binding
3'
Spacer
2°:
5'
Readout
Readout
1° Binding
Readout
Readout
3'
B
Conditional amplifier recruitment upon complete 3WJ assembly
1° Amplifier
Assembled amplifiers
2° Amplifier
3WJ probes
Target
C
2° Binding domain
Readout binding domain
3WJ docking domain
1° Binding domain
Spacer
Target
H = A/ T/ C
D = A/ T/ G
N–N indicates locked nucleotide pair

**Fig. S1. Sequence design and assembly of pre-assembled amplifiers for 3WJ-mediated signal amplification.**

**(A)** Domain organization of the primary (1°) and secondary (2°) amplifiers, each 100 nt in length for 16x amplification. The primary amplifier contains a central 3WJ docking domain flanked by a total of four 2° amplifier-binding domains, while the secondary amplifier contains a central 1° amplifier-binding domain flanked by a total of four readout amplifier-binding domains. Spacer regions are inserted between adjacent domains, with the aim of limiting steric hindrance and undesired secondary structure formation.

(**B**) Schematic of amplifier preassembly. Primary and secondary amplifiers are pre-mixed at a defined stoichiometric ratio and annealed prior to use, allowing complementary amplifier-binding domains to hybridize while leaving the 3WJ docking and readout-binding domains available for subsequent hybridization.

(**C**) Sequence-level representation of the 3WJ-mediated amplification architecture. The 3WJ docking domain of the primary amplifier, composed of D bases (A/T/G), hybridizes with the 9-nt half-binder sequence (HBS; H = A/T/C) domains brought together by the target-bound probe pair. The 1° and 2° amplifier-binding domains contain five (Set 1) or four (Set 2) locked nucleic acid (LNA)-modified nucleotides per domain to stabilize interactions between the primary and secondary amplifiers upon annealing. The secondary amplifier provides four readout binding domains for fluorescent probe hybridization, enabling signal amplification. N–N indicates base pairing between LNA-modified nucleotides.

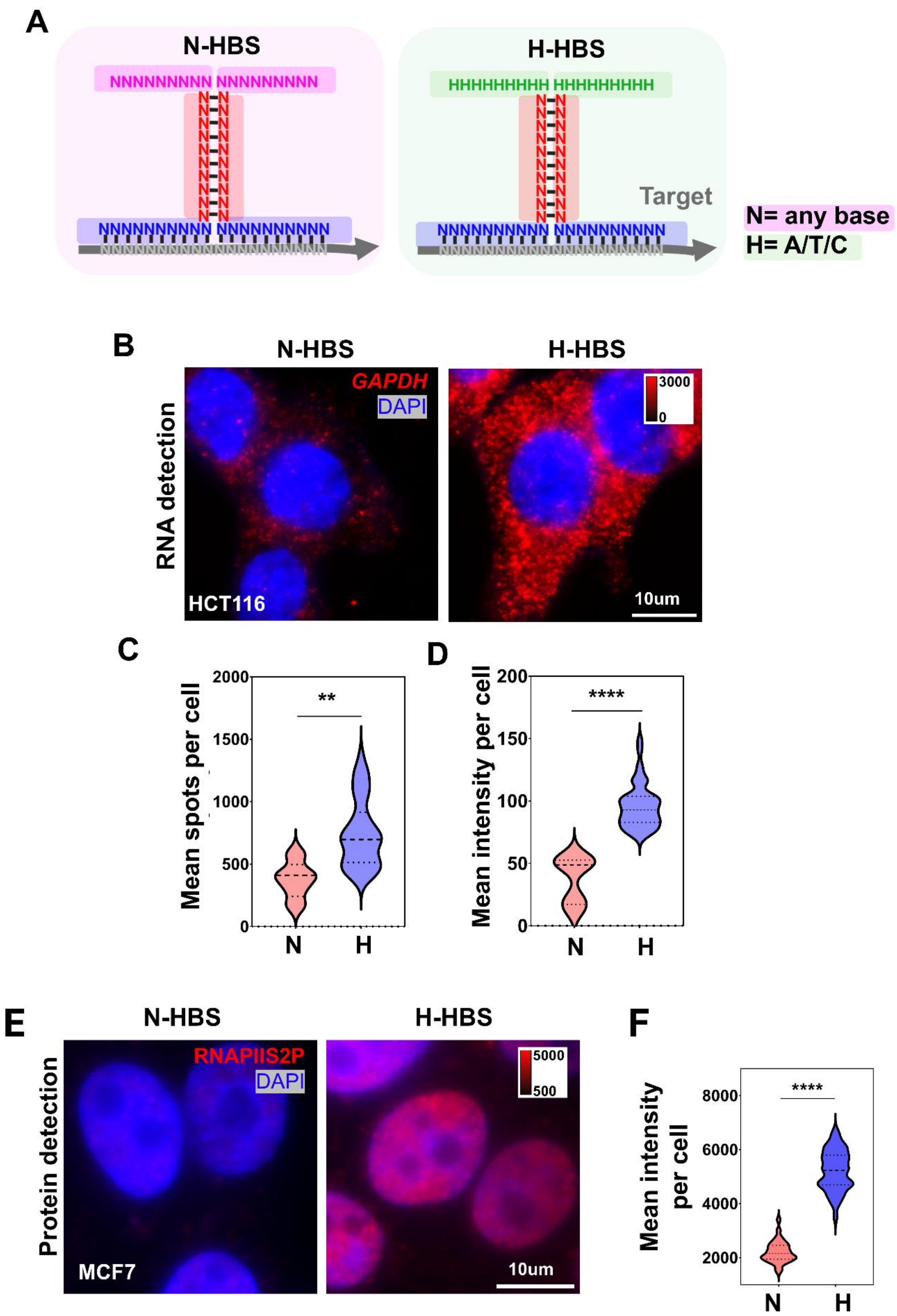
A
N-HBS
NNNNNNNNN NNNNNNNNN
NNNNNNNNNN NNNNNNNNNN
H-HBS
HHHHHHHHH HHHHHHHHH
Target
NNNNNNNNNN NNNNNNNNNN
N= any base
H= A/T/C
B
N-HBS
H-HBS
RNA detection
GAPDH
DAPI
HCT116
3000
0
10um
C
Mean spots per cell
2000
1500
1000
500
0
**
N
H
D
Mean intensity per cell
200
150
100
50
0
****
N
H
E
N-HBS
H-HBS
Protein detection
RNAPIIS2P
DAPI
MCF7
5000
500
10um
F
Mean intensity per cell
8000
6000
4000
2000
****
N
H

**Fig. S2. Optimization of half-binding sequence (HBS) composition to improve amplifier assembly and detection efficiency.**

(**A**) Schematic illustration of HBS design. The N-HBS design uses any base (N = A/T/G/C), whereas the optimized H-HBS design restricts bases to A/T/C to reduce potential G–C base pairing within the half-binding region. Both designs used refolded amplifiers with 16× signal amplification and shared the same secondary amplifier-binding domain and readout probe, differing only in the HBS composition of the primary amplifier.

(**B–D**) RNA detection using N-HBS and H-HBS designs. (B) Representative images of *GAPDH* RNA in HCT116 cells. (C–D) Violin plots showing the distribution of mean spots per cell (C) and mean signal intensity per cell (D). Center lines indicate medians, and dotted lines represent quartiles. Statistical significance was determined using two-tailed Welch's t-tests (**P = 0.0047 for C; **P = 0.0015 for D).

(**E–F**) Protein detection using N-HBS and H-HBS designs. (E) Representative images of RNAPII Ser2P in MCF7 cells. (F) Violin plot showing the distribution of mean signal intensity per cell. Center lines indicate medians, and dotted lines represent quartiles. Statistical significance was determined using a two-tailed Welch's t-test (****$P < 0.0001$).

Intensity scaling and scale bars are consistent within each comparison, and corresponding display ranges are indicated in the images.

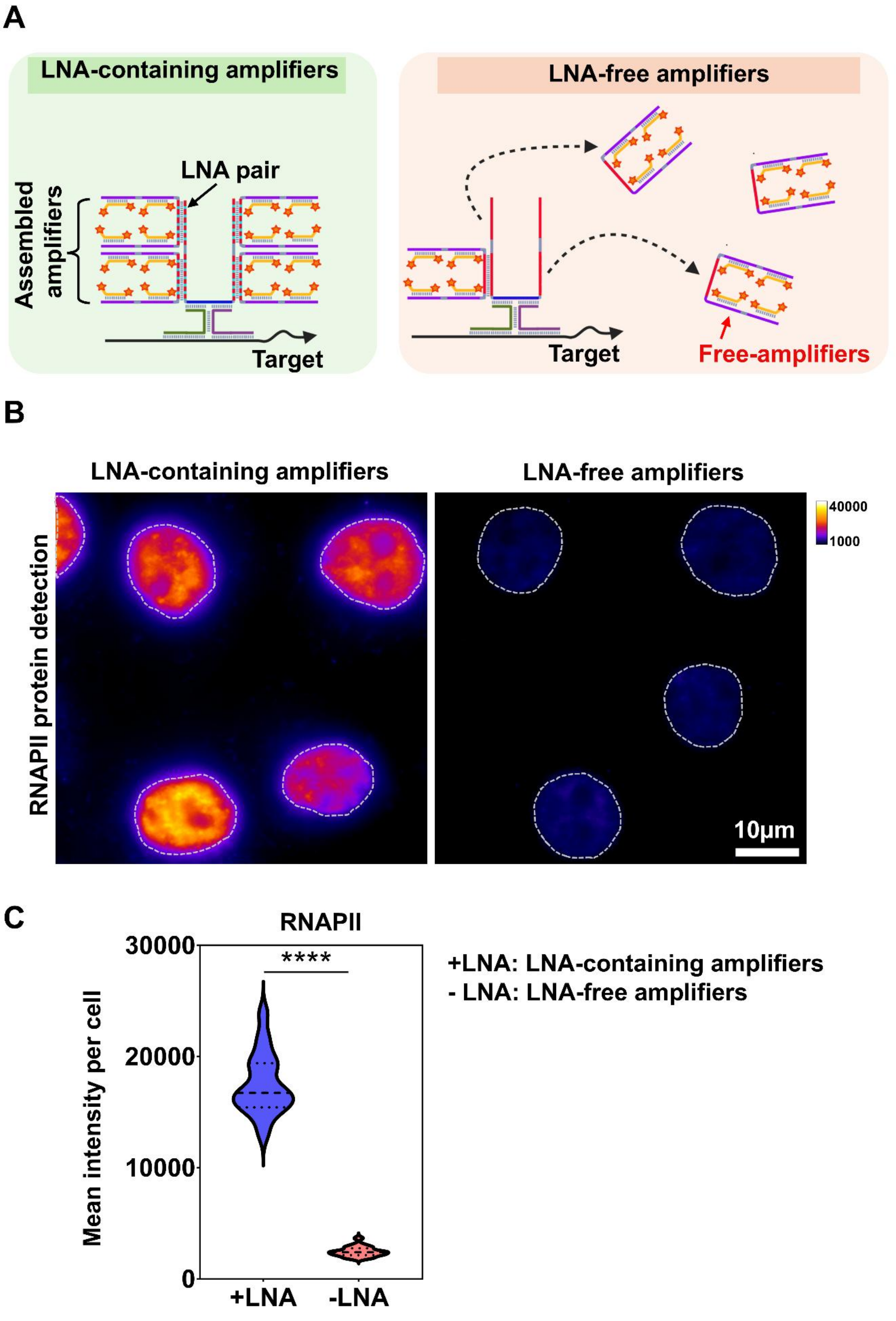
A
LNA-containing amplifiers
LNA pair
Assembled amplifiers
Target
LNA-free amplifiers
Target
Free-amplifiers
B
LNA-containing amplifiers
LNA-free amplifiers
RNAPII protein detection
40000
1000
10μm
C
RNAPII
****
30000
20000
10000
0
Mean intensity per cell
+LNA
-LNA
+LNA: LNA-containing amplifiers
- LNA: LNA-free amplifiers

**Fig. S3. LNA incorporation stabilizes amplifier assembly and enhances signal output.**

(**A**) Proposed model illustrating the role of locked nucleic acid (LNA) incorporation in stabilizing amplifier assembly. LNA-containing amplifiers form stable assembled complexes through enhanced base pairing, whereas LNA-free amplifiers exhibit reduced stability and dissociate into free amplifiers, impairing effective assembly. The LNA-free (−LNA) amplifiers retained the same nucleotide sequences, with all LNA residues replaced by DNA (P-Amp-1a-16x-noLNA and S-Amp-1a-noLNA; Table S1).
(**B**) Representative widefield images of RNA polymerase II (RNAPII) detection using 3WJ-IF with LNA-containing (+LNA) or LNA-free (−LNA) amplifiers. Nuclei are outlined (dashed lines). Scale bar, 10 µm. Intensity scales are indicated in the images.
(**C**) Quantification of mean RNAPII signal intensity per cell under +LNA and −LNA conditions. Data are presented as violin plots with median and interquartile range. Statistical significance was determined using a two-tailed unpaired Welch's t-test; ****$P$ < 0.0001.

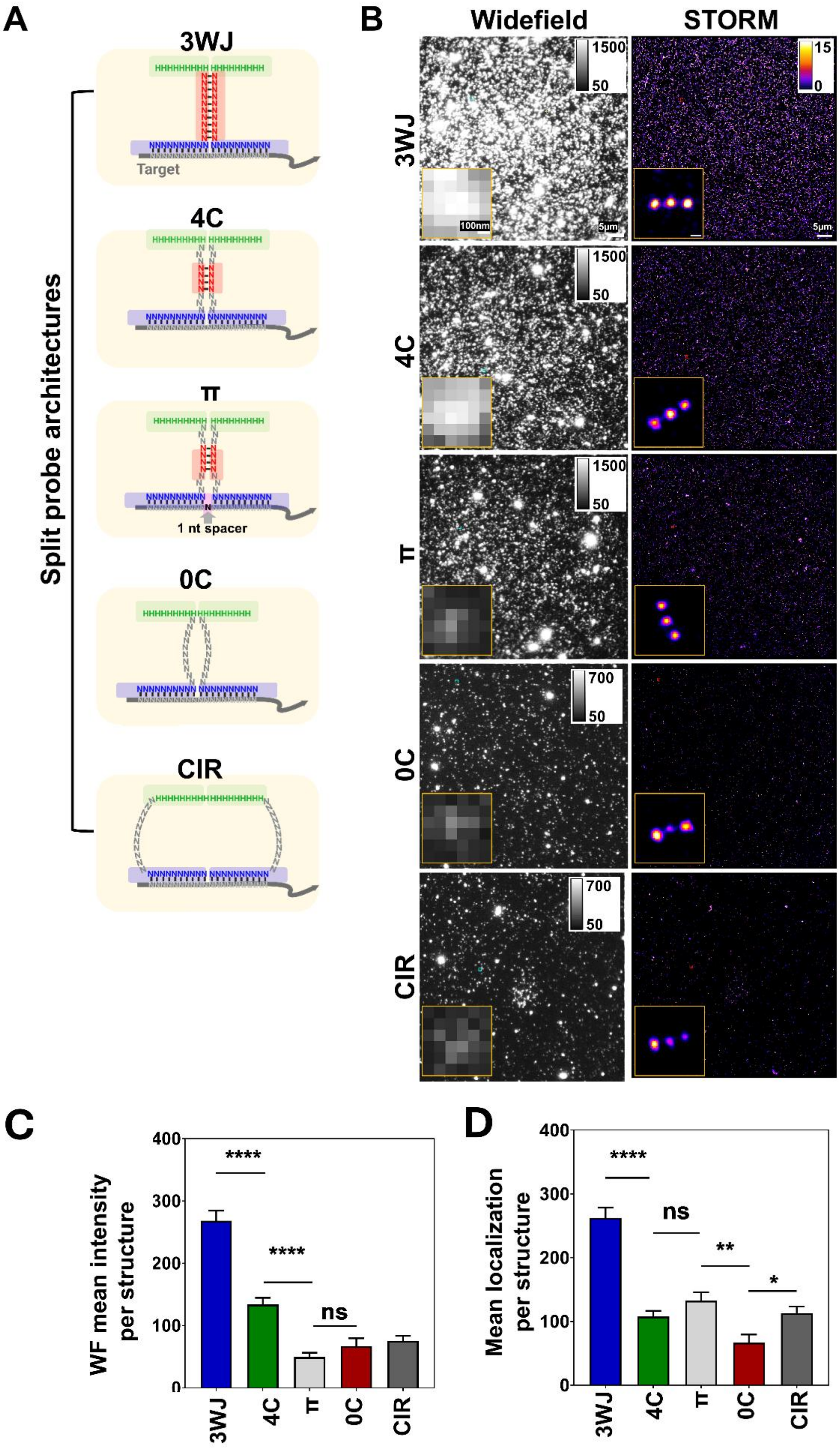
A
Split probe architectures
3WJ
Target
4C
π
1 nt spacer
0C
CIR
B
Widefield
STORM
3WJ
4C
π
0C
CIR
1500
50
700
15
0
100nm
5µm
C
WF mean intensity per structure
400
300
200
100
0
****
ns
3WJ
4C
π
0C
CIR
D
Mean localization per structure
****
ns
**
*

**Fig. S4. Comparison of split probe architectures using DNA origami targets.**

(**A**) Schematic illustration of split probe architectures, including three-way junction (3WJ), four-connector (4C), π-shaped (π), zero-connector (0C), and circular (CIR) designs. The 3WJ design contains a 10-nt complementary JFS, whereas the 4C and 0C variants contain four complementary bases and no designed base pairing in the JFS region, respectively. The π design contains a 1-nucleotide spacer between the two target-recognition regions, whereas the 4C design lacks this spacer.

(**B**) Representative widefield (WF) and super-resolution (STORM) images of DNA origami labeled with each probe design (16x amplification). Full-field images are shown in the main panels, with magnified views of a representative individual DNA origami structure shown in the corresponding insets. Cyan and red boxes indicate the regions magnified in the WF and STORM insets, respectively. WF images were displayed using individually adjusted intensity scales, whereas STORM images were displayed using a common intensity scale. Scale bars, 5 μm (main panels) and 100 nm (insets). Due to the diffraction-limited resolution from WF images, the multiple labeled origami structures resolved by STORM can fall within a single diffraction-limited WF spot.

(**C–D**) Quantification of signal output across probe designs, including mean WF intensity per structure (C) and mean STORM localization count per individually resolved 3-spot origami structure (D). STORM analysis included only structures with a visible three-dot pattern (those with two-dot or one-dot patterns excluded). Higher WF intensity in the 4C condition than in the π condition is consistent with more labeled origami contributing to individual diffraction-limited spots, whereas STORM localization counts per resolved, complete three-dot DNA origami structure were not significantly different between 4C and π-shaped designs. Data are presented as mean ± SD. Statistical significance was assessed using Welch's ANOVA followed by Dunnett's T3 multiple comparisons test (****$P < 0.0001$; **$P < 0.01$; *$P < 0.05$; ns, not significant).

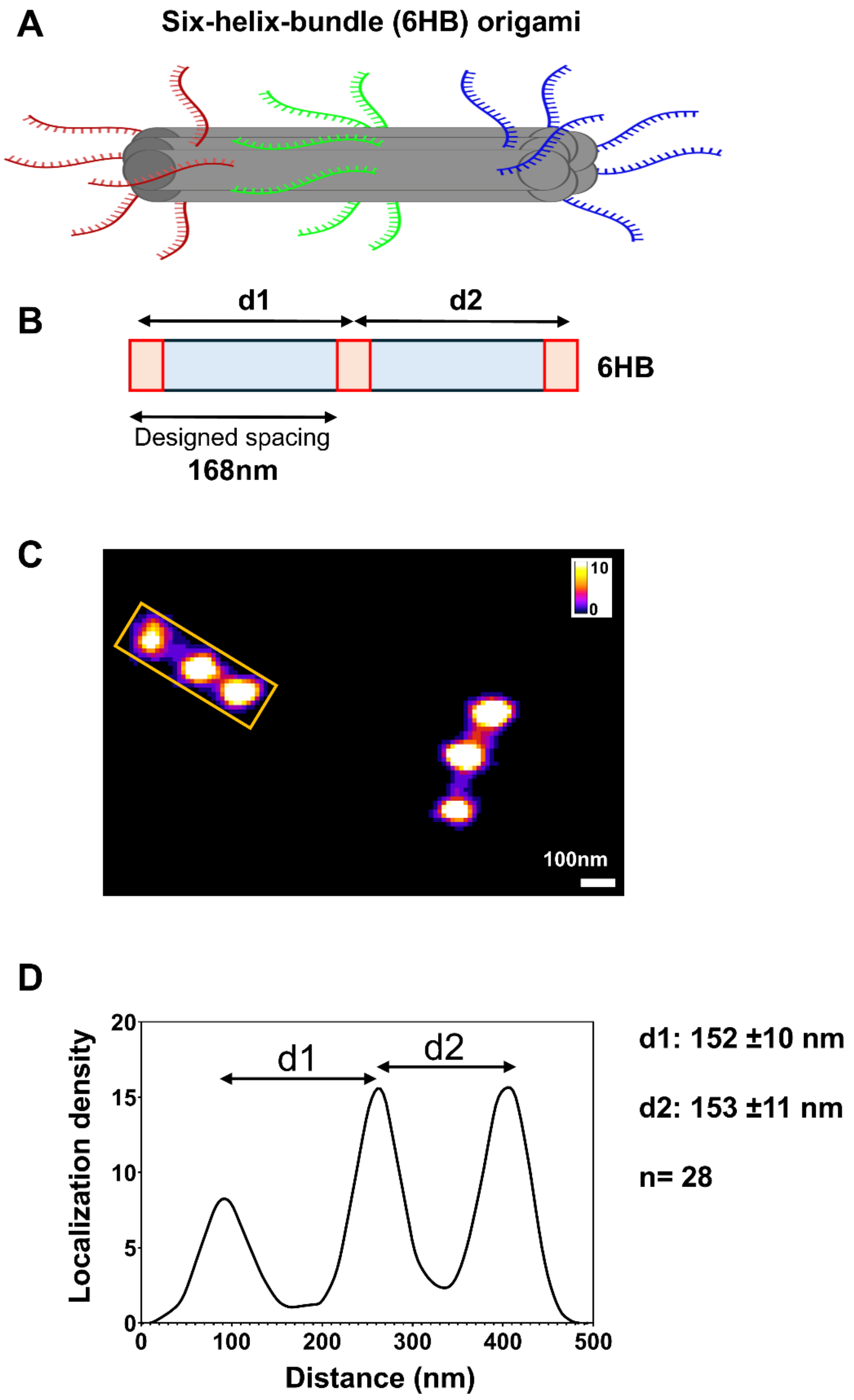

**Fig. S5. Validation of distance measurements using six-helix-bundle (6HB) DNA origami.** (**A**) Schematic representation of the six-helix-bundle (6HB) DNA origami structure. Three spatially separated target regions were positioned along the long axis of the nanostructure, with each target region containing six target sequences for 3WJ probe binding.

(**B**) Simplified schematic of the 6HB origami used to validate the super-resolved structure. The target regions were arranged along the long axis of the nanostructure. The designed spacing of the origami is indicated, while d1 and d2 denote the center-to-center distances between adjacent target regions measured from super-resolution images.
(**C**) Representative super-resolution image of a randomly 6HB DNA origami structure showing three resolved localization clusters corresponding to the three target regions. The boxed structure was selected for distance analysis. Scale bar, 100 nm.
(**D**) Localization density profile measured along the long axis of the boxed structure shown in (C). Distances between adjacent localization clusters (d1 and d2) were determined from peak-to-peak measurements of the localization density profile. Quantification across 28 structures yielded measured distances of 153 ± 11 nm (mean ± SD), compared with the designed spacing of approximately 168 nm. Despite the relatively small number of target sequences (six per target region), super-resolution imaging resolved the three target regions as distinct localization clusters, enabling nanoscale distance measurements.

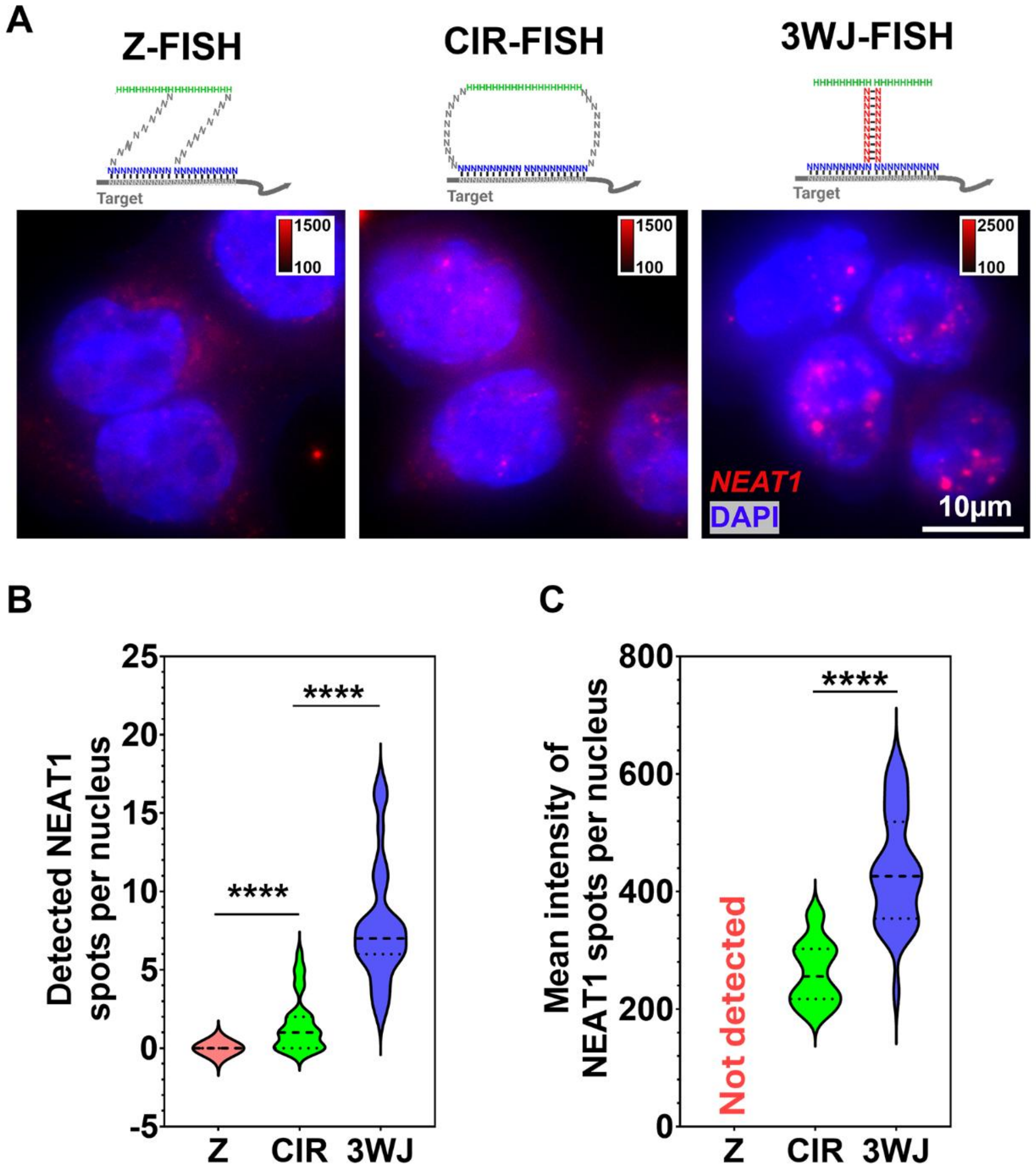

**Fig. S6. 3WJ amplification enhances *NEAT1* detection.**

(**A**) Representative fluorescence images of *NEAT1* detected using 16× amplified Z-FISH, CIR-FISH, and 3WJ-FISH. Scale bar, 10 μm. (**B**) Quantification of detected *NEAT1* spots per nucleus. Data were analyzed using a Kruskal–Wallis test followed by Dunn's multiple-comparisons test (3WJ versus CIR and 3WJ versus Z; ****$P < 0.0001$). n = 32, 28, and 38 nuclei for 3WJ, CIR, and Z, respectively. (**C**) Mean intensity of detected *NEAT1* spots per nucleus. No spots were detected in the Z-FISH condition; therefore, Z-FISH was excluded from the intensity comparison. n = 32 and 18 nuclei for 3WJ and CIR, respectively. Statistical significance was determined using a two-tailed unpaired Welch's t test (****$P < 0.0001$).

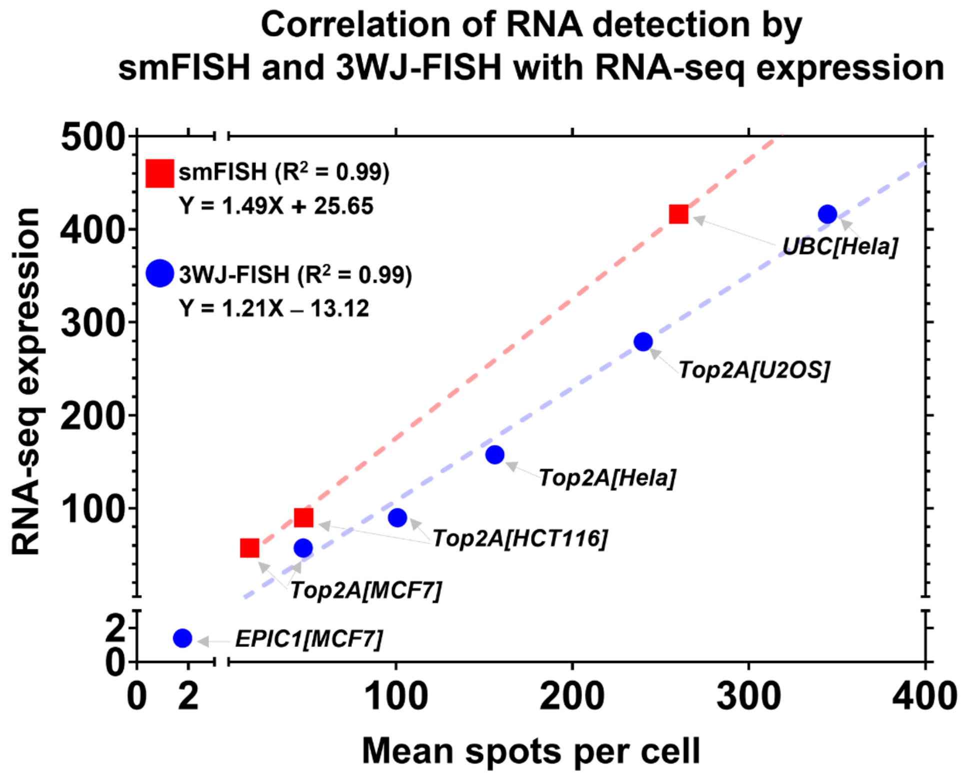

**Fig. S7. Comparison of RNA detection by conventional smFISH and 3WJ-FISH.**

Detection of *UBC*, *TOP2A*, and the low-abundance lncRNA *EPIC1* using conventional smFISH and 3WJ-FISH. Conventional smFISH detected fewer *UBC* and *TOP2A* RNA spots than 3WJ-FISH. The dashed line establishes a correlation between 3WJ-FISH or smFISH measurements (mean spots per cell) and RNA-seq expression levels (transcripts per million, TPM) across multiple genes and cell lines.

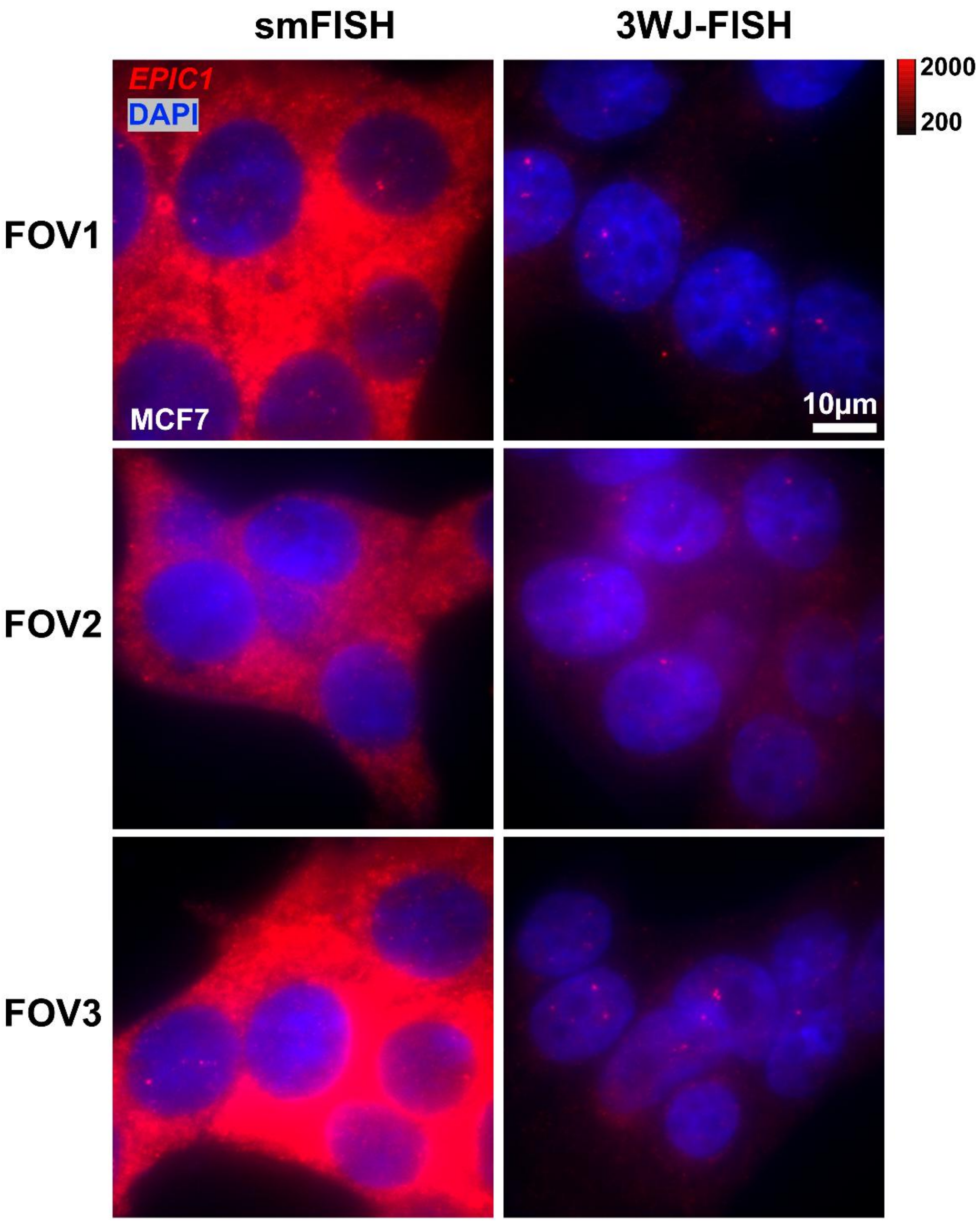

**Fig. S8. Comparison of lncRNA *EPIC1* detection by conventional smFISH and 3WJ-FISH in MCF7 cells.**

Representative images from three independent fields of view (FOV1–FOV3) showing *EPIC1* detection by conventional smFISH (left) and 3WJ-FISH (right). Conventional smFISH exhibited abundant cytoplasmic fluorescence signals, whereas 3WJ-FISH detected discrete *EPIC1* spots predominantly localized within the nuclei. *EPIC1* is shown in red, and nuclei are counterstained with DAPI (blue). Images from both methods are displayed using the same fluorescence intensity scale, as indicated by the shared calibration bar. Scale bar, 10 μm.

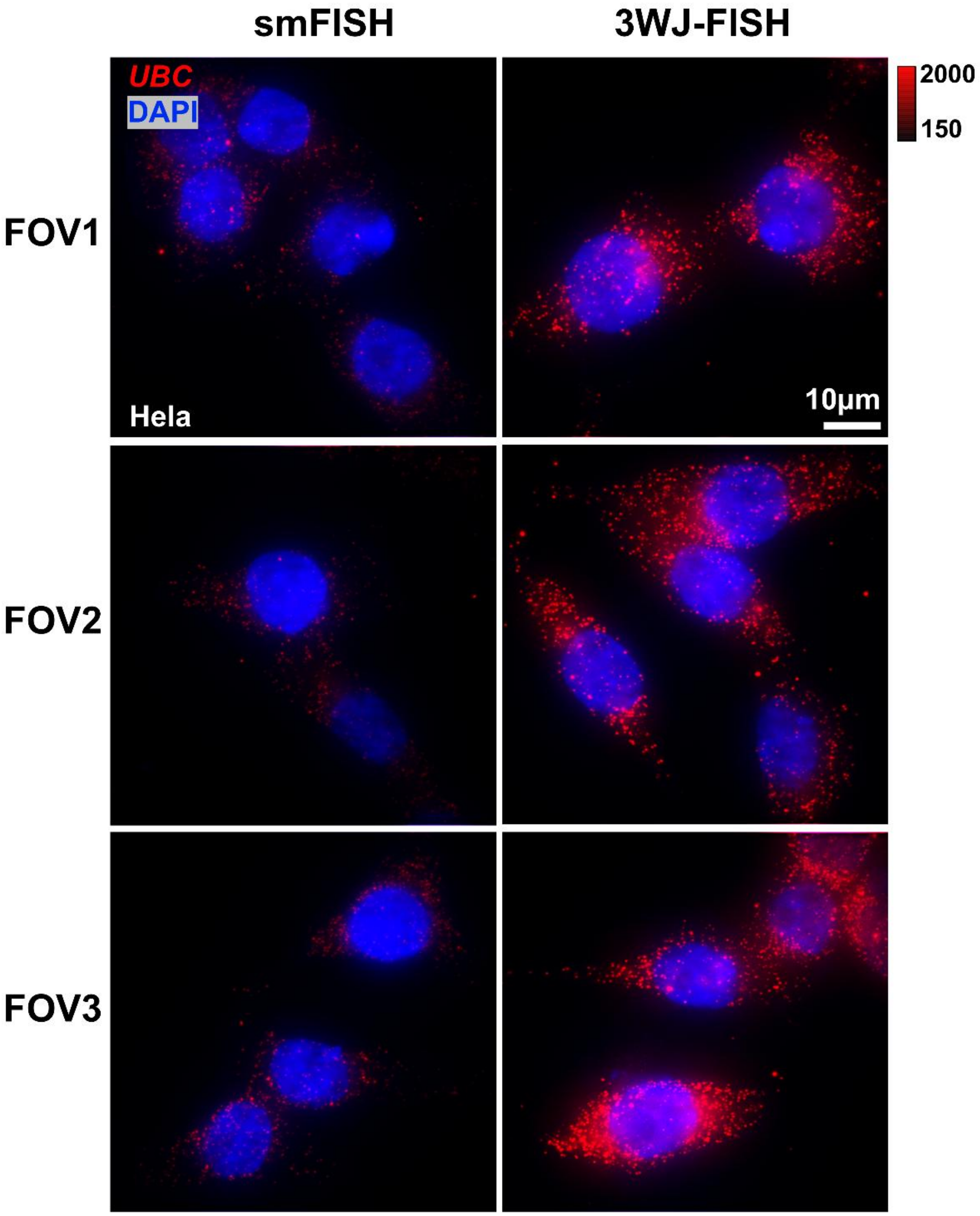

**Fig. S9. Additional representative images of *UBC* detection by conventional smFISH and 3WJ-FISH in HeLa cells.**

Representative images from three independent fields of view (FOV1–FOV3) showing *UBC* detection by conventional smFISH (left) and 3WJ-FISH (right). *UBC* is shown in red, and nuclei are counterstained with DAPI (blue). Images from both methods are displayed using the same fluorescence intensity scale, as indicated by the shared calibration bar. Scale bar, 10 μm.

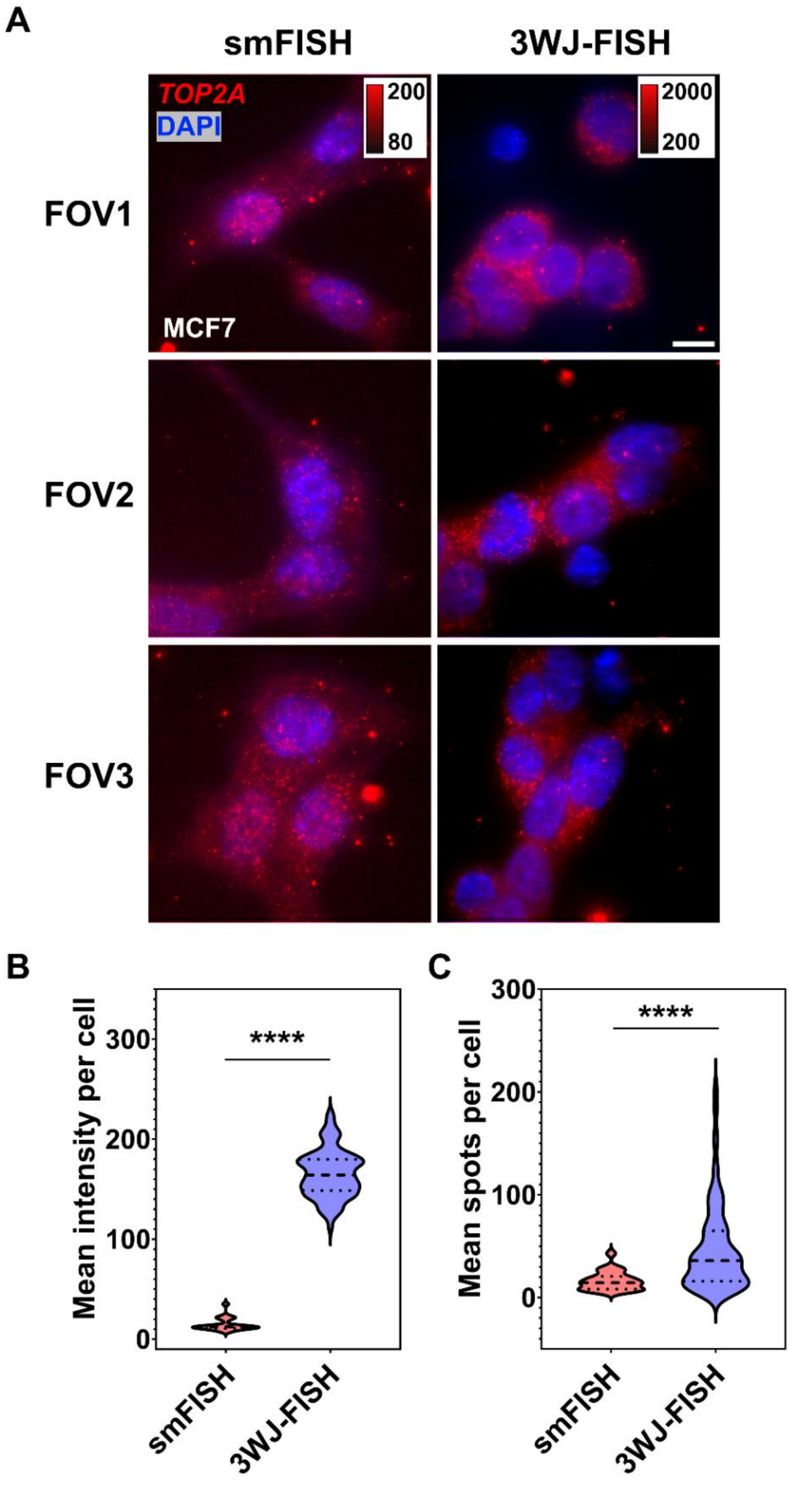

**Fig. S10. Comparison of TOP2A detection by conventional smFISH and 3WJ-FISH in MCF7 cells**

(**A**) Representative images from three independent fields of view (FOV1–FOV3) showing *TOP2A* detection by conventional smFISH (left) and 3WJ-FISH (right). (**B** and **C**) Violin plots comparing the mean fluorescence intensity (B) and mean RNA spot counts per cell (C) between conventional smFISH and 3WJ-FISH. *TOP2A* is shown in red, and nuclei are counterstained with DAPI (blue). Images are displayed using the fluorescence intensity scales indicated by the corresponding calibration bars. Scale bar, 10 µm. Statistical significance was determined using a two-tailed Mann–Whitney U test (smFISH, n = 24 cells; 3WJ-FISH, n = 146 cells; ****$P < 0.0001$ for both comparisons).

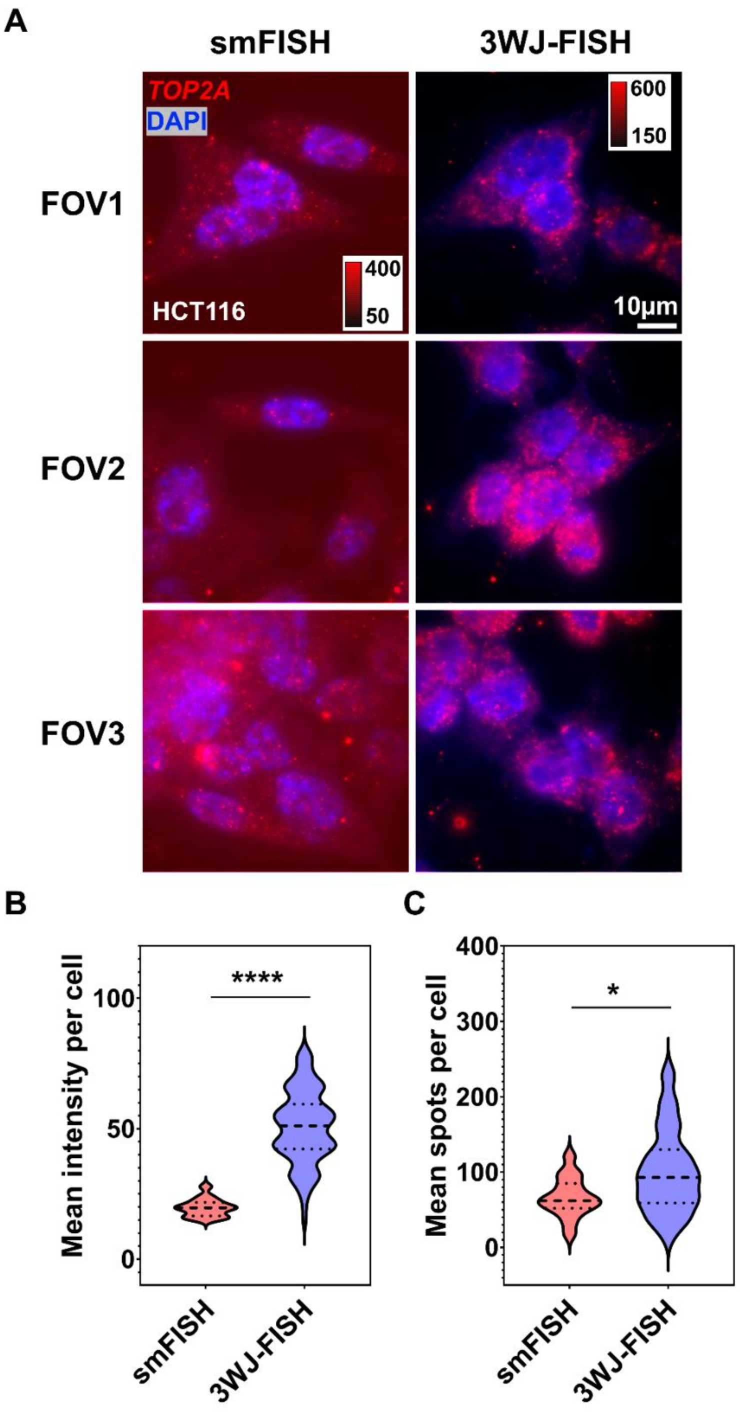

**Fig. S11. Comparison of *TOP2A* detection by conventional smFISH and 3WJ-FISH in HCT116 cells.**

(**A**) Representative images from three independent fields of view (FOV1–FOV3) showing TOP2A detection by conventional smFISH (left) and 3WJ-FISH (right). (**B, C**) Violin plots comparing the mean fluorescence intensity (B) and mean RNA spot counts per cell (C) between conventional smFISH and 3WJ-FISH. *TOP2A* is shown in red, and nuclei are counterstained with DAPI (blue). Images are displayed using the fluorescence intensity scales indicated by the corresponding calibration bars. Scale bar, 10 µm. Statistical significance was determined using a two-tailed Mann–Whitney U test (smFISH, n = 15 cells; 3WJ-FISH, n = 103 cells; ****$P <$ 0.0001 for fluorescence intensity; *$P$ = 0.0169 for RNA spot counts).

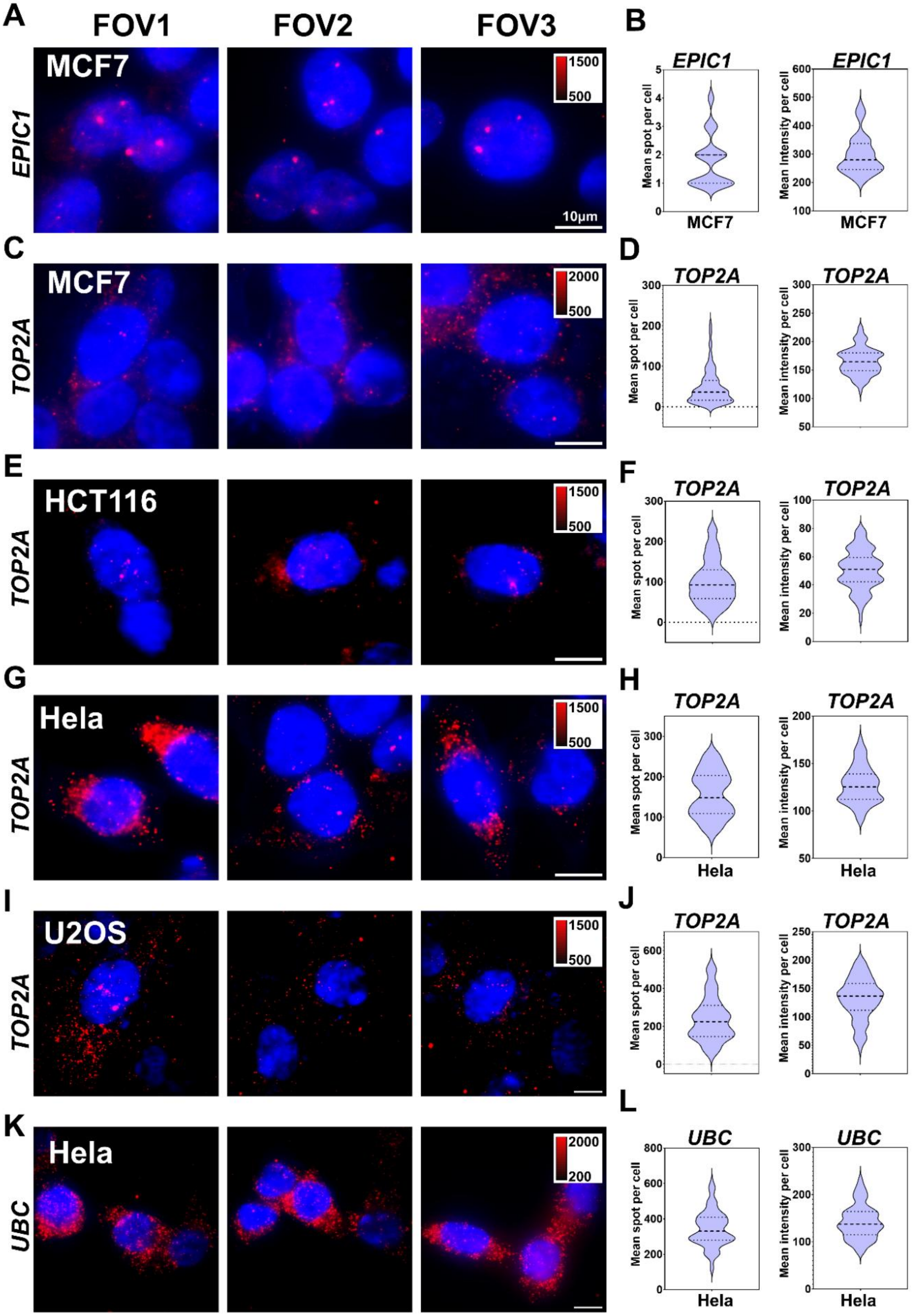
A
FOV1
FOV2
FOV3
MCF7
EPIC1
1500
500
10μm
B
EPIC1
Mean spot per cell
MCF7
Mean intensity per cell
C
MCF7
TOP2A
2000
500
D
TOP2A
E
HCT116
TOP2A
1500
500
F
TOP2A
G
Hela
TOP2A
H
Hela
I
U2OS
TOP2A
J
K
Hela
UBC
2000
200
L
UBC

**Fig. S12. Raw single-cell distributions of RNA spot counts and intensities corresponding to Fig. 3D.**

(**A**, **C, E, G, I, K**) Representative widefield images of RNA detection using 3WJ-FISH across indicated genes and cell lines, including *EPIC1* (MCF7), *TOP2A* (MCF7, HCT116, HeLa, and U2OS), and *UBC* (HeLa). Red signals indicate RNA spots; nuclei are counterstained with DAPI (blue). Multiple fields of view (FOVs) are shown for each condition. Intensity scales are indicated in each panel. Scale bars, 10 μm.
(**B, D, F, H, J, L**) Violin plots showing the distribution of RNA spot counts per cell (left) and mean intensity per cell (right) for the corresponding conditions. Dashed lines indicate median and interquartile ranges. These single-cell measurements form the basis for summary statistics (Table S13) and correlation analysis with RNA-seq data (Figure 3D in the main text).

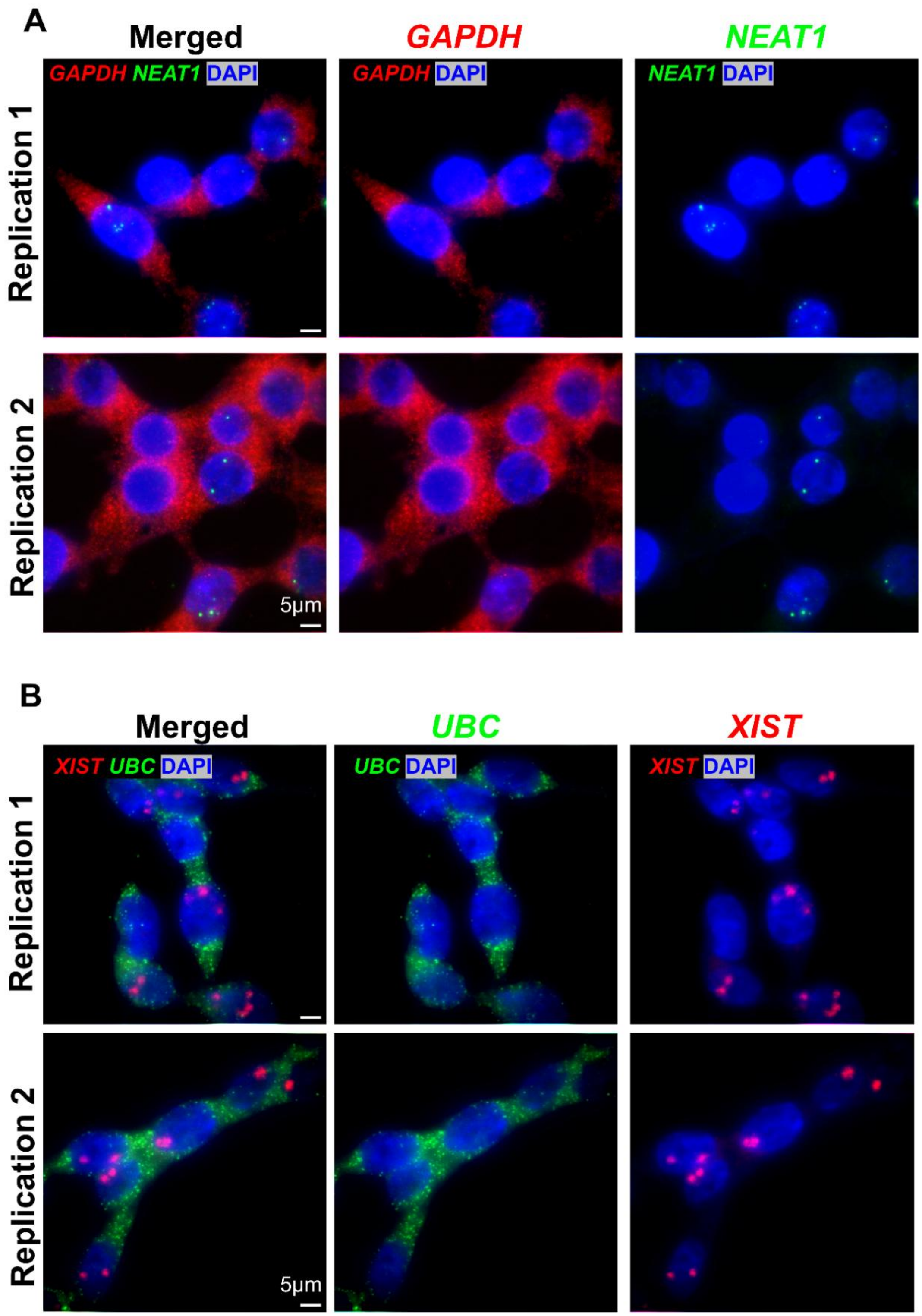

**Fig. S13. Multiplex RNA imaging of distinct transcript classes using 3WJ-FISH.**

**(A)** Simultaneous detection of *GAPDH* mRNA (red) and *NEAT1* lncRNA (green) in HEK293T cells using 3WJ-FISH. Representative images from two independent experiments are shown. Merged and single-channel views are displayed. Scale bar, 5 μm.

**(B)** Simultaneous detection of *UBC* mRNA (green) and *XIST* lncRNA (red) in HEK293T cells using 3WJ-FISH. Representative images from two independent experiments are shown. Merged and single-channel views are displayed. Scale bar, 5 µm. These examples demonstrate the compatibility of 3WJ-FISH with multiplex detection of both messenger RNAs (mRNAs) and long noncoding RNAs (lncRNAs) within the same cells.

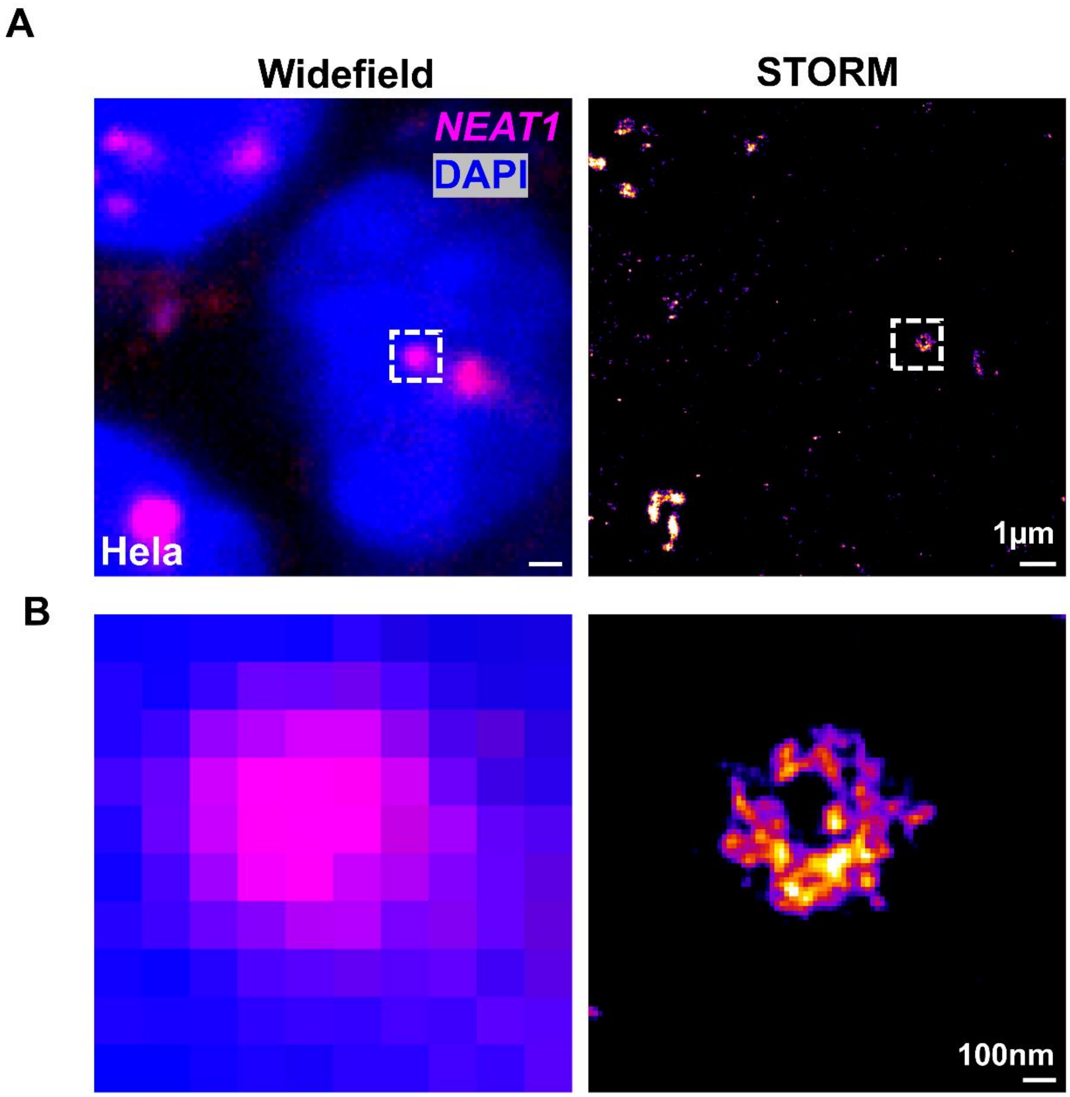

**Fig. S14. Super-resolution imaging of *NEAT1* RNA in HeLa cells using 3WJ-FISH.**
(**A**) Representative widefield (WF) and super-resolution (STORM) images of *NEAT1* RNA (magenta) in HeLa cells, with nuclei counterstained by DAPI (blue). The dashed boxes indicate regions selected for magnification. While WF imaging shows diffraction-limited NEAT1 foci, STORM resolves these structures into sub-diffraction localization clusters. Scale bar, 1 µm.
(**B**) Magnified views of the boxed regions in (A). WF imaging displays blurred, pixel-limited signal, whereas STORM reveals fine structural organization of *NEAT1* at the nanoscale. Scale bar, 100 nm.

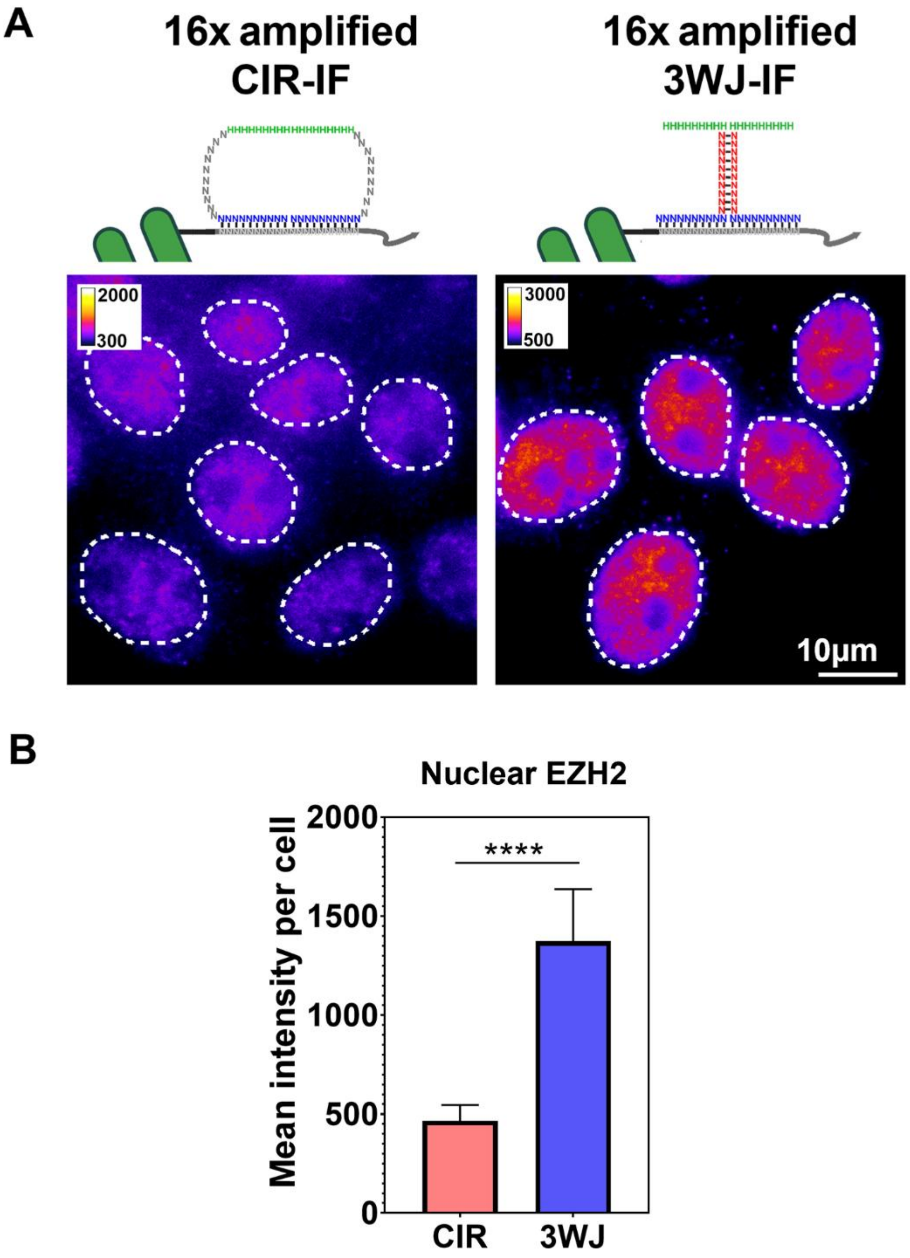

**Fig. S15. 3WJ-IF amplification enhances the detection of nuclear EZH2.**

(**A**) Representative fluorescence images of EZH2 detected using 16× amplified CIR-IF and 16× amplified 3WJ-IF. Nuclear boundaries are indicated by white dashed lines. Scale bar, 10 µm. (**B**) Quantification of the mean nuclear EZH2 fluorescence intensity. Data are presented as mean ± SD (n = 55 nuclei per condition). Statistical significance was determined using a two-tailed unpaired Welch's t-test. *****P* < 0.0001.

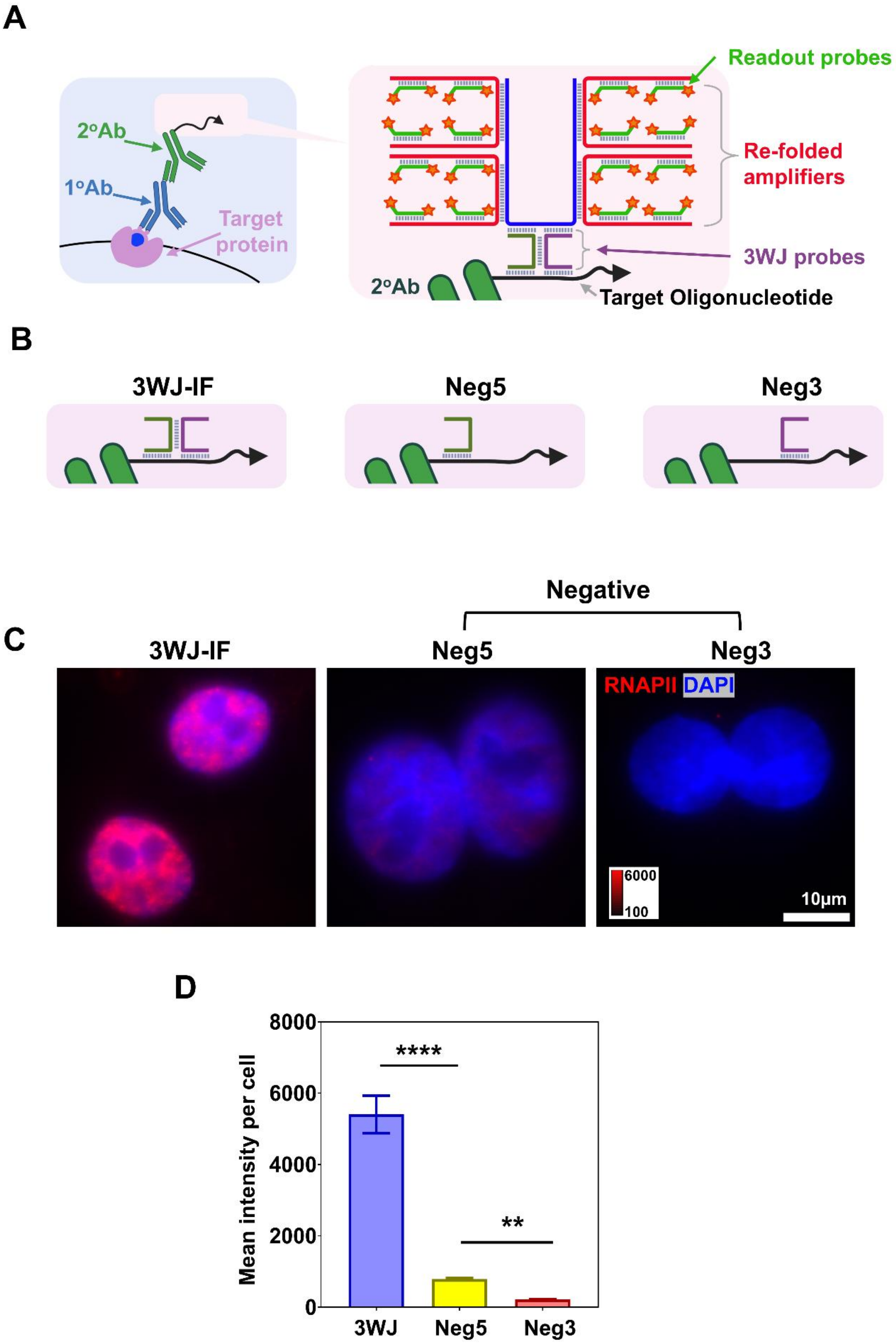
A
2°Ab
1°Ab
Target protein
Readout probes
Re-folded amplifiers
3WJ probes
2°Ab
Target Oligonucleotide
B
3WJ-IF
Neg5
Neg3
C
Negative
3WJ-IF
Neg5
Neg3
RNAPII DAPI
6000
100
10μm
D
Mean intensity per cell
8000
6000
4000
2000
0
****
**
3WJ
Neg5
Neg3

**Fig. S16. Specificity controls for 3WJ-mediated immunofluorescence (3WJ-IF).**

(**A**) Schematic illustration of the 3WJ-IF strategy. Antibody-conjugated target oligonucleotides recruit split-probe pairs, enabling target-dependent 3WJ formation and subsequent hybridization of pre-folded amplifiers and fluorescent readout probes.

(**B**) Design of split-probe control conditions. In the Neg5 control, the 3′ split probe was omitted, whereas in the Neg3 control, the 5′ split probe was omitted. In both cases, failure to assemble the complete probe pair prevents efficient 3WJ formation and signal amplification.

(**C**) Representative images of RNA polymerase II (RNAPII) detection in MCF7 cells under complete 3WJ-IF conditions and the corresponding split-probe control conditions. Strong nuclear labeling was observed only under complete probe-pair assembly conditions, whereas Neg5 and Neg3 controls showed substantially reduced signal. Scale bar, 10 μm.

(**D**) Quantification of mean RNAPII fluorescence intensity per cell under 3WJ-IF, Neg5, and Neg3 conditions. Data are presented as mean ± SD. Statistical significance was assessed using Welch's ANOVA followed by Dunnett's T3 multiple-comparisons test (**$P < 0.01$; ****$P < 0.0001$).

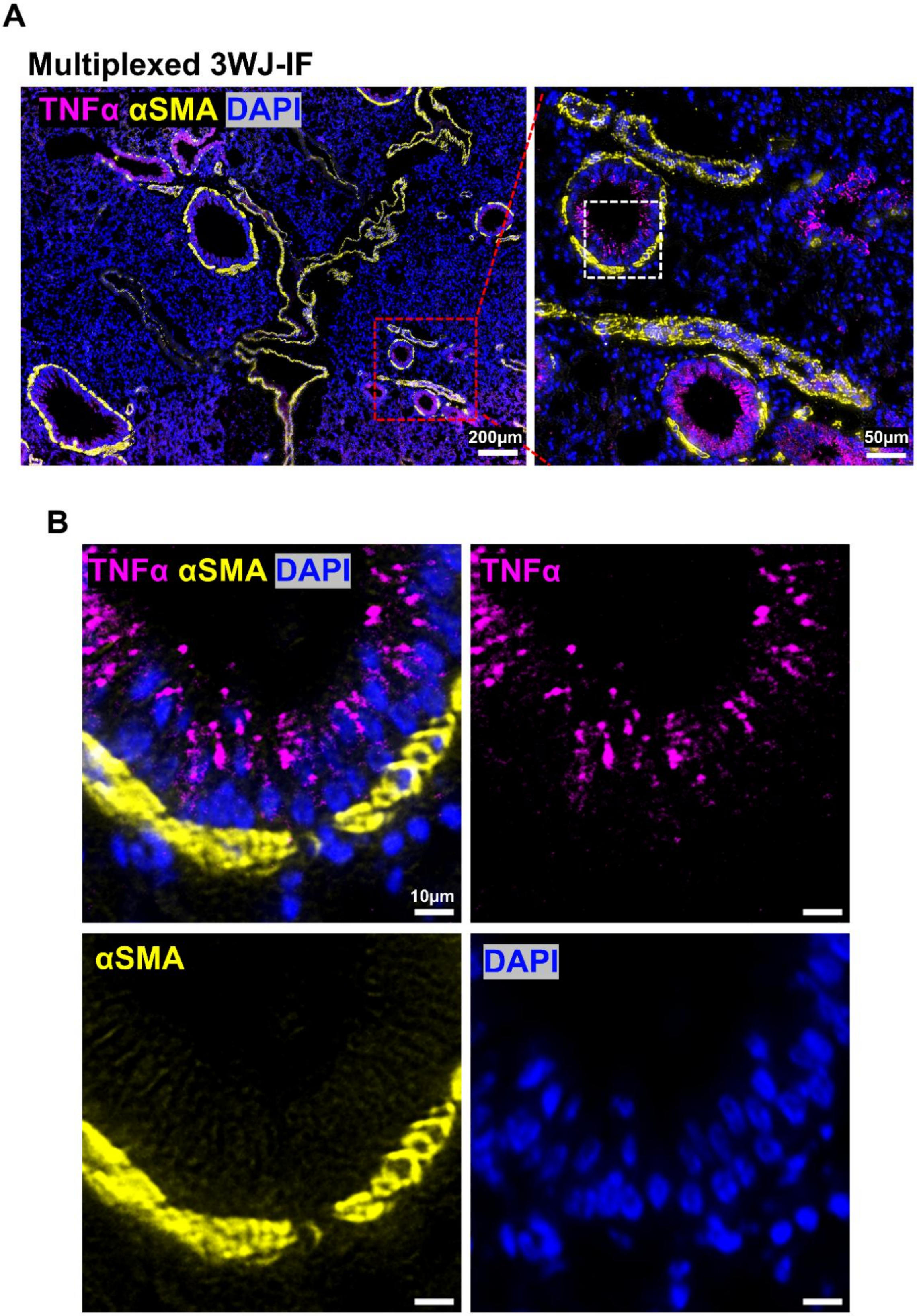
A
Multiplexed 3WJ-IF
TNFα αSMA DAPI
200μm
50μm
B
TNFα αSMA DAPI
10μm
TNFα
αSMA
DAPI

**Fig. S17. Multiplexed 3WJ-IF imaging of TNFα and αSMA in FFPE mouse lung tissue.**

**(A)** Representative low- and high-magnification views of multiplexed 3WJ-IF imaging showing simultaneous detection of TNFα (magenta) and αSMA (yellow) in FFPE mouse lung tissue sections. Insets indicate the magnified regions shown on the right. The image on the right is also presented in Fig. 1E, showing the same region from the larger field of view on the left. Scale bars, 200 µm (left) and 50 µm (right).

**(B)** High-magnification view of an airway region showing the merged image and corresponding single-channel images. TNFα and αSMA signals were detected simultaneously with minimal channel cross-talk, demonstrating the compatibility of 3WJ-IF for multiplexed protein imaging in complex tissue environments. Scale bars, 10 µm.

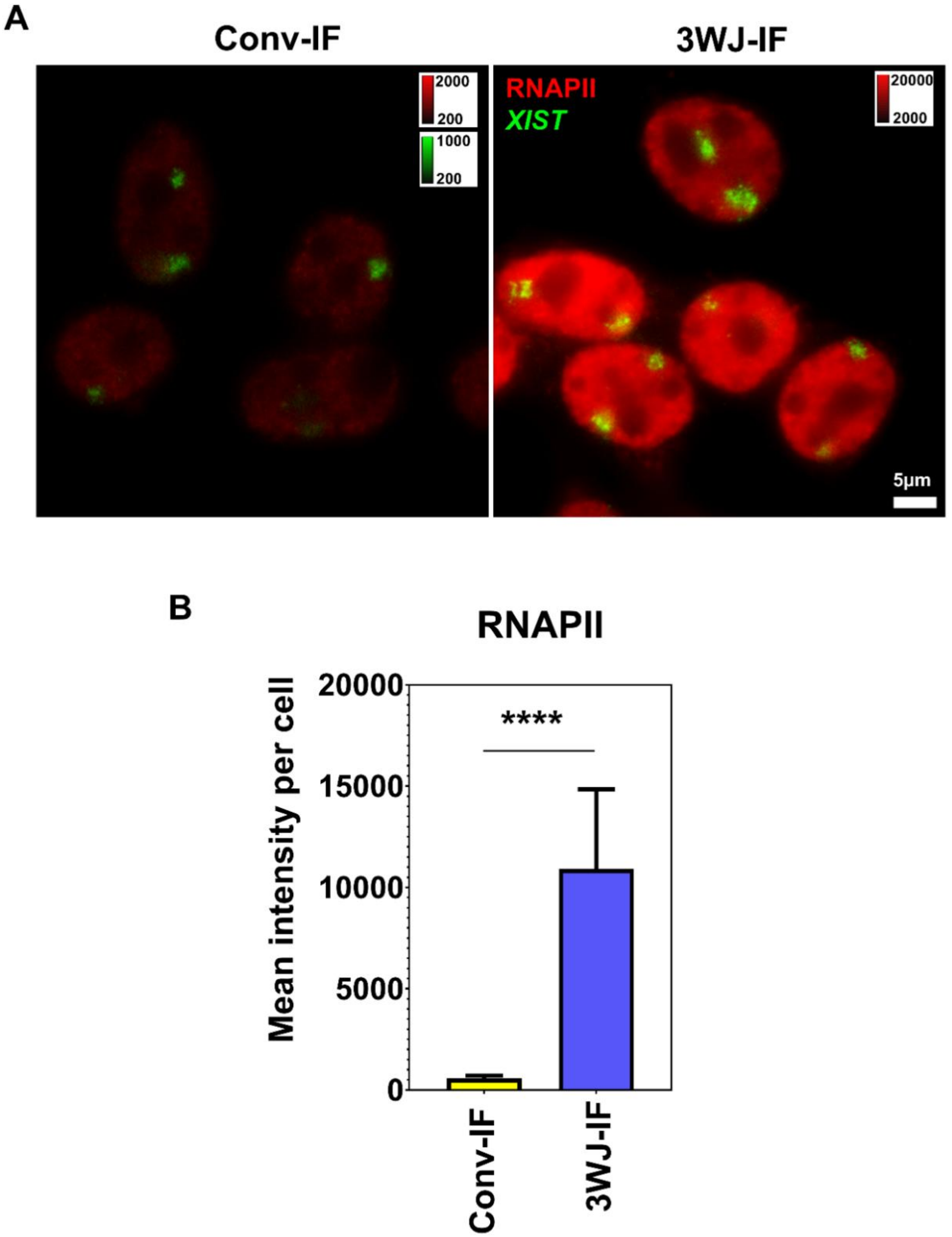

**Fig. S18. Application of 3WJ-IF for RNA–protein co-detection.**

**(A)** Representative images of simultaneous detection of *XIST* RNA (green) and RNA polymerase II (RNAPII, red) in HEK293T cells. *XIST* RNA was detected using 3WJ-FISH, whereas RNAPII was detected using either conventional immunofluorescence (Conv-IF) or 3WJ-mediated immunofluorescence (3WJ-IF). Representative images are displayed using identical imaging and display settings within each channel. Scale bar, 5 μm. **(B)** Quantification of mean RNAPII fluorescence intensity per cell for Conv-IF and 3WJ-IF. Data are presented as mean ± SD. Statistical significance was determined using a two-tailed Welch's t-test (****$P < 0.0001$).

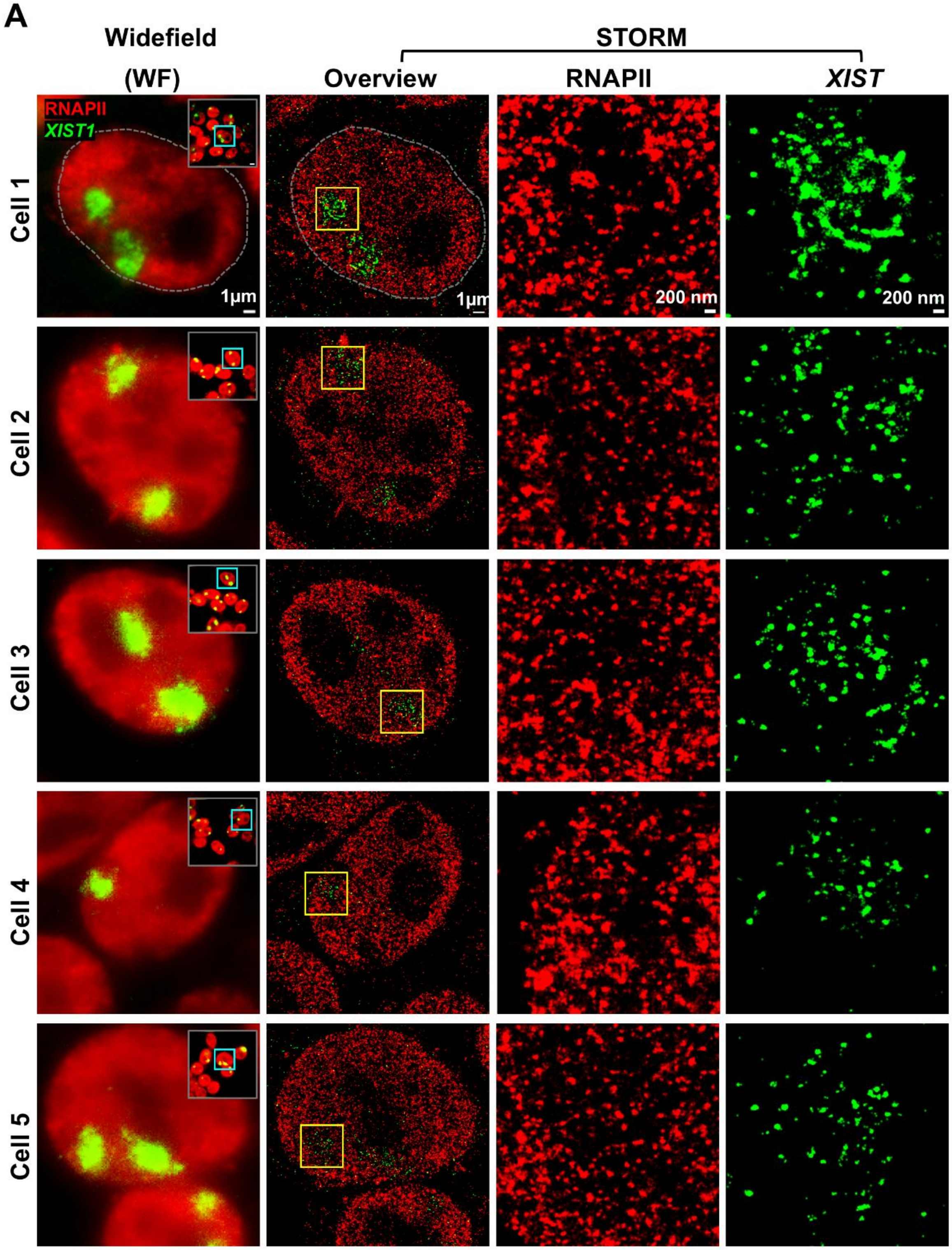

**Fig. S19. Super-resolution imaging of *XIST* RNA and RNAPII organization within the Xi regions.**

(**A**) Representative wide-field (WF) and STORM imaging of *XIST* RNA (green) and RNAPII protein (red) in HEK293T cells. Images are shown for one representative cell selected from each field of view (FOV), where each FOV contains multiple cells. For each FOV, a full WF image is shown as an inset, with the selected cell indicated by a cyan box. The corresponding single-cell STORM reconstruction is displayed, with the Xi region highlighted by a yellow box. The selected Xi region is further magnified to show RNAPII and *XIST* signals at super-resolution. The images of Cell 1 are also presented in Fig. 1G, with the STORM images here to align with the corresponding WF images. Scale bars: 1 µm (WF and STORM overview) and 200 nm (zoomed images).

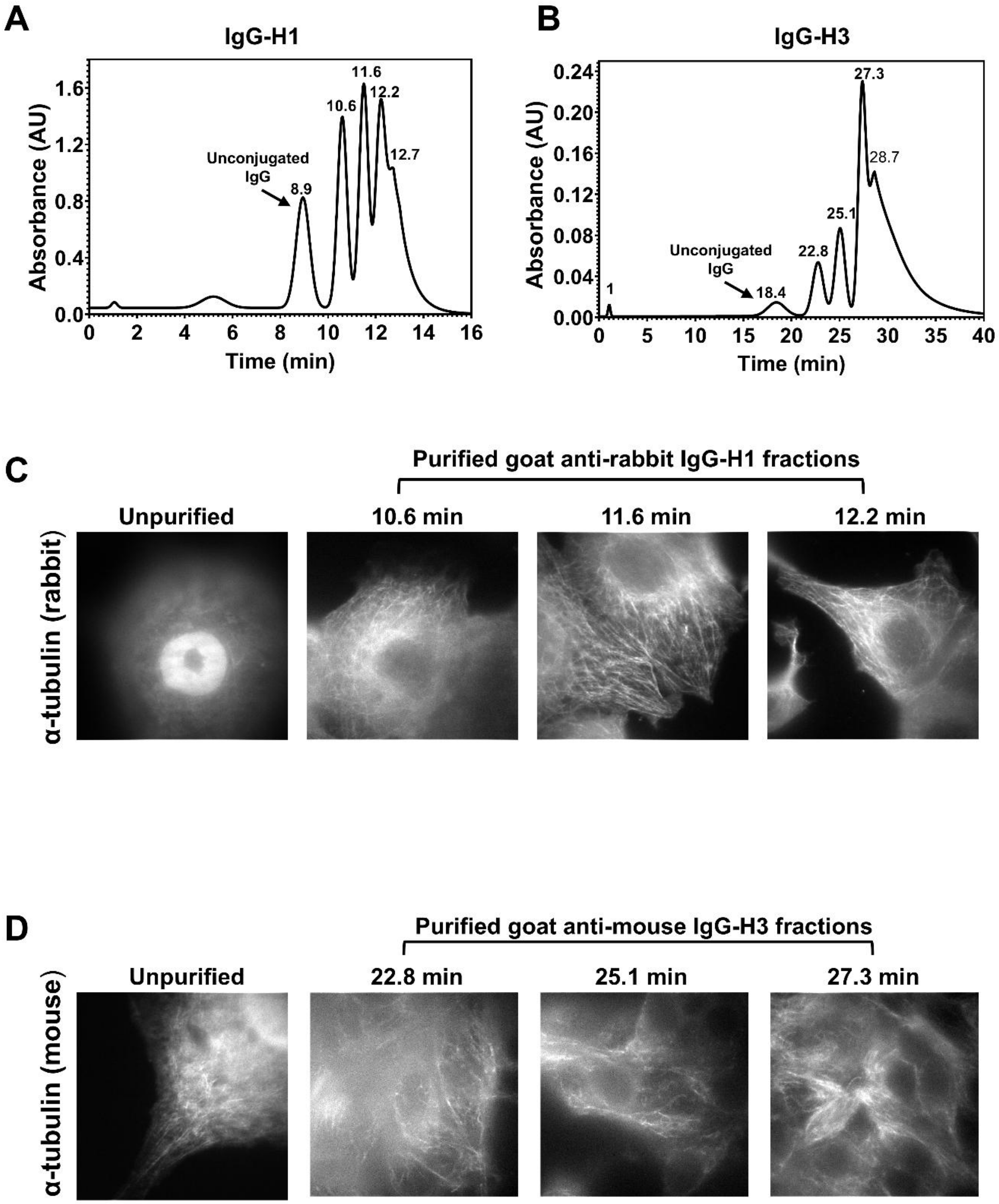

**Fig. S20. IEX-HPLC purification of DNA-conjugated secondary antibodies improves immunofluorescence labeling specificity.**

(**A-B**) Representative ion-exchange high-performance liquid chromatography (IEX-HPLC) chromatograms of DNA-conjugated secondary antibodies prepared for 3WJ-IF, including goat anti-rabbit IgG conjugated with DNA probe H1 (A) and goat anti-mouse IgG conjugated with DNA probe H3 (B). Chromatograms were monitored by UV absorbance at 280 nm. Peaks corresponding to unconjugated antibodies and DNA–antibody conjugates are indicated, with later-eluting peaks corresponding to conjugated species.

(**C**) Representative immunofluorescence images of α-tubulin in MCF7 cells labeled using unpurified goat anti-rabbit IgG–H1 conjugates or purified fractions collected at the indicated retention times (10.6, 11.6, and 12.2 min). Purification reduced diffuse and nuclear-associated background while preserving cytoskeletal labeling.
(**D**) Representative immunofluorescence images of α-tubulin in MCF7 cells labeled using unpurified goat anti-mouse IgG–H3 conjugates or purified fractions collected at the indicated retention times (22.8, 25.1, and 27.3 min). Purified fractions exhibited reduced nonspecific background and improved visualization of microtubule structures.

### Supplementary Tables

| Name | Component | Set | Binding target | Length (nt) | Sequence (5’→3’) | Modification | Application (Figures) |
|---|---|---|---|---|---|---|---|
| P-Amp-1a-16x | Primary amplifier | 1a | HBS (on 3WJ probe) | 100 | +GAAT+CTTA+AACC+CAAC+TTttt +GAAT+CTTA+AACC+CAAC+TTtttgtaggttg aaggttgagtt+GAAT+CTTA+AACC+CAAC+TT ttt+GAAT+CTTA+AACC+CAAC+TT | LNA (internal) | Fig. 1D, Fig. 2A, E (right); Fig. 3A(right); Fig. 3E (right); Fig. 4H; Fig. 5B-C, F; Fig. 6A; Fig. 6C; Fig. S2B (right), S2E (right); Fig. S3B (left); Fig. S4; Fig. S5; Fig. S6; Fig. S8 (right); Fig. S9 (right); Fig. S10 (right); Fig. S11 (right); Fig. S12; Fig. S13; Fig. S14; Fig. S15; Fig. S16; Fig. S17; Fig. S18; Fig. S19; Fig. S20D |
| P-Amp-1a-16x-noLNA | Primary amplifier | 1a | HBS (on 3WJ probe) | 100 | GAATCTTAAACCCAACTTtttGAATCTTAA ACCCAACTTtttgtaggttgaaggttgagttGAATCTT AAACCCAACTTtttGAATCTTAAACCCAAC TT | None (DNA) | Fig. 2E (left); Fig. S3B (right) |
| P-Amp-1c-16x | Primary amplifier | 1c | HBS (on 3WJ probe) | 100 | +GAAT+CTTA+AACC+CAAC+TTttt+GAAT +CTTA+AACC+CAAC+TTtttgtgtctccatagtatgc tt+GAAT+CTTA+AACC+CAAC+TTttt+GAA T+CTTA+AACC+CAAC+TT | LNA (internal) | Fig. S2B (left), S2E (left) |
| P-Amp-1a-8x | Primary amplifier | 1a | HBS (on 3WJ probe) | 58 | +GAAT+CTTA+AACC+CAAC+TTtttgtaggttg aaggttgagtt+GAAT+CTTA+AACC+CAAC+TT | LNA (internal) | Fig. 3E (middle) |
| S-Amp-1a | Secondary amplifier | 1a | 1° amplifier | 100 | ACTAAGTGGTAGATCAGCtttACTAAGTGG TAGATCAGCtta+ag+tt+gg+gt+tt+aa+ga+tt+ctt ACTAAGTGGTAGATCAGCttt ACTAAGTGGTAGATCAGC | LNA (internal) | Fig. 1D, Fig. 2A, E (right); Fig. 3A(right); Fig. 3E (right); Fig. 4A; Fig. 4H; Fig. 5B-C, F; Fig. 6A; Fig. 6C; Fig. S2B, S2E; Fig. S3B (left); Fig. S4; Fig. S5; Fig. S6; Fig. S8 (right); Fig. S9 (right); Fig. S10 (right); Fig. S11 (right); Fig. S12; Fig. S13; Fig. S14; Fig. S15; Fig. S16; Fig. S17; Fig. S18; Fig. S19; Fig. S20D |
| S-Amp-1a-noLNA | Secondary amplifier | 1a | 1° amplifier | 100 | ACTAAGTGGTAGATCAGCtttACTAAGTGG TAGATCAGCttaagttgggtttaagattcttACTAAGT GGTAGATCAGCtttACTAAGTGGTAGATCA GC | None (DNA) | Fig.2E (left); Fig. S3B (right) |
| RO-1a | Readout probe (3WJ) | 1a | 2° amplifier | 20 | /5Alex647N/-GCTGATCTACCACTTAGTTA- /3AlexF647N/ | 5′ and 3′ Alexa Fluor 647 labels | Fig. 1D, Fig. 2A, E; Fig. 3A(right); Fig. 4A; Fig. 4H; Fig. 5B-C, F; Fig. 6A; Fig. 6C; Fig. S2B, S2E; Fig. S3B; Fig. S4; Fig. S5; Fig. S6; Fig. S8 (right); Fig. S9 (right); Fig. S10 (right); Fig. S11 (right); Fig. S12; Fig. S13; Fig. S14; Fig. S15; Fig. S16; Fig. S17; Fig. S18; Fig. S19; Fig. S20D |
| RO-1a-1dye | Readout probe (3WJ) | 1a | 2° amplifier | 20 | /5Alex647N/-GCTGATCTACCACTTAGTTA | 5′ Alexa Fluor 647 labels | Fig. 3E (middle, right) |
| RO-1a-1x | Readout probe (3WJ) | 1a | HBS (on 3WJ probe) | 19 | /5Alex647N/-TTGTAGGTTGAAGGTTGAG | 5′ Alexa Fluor 647 labels | Fig. 3E (left) |

| | | | | | | | |
|---|---|---|---|---|---|---|---|
| P-Amp-2a-16x | Primary amplifier | 2a | HBS (on 3WJ probe) | 100 | TA+GTTTA+GAATC+ATCTC+GtttTA+GTTTA+GAATC+ATCTC+Gttagtagagtggaatgaaggt tTA+GTTTA+GAATC+ATCTC+GtttTA+GTTTA+GAATC+ATCTC+G | LNA (internal) | Fig. 6C; Fig. S13; Fig. S18; Fig. S19 |
| S-Amp-2a | Secondary amplifier | 2a | 1° amplifier | 100 | GCGCGATTGACCGTCTCGtttGCGCGATTGACCGTCTCGtt+cgaga+tgatt+ctaaa+ctattGCGCGATTGACCGTCTCGtttGCGCGATTGACCGTCTCG | LNA (internal) | Fig. 6C; Fig. S13; Fig. S18; Fig. S19 |
| RO-2a | Readout probe (3WJ) | 2a | 2° amplifier | 20 | /5CF568/-AACGAGACGGTCAATCGCGC-/3CF568/ | 5′ and 3′ CF568 labels | Fig. 6C; Fig. S13; Fig. S18; Fig. S19 |
| P-Amp-2b-16x | Primary amplifier | 2b | HBS (on 3WJ probe) | 98 | TA+CTATA+CAATC+ATCTC+CaaTA+CTATA+CAATC+ATCTC+Caaagtagagtggaatgaagg aaTA+CTATA+CAATC+ATCTC+CaaTA+CTATA+CAATC+ATCTC+C | LNA (internal) | Fig. 5F; Fig. S17; Fig. S20C |
| S-Amp-2b | Secondary amplifier | 2b | 1° amplifier | 100 | GTAGAGGGTGAATGTTGGaaaGTAGAGGGTGAATGTTGGaa+ggaga+tgatt+gtata+gtaaaGTAGAGGGTGAATGTTGGaaaGTAGAGGGTGAATGTTGG | LNA (internal) | Fig. 5F; Fig. S17; Fig. S20C |
| RO-2b-1dye | Readout probe (3WJ) | 2b | 2° amplifier | 18 | /5CF568/-CCAACATTCACCCTCTAC | 5′ CF568 labels | Fig. 5F; Fig. S17; Fig. S20C |
| P-Amp-1b-HBSopt | Primary amplifier (truncated) | 1b | HBS (on 3WJ probe) | 80 | +GAAT+CTTA+AACC+CAAC+TTttaccaccac caccaccaccacaacaacaacaacaacaacaatt+GAAT+CTTA+AACC+CAAC+TT | LNA (internal) | Fig. 4A |
| RO-smFISH | Readout probe (smFISH) | – | smFISH probe (on target RNA) | 20 | /5Alex647N/-GTGAAATCGCAGCAGCCCAC-/3AlexF647N/ | 5′ and 3′ Alexa Fluor 647 labels | Fig. 3A(Left); Fig. S8 (left); Fig. S9 (left); Fig. S10 (left); Fig. S11 (left) |

**Table S1. DNA sequences of amplifiers and readout probes.**

P-Amp, primary amplifier; S-Amp, secondary amplifier; RO, readout probe. Two orthogonal amplifier sets (Sets 1 and 2), each containing distinct variants, were designed for multiplexed detection (see Methods). Primary (1°) amplifiers bind to HBS domains formed upon target-dependent 3WJ assembly, whereas secondary (2°) amplifiers bind to 1° amplifiers to enable hierarchical signal amplification. Full-length 1° amplifiers contain four secondary-amplifier binding domains for 16× amplification, whereas the 8× variant (P-Amp-1a-8x) contains two binding domains for tunable signal amplification. A separate 1° amplifier (P-Amp-1b-HBSopt) with an extended HBS-binding region was used for HBS-length optimization experiments. Readout probes hybridize to 2° amplifiers for fluorescence detection, whereas RO-1a-1x binds directly to the HBS for 1× detection without amplification. The smFISH readout probe hybridizes directly to probe-associated domains on target RNA. All sequences are listed in the 5′ to 3′ direction, with lengths excluding fluorophore modifications. Uppercase letters indicate functional domains, whereas lowercase letters indicate binding and spacer sequences. "+" denotes locked nucleic acid (LNA) modifications introduced to enhance duplex stability and binding affinity.

| Name | Target | Sequence (5’→3’) | Application (Figures) |
|---|---|---|---|
| NEAT1-3WJ-3'-1 | *NEAT1* | gatcatttccagggctgctgcg CGCGCGGATC CAACCTACA | Specificity validation (Fig. 4G-H);<br><br>Multiplex 3WJ-FISH (Fig. S13A);<br><br>Super-resolution imaging (Fig. S14A-B)<br><br>Probe architecture comparison for RNA detection (Fig. S6) |
| NEAT1-3WJ-5’-1 | *NEAT1* | CTCAACCTT GATCCGCGCG gcctattcctcctgactcctcc | |
| NEAT1-3WJ-3'-2 | *NEAT1* | ttcacagcccttggtctgga CGCTTGTCGG CAACCTACA | |
| NEAT1-3WJ-5’-2 | *NEAT1* | CTCAACCTT CCGACAAGCG aaaaaaggggctgctggcat | |
| NEAT1-3WJ-3'-3 | *NEAT1* | cctgtcaaacatgctaggtg GAACGGATAC Caacctaca |  |
| NEAT1-3WJ-5’-3 | *NEAT1* | CTCAACCTT GTATCCGTTC ccgcaccagtcaccggtgag | |
| NEAT1-3WJ-3'-4 | *NEAT1* | gcactgccacctggaaaata GCCTCGATTA Caacctaca | |
| NEAT1-3WJ-5’-4 | *NEAT1* | CTCAACCTT TAATCGAGGC aagcgttggtcaatgttgtc | |
| NEAT1-3WJ-3'-5 | *NEAT1* | gagtgagctcacaagaagagtt CGACGGATGT Caacctaca | |
| NEAT1-3WJ-5’-5 | *NEAT1* | CTCAACCTT ACATCCGTCG tagcgccaaacctagagaaaag | |
| NEAT1-3WJ-3'-6 | *NEAT1* | cggtccatgaagcatttttgt AATTCGGCCG CAACCTACA | |
| NEAT1-3WJ-5’-6 | *NEAT1* | CTCAACCTT CGGCCGAATT aactttcagctggatggctgc | |
| NEAT1-3WJ-3'-7 | *NEAT1* | cctggaaacagaacattgga GCCCGTATTC CAACCTACA | |
| NEAT1-3WJ-5’-7 | *NEAT1* | CTCAACCTT GAATACGGGC gaacttcctcctcctaagcc | |
| NEAT1-3WJ-3'-8 | *NEAT1* | accggaggctcaatttagaag CCGCTTGCGA CAACCTACA | |
| NEAT1-3WJ-5’-8 | *NEAT1* | CTCAACCTT TCGCAAGCGG atgcagcatctgaaaaccttt | |
| NEAT1-3WJ-3'-9 | *NEAT1* | acagcttagggatcttcttgaa GTAGGGCAAT CAACCTACA | |
| NEAT1-3WJ-5’-9 | *NEAT1* | CTCAACCTT ATTGCCCTAC gcaaggttccaagcacaaaact | |
| NEAT1-3WJ-3’-10 | *NEAT1* | tacaaggcatcaatctgcg GCGGGAAGCA CAACCTACA | |
| NEAT1-3WJ-5’-10 | *NEAT1* | CTCAACCTT TGCTTCCCGC ttgtggcatcaacgttaaaatgt | |
| NEAT1-3WJ-3’-11 | *NEAT1* | cgccccaagttatttcatcagg CGTGGATTAG CAACCTACA | |
| NEAT1-3WJ-5’-11 | *NEAT1* | CTCAACCTT CTAATCCACG ctaagaacttctccgagaaacg | |
| NEAT1-3WJ-3’-12 | *NEAT1* | aggaatgaccaacttgtacc GGCATCGACC CAACCTACA | |
| NEAT1-3WJ-5’-12 | *NEAT1* | CTCAACCTT GGTCGATGCC ctcccagcgtttagcacaacacaatg | |
| NEAT1-3WJ-3’-13 | *NEAT1* | tcctagtaatctgcaatgcaat AGGGTCATCG CAACCTACA | |
| NEAT1-3WJ-5’-13 | *NEAT1* | CTCAACCTT CGATGACCCT cacaatgcccaaactagacctg | |
| NEAT1-3WJ-3’-14 | *NEAT1* | tttggttctcggaaaactggtga CCGTAAGGTG CAACCTACA | |
| NEAT1-3WJ-5’-14 | *NEAT1* | CTCAACCTT CACCTTACGG ctctgtgtcacctgttttcagtt | |
| NEAT1-3WJ-3’-15 | *NEAT1* | ctctgaaacaggctgtcttggccac GGACCGAGGG CAACCTACA | |
| NEAT1-3WJ-5’-15 | *NEAT1* | CTCAACCTT CCCTCGGTCC caggaggaagctggtaaagacattt | |
| NEAT1-3WJ-3’-16 | *NEAT1* | atccagaagagcccatctaat TTGTGTATCC CAACCTACA | |
| NEAT1-3WJ-5’-16 | *NEAT1* | CTCAACCTT GGATACACAA ctcgtcttaagtggtccctta | |
| NEAT1-3WJ-3’-17 | *NEAT1* | gctcatgattttcaatcacaag CGCCGGATCT CAACCTACA | |
| NEAT1-3WJ-5’-17 | *NEAT1* | CTCAACCTT AGATCCGGCG acacctgtgacaaatgaggaac | |
| NEAT1-3WJ-3’-18 | *NEAT1* | acagtgaccacaaaaggttatt TTGAATCAAC CAACCTACA | |
| NEAT1-3WJ-5’-18 | *NEAT1* | CTCAACCTT GTTGATTCAA tggttgttacaaagactacaat | |
| NEAT1-3WJ-3’-19 | *NEAT1* | tcccaacgacagtaattgtttgca TGCTAGTTCG CAACCTACA | |
| NEAT1-3WJ-5’-19 | *NEAT1* | CTCAACCTT CGAACTAGCA tcatccccaagtcattggttaaga | |
| NEAT1-3WJ-3’-20 | *NEAT1* | caacagcatacccgagactact ATCGCGCAAT CAACCTACA | |
| NEAT1-3WJ-5’-20 | *NEAT1* | CTCAACCTT ATTGCGCGAT tccccatacatgcgtgactaat | |
| Un-3WJ-3′-1 | None | CGCGCGGATC CAACCTACA | Specificity validation (Fig. 4G-H) |
| Un-3WJ-5′-1 | None | CTCAACCTT GATCCGCGCG | |
| Un-3WJ-3′-2 | None | CGCTTGTCGG CAACCTACA | |
| Un-3WJ-5′-2 | None | CTCAACCTT CCGACAAGCG | |
| Un-3WJ-3′-3 | None | GAACGGATAC CAACCTACA |  |
| Un-3WJ-5′-3 | None | CTCAACCTT GTATCCGTTC | |
| Un-3WJ-3′-4 | None | GCCTCGATTA CAACCTACA | |
| Un-3WJ-5′-4 | None | CTCAACCTT TAATCGAGGC | |
| Un-3WJ-3′-5 | None | CGACGGATGT CAACCTACA | |
| Un-3WJ-5′-5 | None | CTCAACCTT ACATCCGTCG | |
| Un-3WJ-3′-6 | None | AATTCGGCCG CAACCTACA | |
| Un-3WJ-5′-6 | None | CTCAACCTT CGGCCGAATT | |

| | | | |
|---|---|---|---|
| Un-3WJ-3′-7 | None | GCCCGTATTC CAACCTACA | |
| Un-3WJ-5′-7 | None | CTCAACCTT GAATACGGGC | |
| Un-3WJ-3′-8 | None | CCGCTTGCGA CAACCTACA | |
| Un-3WJ-5′-8 | None | CTCAACCTT TCGCAAGCGG | |
| Un-3WJ-3′-9 | None | GTAGGGCAAT CAACCTACA | |
| Un-3WJ-5′-9 | None | CTCAACCTT ATTGCCCTAC | |
| Un-3WJ-3′-10 | None | GCGGGAAGCA CAACCTACA | |
| Un-3WJ-5′-10 | None | CTCAACCTT TGCTTCCCGC | |
| Un-3WJ-3′-11 | None | CGTGGATTAG CAACCTACA | |
| Un-3WJ-5′-11 | None | CTCAACCTT CTAATCCACG | |
| Un-3WJ-3′-12 | None | GGCATCGACC CAACCTACA | |
| Un-3WJ-5′-12 | None | CTCAACCTT GGTCGATGCC | |
| Un-3WJ-3′-13 | None | AGGGTCATCG CAACCTACA | |
| Un-3WJ-5′-13 | None | CTCAACCTT CGATGACCCT | |
| Un-3WJ-3′-14 | None | CCGTAAGGTG CAACCTACA | |
| Un-3WJ-5′-14 | None | CTCAACCTT CACCTTACGG | |
| Un-3WJ-3′-15 | None | GGACCGAGGG CAACCTACA | |
| Un-3WJ-5′-15 | None | CTCAACCTT CCCTCGGTCC | |
| Un-3WJ-3′-16 | None | TTGTGTATCC CAACCTACA | |
| Un-3WJ-5′-16 | None | CTCAACCTT GGATACACAA | |
| Un-3WJ-3′-17 | None | CGCCGGATCT CAACCTACA | |
| Un-3WJ-5′-17 | None | CTCAACCTT AGATCCGGCG | |
| Un-3WJ-3′-18 | None | TTGAATCAAC CAACCTACA | |
| Un-3WJ-5′-18 | None | CTCAACCTT GTTGATTCAA | |
| Un-3WJ-3′-19 | None | TGCTAGTTCG CAACCTACA | |
| Un-3WJ-5′-19 | None | CTCAACCTT CGAACTAGCA | |
| Un-3WJ-3′-20 | None | ATCGCGCAAT CAACCTACA | |
| Un-3WJ-5′-20 | None | CTCAACCTT ATTGCGCGAT | |
| GAPDH-3WJ-N-3'-1 | *GAPDH* | ccaatacgaccaaatcagagaataatctag TTTGCTCGCA GGAGACACA | HBS optimization (Fig. S2B-D) |
| GAPDH-3WJ-N-5'-1 | *GAPDH* | GCATACTAT TGCGAGCAAA gaaaagcatcacccggaggagaaatcgggc | |
| GAPDH-3WJ-N-3'-2 | *GAPDH* | ggtcattgatggcaacaatatccactttac AGTGCGCACG GGAGACACA | |
| GAPDH-3WJ-N-5'-2 | *GAPDH* | GCATACTAT CGTGCGCACT cagagttaaaagcagccctggtgaccaggc | |
| GAPDH-3WJ-N-3'-3 | *GAPDH* | atggaatttgccatgggtggaatcatattg AGTTGAACTG GGAGACACA | |
| GAPDH-3WJ-N-5'-3 | *GAPDH* | GCATACTAT CAGTTCAACT gaacatgtaaaccatgtagttgaggtcaat | |
| GAPDH-3WJ-N-3'-4 | *GAPDH* | acttgattttggagggatctcgctcctgga AGTTTCCATA GGAGACACA | |
| GAPDH-3WJ-N-5'-4 | *GAPDH* | GCATACTAT TATGGAAACT agatggtgatgggatttccattgatgacaa | |
| GAPDH-3WJ-N-3'-5 | *GAPDH* | gcaggaggcattgctgatgatcttgaggct CGCGGCGTGA GGAGACACA | |
| GAPDH-3WJ-N-5'-5 | *GAPDH* | GCATACTAT TCACGCCGCG gttgtcatacttctcatggttcacacccat | |
| GAPDH-3WJ-N-3'-6 | *GAPDH* | tggcatggactgtggtcatgagtccttcca ATATCGCGGC GGAGACACA | |
| GAPDH-3WJ-N-5'-6 | *GAPDH* | GCATACTAT GCCGCGATAT cgataccaaagttgtcatggatgaccttgg | |
| GAPDH-3WJ-N-3'-7 | *GAPDH* | agggatgaccttgcccacagccttggcagc GCTACAGCGT GGAGACACA | |
| GAPDH-3WJ-N-5'-7 | *GAPDH* | GCATACTAT ACGCTGTAGC gccagtagaggcagggatgatgttctggag | |
| GAPDH-3WJ-N-3'-8 | *GAPDH* | ctccgacgcctgcttcaccaccttcttgat TACGGGCCCA GGAGACACA | |

| | | | |
|---|---|---|---|
| GAPDH-3WJ-N-5'-8 | *GAPDH* | GCATACTAT TGGGCCCGTA gtcatcatatttggcaggtttttctagacg | |
| GAPDH-3WJ-N-3'-9 | *GAPDH* | cagcgtcaaaggtggaggagtgggtgtcgc GACGTTTCAT GGAGACACA | |
| GAPDH-3WJ-N-5'-9 | *GAPDH* | GCATACTAT ATGAAACGTC tgttgaagtcagaggagaccacctggtgct | |
| GAPDH-3WJ-N-3'-10 | *GAPDH* | gttgctgtagccaaattcgttgtcatacca TTTATCGTGC GGAGACACA | |
| GAPDH-3WJ-N-5'-10 | *GAPDH* | GCATACTAT GCACGATAAA ggaaatgagcttgacaaagtggtcgttgag | |
| GAPDH-3WJ-3'-1 | *GAPDH* | ccaatacgaccaaatcagagaataatctag TTTGCTCGCA CAACCTACA | HBS optimization (Fig. S2B-D); Multiplex 3WJ-FISH (Fig. S13A) |
| GAPDH-3WJ-5'-1 | *GAPDH* | CTCAACCTT TGCGAGCAAA gaaaagcatcacccggaggagaaatcgggc | |
| GAPDH-3WJ-3'-2 | *GAPDH* | ggtcattgatggcaacaatatccactttac AGTGCGCACG CAACCTACA | |
| GAPDH-3WJ-5'-2 | *GAPDH* | CTCAACCTT CGTGCGCACT cagagttaaaagcagccctggtgaccaggc | |
| GAPDH-3WJ-3'-3 | *GAPDH* | atggaatttgccatgggtggaatcatattg AGTTGAACTG CAACCTACA | |
| GAPDH-3WJ-5'-3 | *GAPDH* | CTCAACCTT CAGTTCAACT gaacatgtaaaccatgtagttgaggtcaat | |
| GAPDH-3WJ-3'-4 | *GAPDH* | acttgattttggagggatctcgctcctgga AGTTTCCATA CAACCTACA | |
| GAPDH-3WJ-5'-4 | *GAPDH* | CTCAACCTT TATGGAAACT agatggtgatgggatttccattgatgacaa | |
| GAPDH-3WJ-3'-5 | *GAPDH* | gcaggaggcattgctgatgatcttgaggct CGCGGCGTGA CAACCTACA | |
| GAPDH-3WJ-5'-5 | *GAPDH* | CTCAACCTT TCACGCCGCG gttgtcatacttctcatggttcacacccat | |
| GAPDH-3WJ-3'-6 | *GAPDH* | tggcatggactgtggtcatgagtccttcca ATATCGCGGC CAACCTACA | |
| GAPDH-3WJ-5'-6 | *GAPDH* | CTCAACCTT GCCGCGATAT cgataccaaagttgtcatggatgaccttgg | |
| GAPDH-3WJ-3'-7 | *GAPDH* | agggatgaccttgcccacagccttggcagc GCTACAGCGT CAACCTACA | |
| GAPDH-3WJ-5'-7 | *GAPDH* | CTCAACCTT ACGCTGTAGC gccagtagaggcagggatgatgttctggag | |
| GAPDH-3WJ-3'-8 | *GAPDH* | ctccgacgcctgcttcaccaccttcttgat TACGGGCCCA CAACCTACA | |
| GAPDH-3WJ-5'-8 | *GAPDH* | CTCAACCTT TGGGCCCGTA gtcatcatatttggcaggtttttctagacg | |
| GAPDH-3WJ-3'-9 | *GAPDH* | cagcgtcaaaggtggaggagtgggtgtcgc GACGTTTCAT CAACCTACA | |
| GAPDH-3WJ-5'-9 | *GAPDH* | CTCAACCTT ATGAAACGTC tgttgaagtcagaggagaccacctggtgct | |
| GAPDH-3WJ-3'-10 | *GAPDH* | gttgctgtagccaaattcgttgtcatacca TTTATCGTGC CAACCTACA | |
| GAPDH-3WJ-5'-10 | *GAPDH* | CTCAACCTT GCACGATAAA ggaaatgagcttgacaaagtggtcgttgag | |
| UBC-3WJ-3'-1 | *UBC* | cctcaagcgcaggaccaagt ACGCGCCTTT CAACCTACA | Multiplex 3WJ-FISH (Fig. 1D(iii), S13B); Amplifier assembly optimization (Fig. 2E-G); Sensitivity validation (Fig. 3A-C); 3WJ-FISH vs smFISH (Fig. S9) |
| UBC-3WJ-5'-1 | *UBC* | CTCAACCTT AAAGGCGCGT gcagagtggactctttctgg | |
| UBC-3WJ-3'-2 | *UBC* | ggtgcgtccatcttccagct TACTTAATCG CAACCTACA | |
| UBC-3WJ-5'-2 | *UBC* | CTCAACCTT CGATTAAGTA gtttcccagcaaagatcaac | |
| UBC-3WJ-3'-3 | *UBC* | ctgctgatcaggagggatgc CTTTGTTCGG CAACCTACA | |
| UBC-3WJ-5'-3 | *UBC* | CTCAACCTT CCGAACAAAG cttccttatcttggatcttt | |
| UBC-3WJ-3'-4 | *UBC* | atggtcttaccagtcagggt CTCGGAGCGT CAACCTACA | |
| UBC-3WJ-5'-4 | *UBC* | CTCAACCTT ACGCTCCGAG cttcacgaagatttgcatcc | |
| UBC-3WJ-3'-5 | *UBC* | ccttgtcttggatctttgcc TACTGCGGGC CAACCTACA | |
| UBC-3WJ-5'-5 | *UBC* | CTCAACCTT GCCCGCAGTA ttgacattctcaatggtgtc | |
| UBC-3WJ-3'-6 | *UBC* | ctcggctccacttcgagagt TAATTCCGGT CAACCTACA | |
| UBC-3WJ-5'-6 | *UBC* | CTCAACCTT ACCGGAATTA gatggtcttaccagtcaggg | |
| UBC-3WJ-3'-7 | *UBC* | ctcggctccacctcgagagt ATCGGGCCCT CAACCTACA | |

| | | | |
|---|---|---|---|
| UBC-3WJ-5'-7 | *UBC* | CTCAACCTT AGGGCCCGAT gatggtcttaccagtcaggg | RNA-seq correlation<br>(Fig. 3D); (Fig. S7)<br><br>Additional expression<br>correlation dataset<br>(Fig. S12K-L) |
| UBC-3WJ-3'-8 | *UBC* | cttcacgaagatctgcatcc TTTGTCTGAC CAACCTACA | |
| UBC-3WJ-5'-8 | *UBC* | CTCAACCTT GTCAGACAAA cacctctgagacggagcacc | |
| UBC-3WJ-3'-9 | *UBC* | acagggtacgaccatcttcc CGTGAGAATG CAACCTACA | |
| UBC-3WJ-5'-9 | *UBC* | CTCAACCTT CATTCTCACG agctgttttccggcaaagat | |
| UBC-3WJ-3'-10 | *UBC* | cagggtagactctttctgga CGTCGCGATA CAACCTACA | |
| UBC-3WJ-5'-10 | *UBC* | CTCAACCTT TATCGCGACG tgttgtagtcagacagggtg | |
| UBC-3WJ-3'-11 | *UBC* | gctttccggcaaagatcaac AGCCGTATCA CAACCTACA | |
| UBC-3WJ-5'-11 | *UBC* | CTCAACCTT TGATACGGCT ctctgctggtcaggaggaat | |
| UBC-3WJ-3'-12 | *UBC* | cttccttgtcttggatcttt GTTTCAAGAG CAACCTACA | |
| UBC-3WJ-5'-12 | *UBC* | CTCAACCTT CTCTTGAAAC gctttgacgttctcgatagt | |
| UBC-3WJ-3'-13 | *UBC* | gggctcgacctcaagggtga TCCGTCGCAA CAACCTACA | |
| UBC-3WJ-5'-13 | *UBC* | CTCAACCTT TTGCGACGGA tggtcttgccagtgagtgtc | |
| UBC-3WJ-3'-14 | *UBC* | cacgaagatttgcatcccac ATTCCACTAC CAACCTACA | |
| UBC-3WJ-5'-14 | *UBC* | CTCAACCTT GTAGTGGAAT ctctgagacggagtaccagg | |
| UBC-3WJ-3'-15 | *UBC* | ctctgctgatcaggaggaat ACGCTACGCC CAACCTACA | |
| UBC-3WJ-5'-15 | *UBC* | CTCAACCTT GGCGTAGCGT gccttccttatcttggatct | |
| UBC-3WJ-3'-16 | *UBC* | tccttgtcctggatctttgc AGTGTTATCC CAACCTACA | |
| UBC-3WJ-5'-16 | *UBC* | CTCAACCTT GGATAACACT tttgacgttctcgatggtgt | |
| UBC-3WJ-3'-17 | *UBC* | cacgaagatttgcatcccac CCTTAATGCG CAACCTACA | |
| UBC-3WJ-5'-17 | *UBC* | CTCAACCTT CGCATTAAGG ctctgagacggagcaccagg | |
| UBC-3WJ-3'-18 | *UBC* | atggtcttaccagtcagagt CTGTCTTCGC CAACCTACA | |
| UBC-3WJ-5'-18 | *UBC* | CTCAACCTT GCGAAGACAG cttcacgaagatctgcattg | |
| UBC-3WJ-3'-19 | *UBC* | agcgatccacaaacaagaac CGAGATCACC CAACCTACA | |
| UBC-3WJ-5'-19 | *UBC* | CTCAACCTT GGTGATCTCG tgcgacccaaatcccggctg | |
| TOP2A-3WJ-3'-1 | *TOP2A* | atggaagtcatcactctccc ACATTCTCCC caaccTAcA | RNA-seq correlation<br>(Fig. 3D);<br><br>Additional expression<br>correlation dataset<br>(Fig. S7); (Fig. S12C-J);<br><br>3WJ-FISH vs smFISH<br>(Fig. S10, S11) |
| TOP2A-3WJ-5'-1 | *TOP2A* | ctcaaccTt GGGAGAATGT ccttggatttcttgcttgtg | |
| TOP2A-3WJ-3'-2 | *TOP2A* | aggatcaggcttttgagaga AGCCGTTACA caaccTAcA | |
| TOP2A-3WJ-5'-2 | *TOP2A* | ctcaaccTt TGTAACGGCT caccagaattcaaagctgga | |
| TOP2A-3WJ-3'-3 | *TOP2A* | cgtcactgctcctatctgat TAATACCTTC caaccTAcA | |
| TOP2A-3WJ-5'-3 | *TOP2A* | ctcaaccTt GAAGGTATTA tctgaatcagaccagggatt | |
| TOP2A-3WJ-3'-4 | *TOP2A* | gttattcgtggaatgactct CGCACGCTGT caaccTAcA | |
| TOP2A-3WJ-5'-4 | *TOP2A* | ctcaaccTt ACAGCGTGCG ttgaccacgcggagaaggca | |
| TOP2A-3WJ-3'-5 | *TOP2A* | aggccttcacaggatccgaa TATGCAGAGT caaccTAcA | |
| TOP2A-3WJ-5'-5 | *TOP2A* | ctcaaccTt ACTCTGCATA tcatatcccctctgaatcag | |
| TOP2A-3WJ-3'-6 | *TOP2A* | cgagcctgattattcagttt CGTGTATTTC caaccTAcA | |
| TOP2A-3WJ-5'-6 | *TOP2A* | ctcaaccTt GAAATACACG agcagattcagcaccaagca | |
| TOP2A-3WJ-3'-7 | *TOP2A* | tatggtattccctatagtct CTTTGCGCAC caaccTAcA | |
| TOP2A-3WJ-5'-7 | *TOP2A* | ctcaaccTt GTGCGCAAAG gttatgagaggaggtgtctt | |
| TOP2A-3WJ-3'-8 | *TOP2A* | gttcttcaatagtacccttg GTACTCAATT caaccTAcA | |
| TOP2A-3WJ-5'-8 | *TOP2A* | ctcaaccTt AATTGAGTAC aagttcttgtaacttggaag | |
| TOP2A-3WJ-3'-9 | *TOP2A* | ggttcttctccatccatcaa TGTAGTAGTG caaccTAcA | |
| TOP2A-3WJ-5'-9 | *TOP2A* | ctcaaccTt CACTACTACA acgcctgatgttatttacaa | |
| TOP2A-3WJ-3'-10 | *TOP2A* | gtaccattcaggctcaacac TTTCCGGCCG caaccTAcA | |
| TOP2A-3WJ-5'-10 | *TOP2A* | ctcaaccTt CGGCCGGAAA gctggttgtcatcatataaa | |
| TOP2A-3WJ-3'-11 | *TOP2A* | tgtaaagatgtatcgtggac CCATAGAATT caaccTAcA | |
| TOP2A-3WJ-5'-11 | *TOP2A* | ctcaaccTt AATTCTATGG tagcagaatccttgccacca | |
| TOP2A-3WJ-3'-12 | *TOP2A* | ctgatccagctaattgggca CACCGCGCTG caaccTAcA | |
| TOP2A-3WJ-5'-12 | *TOP2A* | ctcaaccTt CAGCGCGGTG acctttacttctcgcttgtc | |
| TOP2A-3WJ-3'-13 | *TOP2A* | ctccatgaaattagttaacc CCACACACCG caaccTAcA | |
| TOP2A-3WJ-5'-13 | *TOP2A* | ctcaaccTt CGGTGTGTGG attcctttcgatcatctatc | |
| TOP2A-3WJ-3'-14 | *TOP2A* | gagtgataaattcctccaga ATATTACTTA caaccTAcA | |
| TOP2A-3WJ-5'-14 | *TOP2A* | ctcaaccTt TAAGTAATAT aaacgatgtcgcagaagaga | |

| | | |
|---|---|---|
| TOP2A-3WJ-3'-15 | *TOP2A* | tgggaaccatcttggtcctg CCGCACGGAC caaccTAcA |
| TOP2A-3WJ-5'-15 | *TOP2A* | ctcaaccTt GTCCGTGCGG atctgtcataatcattatct |
| TOP2A-3WJ-3'-16 | *TOP2A* | atagtttttcttgtactgaa ATAACGCAAA caaccTAcA |
| TOP2A-3WJ-5'-16 | *TOP2A* | ctcaaccTt TTTGCGTTAT gacccacaatcttgatgata |
| TOP2A-3WJ-3'-17 | *TOP2A* | ctaagagggaaaaccccata AGATTCCATG caaccTAcA |
| TOP2A-3WJ-5'-17 | *TOP2A* | ctcaaccTt CATGGAATCT tttgtctctcccaaccacac |
| TOP2A-3WJ-3'-18 | *TOP2A* | tccctcagtcaggataagcg TCGTTCGTTG caaccTAcA |
| TOP2A-3WJ-5'-18 | *TOP2A* | ctcaaccTt CAACGAACGA tacactcagtggagtttcgg |
| TOP2A-3WJ-3'-19 | *TOP2A* | catcattggcatcatcgagt CTTTCGGGTC caaccTAcA |
| TOP2A-3WJ-5'-19 | *TOP2A* | ctcaaccTt GACCCGAAAG ttgggaattcccttgattct |
| TOP2A-3WJ-3'-20 | *TOP2A* | ttgtttaactggacttgggc ATGTTCCGGG caaccTAcA |
| TOP2A-3WJ-5'-20 | *TOP2A* | ctcaaccTt CCCGGAACAT cttaaacttcacccagttta |
| TOP2A-3WJ-3'-21 | *TOP2A* | caattggcatgttgatccaa AGTTAAGTGC caaccTAcA |
| TOP2A-3WJ-5'-21 | *TOP2A* | ctcaaccTt GCACTTAACT agctcttgggttgtaaagtc |
| TOP2A-3WJ-3'-22 | *TOP2A* | actacgaaatccttttactg CCTAGTGATT caaccTAcA |
| TOP2A-3WJ-5'-22 | *TOP2A* | ctcaaccTt AATCACTAGG gcagtttatttccattaaga |
| TOP2A-3WJ-3'-23 | *TOP2A* | tggatccagcaatatcatat TTCCCGGACA caaccTAcA |
| TOP2A-3WJ-5'-23 | *TOP2A* | ctcaaccTt TGTCCGGGAA gctcttctgaccattagtgc |
| TOP2A-3WJ-3'-24 | *TOP2A* | catagttactagaagttagg AGACCATACG caaccTAcA |
| TOP2A-3WJ-5'-24 | *TOP2A* | ctcaaccTt CGTATGGTCT agctgtccaaatatgagagc |
| TOP2A-3WJ-3'-25 | *TOP2A* | acaacaggaatacctttcc CTTTGTATCC caaccTAcA |
| TOP2A-3WJ-5'-25 | *TOP2A* | ctcaaccTt GGATACAAAG attattccatatactaatta |
| TOP2A-3WJ-3'-26 | *TOP2A* | aatgccaacatcttcatcgt CCGCATTGTC caaccTAcA |
| TOP2A-3WJ-5'-26 | *TOP2A* | ctcaaccTt GACAATGCGG aaacccacatttgctgggtc |
| TOP2A-3WJ-3'-27 | *TOP2A* | accaatgtaggtgtctgggc AACCGTGATA caaccTAcA |
| TOP2A-3WJ-5'-27 | *TOP2A* | ctcaaccTt TATCACGGTT ggagcaaaatatgttccaat |
| TOP2A-3WJ-3'-28 | *TOP2A* | tacacaatttggctccatag AACGCCCCAC caaccTAcA |
| TOP2A-3WJ-5'-28 | *TOP2A* | ctcaaccTt GTGGGGCGTT ccatttcgaccacctgtcac |
| TOP2A-3WJ-3'-29 | *TOP2A* | gttccatctcaccagctctt AACCGGTCTA caaccTAcA |
| TOP2A-3WJ-5'-29 | *TOP2A* | ctcaaccTt TAGACCGGTT cccatattatccatccatgt |
| TOP2A-3WJ-3'-30 | *TOP2A* | acatcaacaagtttagtcac TTCTGCCCCG caaccTAcA |
| TOP2A-3WJ-5'-30 | *TOP2A* | ctcaaccTt CGGGGCAGAA aatctgatcagctacataat |
| TOP2A-3WJ-3'-31 | *TOP2A* | ccatgatctgcttatgagaa GATTGTGTAG caaccTAcA |
| TOP2A-3WJ-5'-31 | *TOP2A* | ctcaaccTt CTACACAATC gcttctcgaacattgagtat |
| TOP2A-3WJ-3'-32 | *TOP2A* | ctcttccactcttcaaattc CTTAGTTCGA caaccTAcA |
| TOP2A-3WJ-5'-32 | *TOP2A* | ctcaaccTt TCGAACTAAG aggaaggctgtaaaatgcca |
| TOP2A-3WJ-3'-33 | *TOP2A* | gtattctttagcttcctttg CCGTCTCAAG caaccTAcA |
| TOP2A-3WJ-5'-33 | *TOP2A* | ctcaaccTt CTTGAGACGG atgtgctggtgcccaaacct |
| TOP2A-3WJ-3'-34 | *TOP2A* | catcatcttcaggaccagaa CTCTCACCGG caaccTAcA |
| TOP2A-3WJ-5'-34 | *TOP2A* | ctcaaccTt CCGGTGAGAG tatttgaactggatacgatg |
| TOP2A-3WJ-3'-35 | *TOP2A* | tgtcagatatgtggtagttt GTATGGGCCT caaccTAcA |
| TOP2A-3WJ-5'-35 | *TOP2A* | ctcaaccTt AGGCCCATAC gtccatacaagtaatcctca |
| TOP2A-3WJ-3'-36 | *TOP2A* | atctctcgttatcagaattt GGTTACTCGT caaccTAcA |
| TOP2A-3WJ-5'-36 | *TOP2A* | ctcaaccTt ACGAGTAACC gagaacaagataagttcctt |
| TOP2A-3WJ-3'-37 | *TOP2A* | ttctctgacctggtttcaaa CGATAGTGAT caaccTAcA |
| TOP2A-3WJ-5'-37 | *TOP2A* | ctcaaccTt ATCACTATCG ccatccaccatagaagggat |
| TOP2A-3WJ-3'-38 | *TOP2A* | ggtcatcattagtgacatct TATCGCCCAG caaccTAcA |
| TOP2A-3WJ-5'-38 | *TOP2A* | ctcaaccTt CTGGGCGATA caccatgatgataagaagac |
| TOP2A-3WJ-3'-39 | *TOP2A* | tgtgagactagtttggagtt GTTCTTCTAT caaccTAcA |
| TOP2A-3WJ-5'-39 | *TOP2A* | ctcaaccTt ATAGAAGAAC tgaagactttgtgtagtcca |
| TOP2A-3WJ-3'-40 | *TOP2A* | gctgctctccgtggctctgt CGGGCCTGAC caaccTAcA |
| TOP2A-3WJ-5'-40 | *TOP2A* | ctcaaccTt GTCAGGCCCG ttctcgtggagggacatcaa |
| TOP2A-3WJ-3'-41 | *TOP2A* | tgcccttaacatcatcagct AATTCGGTAT caaccTAcA |

| | | | |
|---|---|---|---|
| TOP2A-3WJ-5'-41 | *TOP2A* | ctcaaccTt ATACCGAATT tcaaggtctgacacgacact | |
| TOP2A-3WJ-3'-42 | *TOP2A* | aggtttctttgcccgtacag ATTGTTGCAC caaccTAcA | |
| TOP2A-3WJ-5'-42 | *TOP2A* | ctcaaccTt GTGCAACAAT attttgcccgaggagccaca | |
| TOP2A-3WJ-3'-43 | *TOP2A* | cagtcttgggcttggtaaga CCTGCTCAAC caaccTAcA | |
| TOP2A-3WJ-5'-43 | *TOP2A* | ctcaaccTt GTTGAGCAGG taattacttaaaataatcgc | |
| TOP2A-3WJ-3'-44 | *TOP2A* | acaatgcccatgagatggtc AACTCGTCAC caaccTAcA | |
| TOP2A-3WJ-5'-44 | *TOP2A* | ctcaaccTt GTGACGAGTT actatttagacagtattata | |
| TOP2A-3WJ-3'-45 | *TOP2A* | gaagagtgacacttttaaat AATAGCTCTC caaccTAcA | |
| TOP2A-3WJ-5'-45 | *TOP2A* | ctcaaccTt GAGAGCTATT caaactgctctagttttagc | |
| TOP2A-3WJ-3'-46 | *TOP2A* | ctaagataaaacagataatt CGCGCTTATA caaccTAcA | |
| TOP2A-3WJ-5'-46 | *TOP2A* | ctcaaccTt TATAAGCGCG atgctctatctcatatctac | |
| TOP2A-3WJ-3'-47 | *TOP2A* | agtatctgtactagaaccat TGAATCCCCA caaccTAcA | |
| TOP2A-3WJ-5'-47 | *TOP2A* | ctcaaccTt TGGGGATTCA aaagttctatctgatggtaa | |
| TOP2A-3WJ-3'-48 | *TOP2A* | cggcctctgatgatttgaga TATAAGCGGC caaccTAcA | |
| TOP2A-3WJ-5'-48 | *TOP2A* | ctcaaccTt GCCGCTTATA agatgaagaacatgagcaat | |
| TOP2A-3WJ-3'-49 | *TOP2A* | ctctaattttcttcattcta CTCTCATCAT caaccTAcA | |
| TOP2A-3WJ-5'-49 | *TOP2A* | ctcaaccTt ATGATGAGAG ttgactgtgtcaagttatag | |
| TOP2A-3WJ-3'-50 | *TOP2A* | catttaagatcttggatcaa GTCGACCGTC caaccTAcA | |
| TOP2A-3WJ-5'-50 | *TOP2A* | ctcaaccTt GACGGTCGAC atgttgtccccgagtcttct | |
| EPIC1-3WJ-3'-1 | *EPIC1* | gagagaaagcgggcgaggca CAAGTATGCA CAACCTACA | RNA-seq correlation (Fig. 3D);<br><br>Additional expression correlation dataset (Fig. S12A-B)<br><br>3WJ-FISH vs smFISH (Fig. S8) |
| EPIC1-3WJ-5'-1 | *EPIC1* | CTCAACCTT TGCATACTTG gcaggagctctgagggataa | |
| EPIC1-3WJ-3'-2 | *EPIC1* | aggcccacctcactttcagc GCATGAGTTG CAACCTACA | |
| EPIC1-3WJ-5'-2 | *EPIC1* | CTCAACCTT CAACTCATGC tgaggccacacttcacgttt | |
| EPIC1-3WJ-3'-3 | *EPIC1* | gccagcaggctggaaattcc GCCTCGATTA CAACCTACA | |
| EPIC1-3WJ-5'-3 | *EPIC1* | CTCAACCTT TAATCGAGGC ggcaggagtcgacgctacag | |
| EPIC1-3WJ-3'-4 | *EPIC1* | tgagcccttagggttcctcc TTGTGTATCC CAACCTACA | |
| EPIC1-3WJ-5'-4 | *EPIC1* | CTCAACCTT GGATACACAA gcctcagctgcgattgtgga | |
| EPIC1-3WJ-3'-5 | *EPIC1* | ggtgctccatcaatgcatag TGCTAGTTCG CAACCTACA | |
| EPIC1-3WJ-5'-5 | *EPIC1* | CTCAACCTT CGAACTAGCA aagggcaaatccatgatctc | |
| EPIC1-3WJ-3'-6 | *EPIC1* | tcctcagcatcccagcacca TGTTTACGAC CAACCTACA | |
| EPIC1-3WJ-5'-6 | *EPIC1* | CTCAACCTT GTCGTAAACA aatacagacgctgtgggcag | |
| EPIC1-3WJ-3'-7 | *EPIC1* | ctgtttgcttagagtggcag GGGAAACGGT CAACCTACA | |
| EPIC1-3WJ-5'-7 | *EPIC1* | CTCAACCTT ACCGTTTCCC cctctgtttgaagatctcag | |
| EPIC1-3WJ-3'-8 | *EPIC1* | aaggagaactgggcccaagc GATGGTCGAC CAACCTACA | |
| EPIC1-3WJ-5'-8 | *EPIC1* | CTCAACCTT GTCGACCATC ttcagagtccagggctcggg | |
| EPIC1-3WJ-3'-9 | *EPIC1* | aagggccccctcacttcaca ATGATCCACG CAACCTACA | |
| EPIC1-3WJ-5'-9 | *EPIC1* | CTCAACCTT CGTGGATCAT gtgggatctgaaacccggag | |
| EPIC1-3WJ-3'-10 | *EPIC1* | agaagggtccttgaaataga GAATTTGGCC CAACCTACA | |
| EPIC1-3WJ-5'-10 | *EPIC1* | CTCAACCTT GGCCAAATTC gaagaggcaccgatgctcat | |
| EPIC1-3WJ-3'-11 | *EPIC1* | tatcttgcagtggtgggaat CGTTCTATGT CAACCTACA | |
| EPIC1-3WJ-5'-11 | *EPIC1* | CTCAACCTT ACATAGAACG ccatgtaaggcatctcagag | |
| EPIC1-3WJ-3'-12 | *EPIC1* | ttcaattacgctctgctggt CTAACTCGTG CAACCTACA | |
| EPIC1-3WJ-5'-12 | *EPIC1* | CTCAACCTT CACGAGTTAG cataacactttacatacgat | |
| EPIC1-3WJ-3'-13 | *EPIC1* | tgaagcctcatctgagccgc TTCCCGCTTG CAACCTACA | |
| EPIC1-3WJ-5'-13 | *EPIC1* | CTCAACCTT CAAGCGGGAA tgtctttccctctggaatgc | |
| EPIC1-3WJ-3'-14 | *EPIC1* | gtagcagtgtctccggcagc GTCCTCGCTC CAACCTACA | |
| EPIC1-3WJ-5'-14 | *EPIC1* | CTCAACCTT GAGCGAGGAC cagccagtcagatttggtca | |
| EPIC1-3WJ-3'-15 | *EPIC1* | tcacatgatgatcaaatgag TTTCCGCATT CAACCTACA | |
| EPIC1-3WJ-5'-15 | *EPIC1* | CTCAACCTT AATGCGGAAA gccatttattaagggccgat | |
| EPIC1-3WJ-3'-16 | *EPIC1* | gcattttgtttggcaaggat TTCCCGTATG CAACCTACA | |
| EPIC1-3WJ-5'-16 | *EPIC1* | CTCAACCTT CATACGGGAA aatcaagcccagcttctgaa | |
| EPIC1-3WJ-3'-17 | *EPIC1* | atcaggggagacaggtaga CGCTTTGTAT CAACCTACA | |
| EPIC1-3WJ-5'-17 | *EPIC1* | CTCAACCTT ATACAAAGCG ctctgtcgcccaagggttta | |

| | | | |
|---|---|---|---|
| EPIC1-3WJ-3'-18 | *EPIC1* | catctttgtcagggctgctg TCCTGTCCCG CAACCTACA | |
| EPIC1-3WJ-5'-18 | *EPIC1* | CTCAACCTT CGGGACAGGA gcaatgaagatagcatcgaa | |
| EPIC1-3WJ-3'-19 | *EPIC1* | ggctctgaggtaagcaggtt ACGATCCTAC CAACCTACA | |
| EPIC1-3WJ-5'-19 | *EPIC1* | CTCAACCTT GTAGGATCGT cctgagaggctatccgtctg | |
| EPIC1-3WJ-3'-20 | *EPIC1* | gcctccacaaatggattttg CCTTAAAGTT CAACCTACA | |
| EPIC1-3WJ-5'-20 | *EPIC1* | CTCAACCTT AACTTTAAGG gagaagacaactcagtccgc | |
| EPIC1-3WJ-3'-21 | *EPIC1* | caaatgcttcatctccaaat ACTGCGCACC CAACCTACA | |
| EPIC1-3WJ-5'-21 | *EPIC1* | CTCAACCTT GGTGCGCAGT actctcccactggggatgag | |
| EPIC1-3WJ-3'-22 | *EPIC1* | atcctgaggctccactctga CGGTGAGGTA CAACCTACA | |
| EPIC1-3WJ-5'-22 | *EPIC1* | CTCAACCTT TACCTCACCG gctcatctaagcttaatcac | |
| EPIC1-3WJ-3'-23 | *EPIC1* | gagcaagggagggccaggtc GGAGTCGTGC CAACCTACA | |
| EPIC1-3WJ-5'-23 | *EPIC1* | CTCAACCTT GCACGACTCC tcagtgtccctctttaccag | |
| EPIC1-3WJ-3'-24 | *EPIC1* | tggcttgtaggtggccacca GTATCGTTTC CAACCTACA | |
| EPIC1-3WJ-5'-24 | *EPIC1* | CTCAACCTT GAAACGATAC tctcattgtgtcctcacacg | |
| EPIC1-3WJ-3'-25 | *EPIC1* | aggctggaacggccagtgtc CTATATAGCC CAACCTACA | |
| EPIC1-3WJ-5'-25 | *EPIC1* | CTCAACCTT GGCTATATAG aaggccccgtagggttcagt | |
| EPIC1-3WJ-3'-26 | *EPIC1* | cccagtgacttaaagagcag AGTTCCTGTA CAACCTACA | |
| EPIC1-3WJ-5'-26 | *EPIC1* | CTCAACCTT TACAGGAACT acatttattcctagccattc | |
| EPIC1-3WJ-3'-27 | *EPIC1* | caaaggctctcaaccagccc GTATCCCGTC CAACCTACA | |
| EPIC1-3WJ-5'-27 | *EPIC1* | CTCAACCTT GACGGGATAC ctggtgctgccataacaaag | |
| EPIC1-3WJ-3'-28 | *EPIC1* | ccaagagttagaaatctcat GCATAGTCGT CAACCTACA | |
| EPIC1-3WJ-5'-28 | *EPIC1* | CTCAACCTT ACGACTATGC tccaggtggaacaagagcca | |
| EPIC1-3WJ-3'-29 | *EPIC1* | gtgggtggttgcaacatgaa ACATTCACCG CAACCTACA | |
| EPIC1-3WJ-5'-29 | *EPIC1* | CTCAACCTT CGGTGAATGT acaatgaggtaagggggtga | |
| EPIC1-3WJ-3'-30 | *EPIC1* | ctaccacacgccaagcattt AACAATTCAG CAACCTACA | |
| EPIC1-3WJ-5'-30 | *EPIC1* | CTCAACCTT CTGAATTGTT tctgggtccagggaggcaga | |
| XIST-H1-3WJ-3′-1 | *XIST1* | cctgcggcaaaacccaacacgaaaagcaca TAGACGCACC CACTCTACT | Co-detection RNA–protein (Fig.1D (iv), Fig.6C-E, Fig.S18A-B, Fig. S19); Multiplex 3WJ-FISH (Fig. 1D(iii), S13B); |
| XIST-H1-3WJ-5′-1 | *XIST1* | CCTTCATTC GGTGCGTCTA cagcaaagacaaagaggcccgccatgtttt | |
| XIST-H1-3WJ-3′-2 | *XIST1* | ccgccatgatgtccacgtgacaaaagccat ACGCTCGACA CACTCTACT | |
| XIST-H1-3WJ-5′-2 | *XIST1* | CCTTCATTC ACGCTCGACA gatatacatatgacaacgcctgccatattg | |
| XIST-H1-3WJ-3′-3 | *XIST1* | aataagtctgaatagaggacaccagaccac AGAACAGGAC CACTCTACT | |
| XIST-H1-3WJ-5′-3 | *XIST1* | CCTTCATTC GTCCTGTTCT agcaattcaagtttcctgtctgataagtag | |
| XIST-H1-3WJ-3′-4 | *XIST1* | gcaatgacagagaaatggtttctcctttaa ACTCGTTTAA CACTCTACT | |
| XIST-H1-3WJ-5′-4 | *XIST1* | CCTTCATTC TTAAACGAGT gggcaggggatcgagaaaatgataagtaac | |
| XIST-H1-3WJ-3′-5 | *XIST1* | ccactgctgttatgcaaagatgtttgcaaa CGGCCGGTTG CACTCTACT | |
| XIST-H1-3WJ-5′-5 | *XIST1* | CCTTCATTC CAACCGGCCG ggccatagtgtaactaacagaactatggat | |
| XIST-H1-3WJ-3′-6 | *XIST1* | gggcacatcaactactgctaattatgcgca GTCTACGGAT CACTCTACT | |
| XIST-H1-3WJ-5′-6 | *XIST1* | CCTTCATTC ATCCGTAGAC tttaagcagtataagagaagaagcactagc | |
| XIST-H1-3WJ-3′-7 | *XIST1* | aatataagtatgcacattaacagtccaaga GGGTACTCGC CACTCTACT | |
| XIST-H1-3WJ-5′-7 | *XIST1* | CCTTCATTC GCGAGTACCC gagtgaattcaggctagttagaagctcctg | |
| XIST-H1-3WJ-3′-8 | *XIST1* | tattgattagcactctctgctttgataagt GTCTGGAACG CACTCTACT | |
| XIST-H1-3WJ-5′-8 | *XIST1* | CCTTCATTC CGTTCCAGAC acaggagtcctgataaaaggcacaactgta | |
| XIST-H1-3WJ-3′-9 | *XIST1* | agtctaattgaaggaccattgacaactgca ACGGATTCAG CACTCTACT | |
| XIST-H1-3WJ-5′-9 | *XIST1* | CCTTCATTC CTGAATCCGT attacatgccatctacagttcgaagagaag | |
| XIST-H1-3WJ-3′-10 | *XIST1* | ttattcacatggaatgagcagtgtgcgatt AACATCGGAT CACTCTACT | |
| XIST-H1-3WJ-5′-10 | *XIST1* | CCTTCATTC ATCCGATGTT acgcacataaatgtccaagatgcatgcttg | |

| | | |
|---|---|---|
| XIST-H1-3WJ-3′-11 | *XIST1* | gagagtaggattttattcagataggttgag CGGTGCGGTG CACTCTACT |
| XIST-H1-3WJ-5′-11 | *XIST1* | CCTTCATTC CACCGCACCG ctgtgtgtagttatgcaaatccatgtctaa |
| XIST-H1-3WJ-3′-12 | *XIST1* | ctgagagtaggaccttattcagatggaatg GGATGGATAA CACTCTACT |
| XIST-H1-3WJ-5′-12 | *XIST1* | CCTTCATTC TTATCCATCC ggcaaagtggttatgcacattaatgtccaa |
| XIST-H1-3WJ-3′-13 | *XIST1* | tccttgtctaatacacaggaacccagacaa AGACGACGCA CACTCTACT |
| XIST-H1-3WJ-5′-13 | *XIST1* | CCTTCATTC TGCGTCGTCT atataatcacgcacataacaggccaagaaa |
| XIST-H1-3WJ-3′-14 | *XIST1* | ggtcacttataattgtgcaccttgattgtc CGTTCGTACC CACTCTACT |
| XIST-H1-3WJ-5′-14 | *XIST1* | CCTTCATTC GGTACGAACG caaatgtaatgatcttatggagggggcact |
| XIST-H1-3WJ-3′-15 | *XIST1* | ggtctgagagtaaggccttattcagatgag GCGTACTTCG CACTCTACT |
| XIST-H1-3WJ-5′-15 | *XIST1* | CCTTCATTC CGAAGTACGC ataggccgtgtgcagttacacacattattg |

**Table S2. DNA sequences of gene-specific probes used for 3WJ-FISH experiments.**

Each probe consists of a target-recognition sequence (TRS; lowercase) fused to functional domains (uppercase), including half-binding sequences (HBS) and junction-forming sequences (JFS). Pairs of 3′ and 5′ probes hybridize to adjacent regions of the target sequence to enable three-way junction formation. All sequences are listed in the 5′ to 3′ direction. Probes were used for specificity, sensitivity, expression correlation, and multiplexed RNA–protein co-detection analyses. The HBS of XIST probes pair with Set 2 amplifiers (Set 2a), whereas those of all other probes pair with Set 1 amplifiers (Table S1).

| Name | Target | HBS length (nt) | Sequence (5'→3') | Application (Figures) |
|---|---|---|---|---|
| UBC-HBS5-3WJ-3′-1 | *UBC* | 5 | cctcaagcgcaggaccaagt ACGCGCCTTT GTGGT | HBS length optimization (Fig. 4A-F); |
| UBC-HBS5-3WJ-5′-1 | *UBC* | 5 | cctcaagcgcaggaccaagt ACGCGCCTTT GTGGT | |
| UBC-HBS5-3WJ-3′-2 | *UBC* | 5 | ggtgcgtccatcttccagct TACTTAATCG GTGGT | |
| UBC-HBS5-3WJ-5′-2 | *UBC* | 5 | ggtgcgtccatcttccagct TACTTAATCG GTGGT | |
| UBC-HBS5-3WJ-3′-3 | *UBC* | 5 | ctgctgatcaggagggatgc CTTTGTTCGG GTGGT | |
| UBC-HBS5-3WJ-5′-3 | *UBC* | 5 | ctgctgatcaggagggatgc CTTTGTTCGG GTGGT | |
| UBC-HBS5-3WJ-3′-4 | *UBC* | 5 | atggtcttaccagtcagggt CTCGGAGCGT GTGGT | |
| UBC-HBS5-3WJ-5′-4 | *UBC* | 5 | atggtcttaccagtcagggt CTCGGAGCGT GTGGT | |
| UBC-HBS5-3WJ-3′-5 | *UBC* | 5 | ccttgtcttggatctttgcc TACTGCGGGC GTGGT | |
| UBC-HBS5-3WJ-5′-5 | *UBC* | 5 | ccttgtcttggatctttgcc TACTGCGGGC GTGGT | |
| UBC-HBS5-3WJ-3′-6 | *UBC* | 5 | ctcggctccacttcgagagt TAATTCCGGT GTGGT | |
| UBC-HBS5-3WJ-5′-6 | *UBC* | 5 | ctcggctccacttcgagagt TAATTCCGGT GTGGT | |
| UBC-HBS5-3WJ-3′-7 | *UBC* | 5 | ctcggctccacctcgagagt ATCGGGCCCT GTGGT | |
| UBC-HBS5-3WJ-5′-7 | *UBC* | 5 | ctcggctccacctcgagagt ATCGGGCCCT GTGGT | |
| UBC-HBS5-3WJ-3′-8 | *UBC* | 5 | cttcacgaagatctgcatcc TTTGTCTGAC GTGGT | |
| UBC-HBS5-3WJ-5′-8 | *UBC* | 5 | cttcacgaagatctgcatcc TTTGTCTGAC GTGGT | |
| UBC-HBS5-3WJ-3′-9 | *UBC* | 5 | acagggtacgaccatcttcc CGTGAGAATG GTGGT | |
| UBC-HBS5-3WJ-5′-9 | *UBC* | 5 | acagggtacgaccatcttcc CGTGAGAATG GTGGT | |
| UBC-HBS5-3WJ-3′-10 | *UBC* | 5 | cagggtagactctttctgga CGTCGCGATA GTGGT | |
| UBC-HBS5-3WJ-5′-10 | *UBC* | 5 | cagggtagactctttctgga CGTCGCGATA GTGGT | |
| UBC-HBS5-3WJ-3′-11 | *UBC* | 5 | gctttccggcaaagatcaac AGCCGTATCA GTGGT | |
| UBC-HBS5-3WJ-5′-11 | *UBC* | 5 | gctttccggcaaagatcaac AGCCGTATCA GTGGT | |
| UBC-HBS5-3WJ-3′-12 | *UBC* | 5 | cttccttgtcttggatcttt GTTTCAAGAG GTGGT | |
| UBC-HBS5-3WJ-5′-12 | *UBC* | 5 | cttccttgtcttggatcttt GTTTCAAGAG GTGGT | |
| UBC-HBS5-3WJ-3′-13 | *UBC* | 5 | gggctcgacctcaagggtga TCCGTCGCAA GTGGT | |
| UBC-HBS5-3WJ-5′-13 | *UBC* | 5 | gggctcgacctcaagggtga TCCGTCGCAA GTGGT | |
| UBC-HBS5-3WJ-3′-14 | *UBC* | 5 | cacgaagatttgcatcccac ATTCCACTAC GTGGT | |
| UBC-HBS5-3WJ-5′-14 | *UBC* | 5 | cacgaagatttgcatcccac ATTCCACTAC GTGGT | |
| UBC-HBS5-3WJ-3′-15 | *UBC* | 5 | ctctgctgatcaggaggaat ACGCTACGCC GTGGT | |
| UBC-HBS5-3WJ-5′-15 | *UBC* | 5 | ctctgctgatcaggaggaat ACGCTACGCC GTGGT | |
| UBC-HBS5-3WJ-3′-16 | *UBC* | 5 | tccttgtcctggatctttgc AGTGTTATCC GTGGT | |
| UBC-HBS5-3WJ-5′-16 | *UBC* | 5 | tccttgtcctggatctttgc AGTGTTATCC GTGGT | |
| UBC-HBS5-3WJ-3′-17 | *UBC* | 5 | cacgaagatttgcatcccac CCTTAATGCG GTGGT | |
| UBC-HBS5-3WJ-5′-17 | *UBC* | 5 | cacgaagatttgcatcccac CCTTAATGCG GTGGT | |
| UBC-HBS5-3WJ-3′-18 | *UBC* | 5 | atggtcttaccagtcagagt CTGTCTTCGC GTGGT | |
| UBC-HBS5-3WJ-5′-18 | *UBC* | 5 | atggtcttaccagtcagagt CTGTCTTCGC GTGGT | |
| UBC-HBS5-3WJ-3′-19 | *UBC* | 5 | agcgatccacaaacaagaac CGAGATCACC GTGGT | |
| UBC-HBS5-3WJ-5′-19 | *UBC* | 5 | agcgatccacaaacaagaac CGAGATCACC GTGGT | |
| UBC-HBS9-3WJ-3′-1 | *UBC* | 9 | cctcaagcgcaggaccaagt ACGCGCCTTT GTGGTGGTG | HBS length optimization (Fig. 4A-F); |
| UBC-HBS9-3WJ-5′-1 | *UBC* | 9 | GTTGTTGTT AAAGGCGCGT gcagagtggactctttctgg | |
| UBC-HBS9-3WJ-3′-2 | *UBC* | 9 | ggtgcgtccatcttccagct TACTTAATCG GTGGTGGTG | |
| UBC-HBS9-3WJ-5′-2 | *UBC* | 9 | GTTGTTGTT CGATTAAGTA gtttcccagcaaagatcaac | |
| UBC-HBS9-3WJ-3′-3 | *UBC* | 9 | ctgctgatcaggagggatgc CTTTGTTCGG GTGGTGGTG | |
| UBC-HBS9-3WJ-5′-3 | *UBC* | 9 | GTTGTTGTT CCGAACAAAG cttccttatcttggatcttt | |
| UBC-HBS9-3WJ-3′-4 | *UBC* | 9 | atggtcttaccagtcagggt CTCGGAGCGT GTGGTGGTG | |
| UBC-HBS9-3WJ-5′-4 | *UBC* | 9 | GTTGTTGTT ACGCTCCGAG cttcacgaagatttgcatcc | |
| UBC-HBS9-3WJ-3′-5 | *UBC* | 9 | ccttgtcttggatctttgcc TACTGCGGGC GTGGTGGTG | |
| UBC-HBS9-3WJ-5′-5 | *UBC* | 9 | GTTGTTGTT GCCCGCAGTA ttgacattctcaatggtgtc | |
| UBC-HBS9-3WJ-3′-6 | *UBC* | 9 | ctcggctccacttcgagagt TAATTCCGGT GTGGTGGTG | |
| UBC-HBS9-3WJ-5′-6 | *UBC* | 9 | GTTGTTGTT ACCGGAATTA gatggtcttaccagtcaggg | |

| | | | | |
|---|---|---|---|---|
| UBC-HBS9-3WJ-3′-7 | *UBC* | 9 | ctcggctccacctcgagagt ATCGGGCCCT GTGGTGGTG | |
| UBC-HBS9-3WJ-5′-7 | *UBC* | 9 | GTTGTTGTT AGGGCCCGAT gatggtcttaccagtcaggg | |
| UBC-HBS9-3WJ-3′-8 | *UBC* | 9 | cttcacgaagatctgcatcc TTTGTCTGAC GTGGTGGTG | |
| UBC-HBS9-3WJ-5′-8 | *UBC* | 9 | GTTGTTGTT GTCAGACAAA cacctctgagacggagcacc | |
| UBC-HBS9-3WJ-3′-9 | *UBC* | 9 | acagggtacgaccatcttcc CGTGAGAATG GTGGTGGTG | |
| UBC-HBS9-3WJ-5′-9 | *UBC* | 9 | GTTGTTGTT CATTCTCACG agctgttttccggcaaagat | |
| UBC-HBS9-3WJ-3′-10 | *UBC* | 9 | cagggtagactctttctgga CGTCGCGATA GTGGTGGTG | |
| UBC-HBS9-3WJ-5′-10 | *UBC* | 9 | GTTGTTGTT TATCGCGACG tgttgtagtcagacagggtg | |
| UBC-HBS9-3WJ-3′-11 | *UBC* | 9 | gctttccggcaaagatcaac AGCCGTATCA GTGGTGGTG | |
| UBC-HBS9-3WJ-5′-11 | *UBC* | 9 | GTTGTTGTT TGATACGGCT ctctgctggtcaggaggaat | |
| UBC-HBS9-3WJ-3′-12 | *UBC* | 9 | cttccttgtcttggatcttt GTTTCAAGAG GTGGTGGTG | |
| UBC-HBS9-3WJ-5′-12 | *UBC* | 9 | GTTGTTGTT CTCTTGAAAC gctttgacgttctcgatagt | |
| UBC-HBS9-3WJ-3′-13 | *UBC* | 9 | gggctcgacctcaagggtga TCCGTCGCAA GTGGTGGTG | |
| UBC-HBS9-3WJ-5′-13 | *UBC* | 9 | GTTGTTGTT TTGCGACGGA tggtcttgccagtgagtgtc | |
| UBC-HBS9-3WJ-3′-14 | *UBC* | 9 | cacgaagatttgcatcccac ATTCCACTAC GTGGTGGTG | |
| UBC-HBS9-3WJ-5′-14 | *UBC* | 9 | GTTGTTGTT GTAGTGGAAT ctctgagacggagtaccagg | |
| UBC-HBS9-3WJ-3′-15 | *UBC* | 9 | ctctgctgatcaggaggaat ACGCTACGCC GTGGTGGTG | |
| UBC-HBS9-3WJ-5′-15 | *UBC* | 9 | GTTGTTGTT GGCGTAGCGT gccttccttatcttggatct | |
| UBC-HBS9-3WJ-3′-16 | *UBC* | 9 | tccttgtcctggatctttgc AGTGTTATCC GTGGTGGTG | |
| UBC-HBS9-3WJ-5′-16 | *UBC* | 9 | GTTGTTGTT GGATAACACT tttgacgttctcgatggtgt | |
| UBC-HBS9-3WJ-3′-17 | *UBC* | 9 | cacgaagatttgcatcccac CCTTAATGCG GTGGTGGTG | |
| UBC-HBS9-3WJ-5′-17 | *UBC* | 9 | GTTGTTGTT CGCATTAAGG ctctgagacggagcaccagg | |
| UBC-HBS9-3WJ-3′-18 | *UBC* | 9 | atggtcttaccagtcagagt CTGTCTTCGC GTGGTGGTG | |
| UBC-HBS9-3WJ-5′-18 | *UBC* | 9 | GTTGTTGTT GCGAAGACAG cttcacgaagatctgcattg | |
| UBC-HBS9-3WJ-3′-19 | *UBC* | 9 | agcgatccacaaacaagaac CGAGATCACC GTGGTGGTG | |
| UBC-HBS9-3WJ-5′-19 | *UBC* | 9 | GTTGTTGTT GGTGATCTCG tgcgacccaaatcccggctg | |
| UBC-HBS15-3WJ-5′-1 | *UBC* | 15 | GTTGTTGTTGTTGTT AAAGGCGCGT gcagagtggactctttctgg | HBS length optimization (Fig. 4A-F); |
| UBC-HBS15-3WJ-5′-2 | *UBC* | 15 | GTTGTTGTTGTTGTT CGATTAAGTA gtttcccagcaaagatcaac | |
| UBC-HBS15-3WJ-5′-3 | *UBC* | 15 | GTTGTTGTTGTTGTT CCGAACAAAG cttccttatcttggatcttt | |
| UBC-HBS15-3WJ-5′-4 | *UBC* | 15 | GTTGTTGTTGTTGTT ACGCTCCGAG cttcacgaagatttgcatcc | |
| UBC-HBS15-3WJ-5′-5 | *UBC* | 15 | GTTGTTGTTGTTGTT GCCCGCAGTA ttgacattctcaatggtgtc | |
| UBC-HBS15-3WJ-5′-6 | *UBC* | 15 | GTTGTTGTTGTTGTT ACCGGAATTA gatggtcttaccagtcaggg | |
| UBC-HBS15-3WJ-5′-7 | *UBC* | 15 | GTTGTTGTTGTTGTT AGGGCCCGAT gatggtcttaccagtcaggg | |
| UBC-HBS15-3WJ-5′-8 | *UBC* | 15 | GTTGTTGTTGTTGTT GTCAGACAAA cacctctgagacggagcacc | |
| UBC-HBS15-3WJ-5′-9 | *UBC* | 15 | GTTGTTGTTGTTGTT CATTCTCACG agctgttttccggcaaagat | |
| UBC-HBS15-3WJ-5′-10 | *UBC* | 15 | GTTGTTGTTGTTGTT TATCGCGACG tgttgtagtcagacagggtg | |
| UBC-HBS15-3WJ-5′-11 | *UBC* | 15 | GTTGTTGTTGTTGTT TGATACGGCT ctctgctggtcaggaggaat | |
| UBC-HBS15-3WJ-5′-12 | *UBC* | 15 | GTTGTTGTTGTTGTT CTCTTGAAAC gctttgacgttctcgatagt | |
| UBC-HBS15-3WJ-5′-13 | *UBC* | 15 | GTTGTTGTTGTTGTT TTGCGACGGA tggtcttgccagtgagtgtc | |
| UBC-HBS15-3WJ-5′-14 | *UBC* | 15 | GTTGTTGTTGTTGTT GTAGTGGAAT ctctgagacggagtaccagg | |

| | | | |
|---|---|---|---|
| UBC-HBS15-3WJ-5′-15 | *UBC* | 15 | GTTGTTGTTGTTGTT GGCGTAGCGT gccttccttatcttggatct |
| UBC-HBS15-3WJ-5′-16 | *UBC* | 15 | GTTGTTGTTGTTGTT GGATAACACT tttgacgttctcgatggtgt |
| UBC-HBS15-3WJ-5′-17 | *UBC* | 15 | GTTGTTGTTGTTGTT CGCATTAAGG ctctgagacggagcaccagg |
| UBC-HBS15-3WJ-5′-18 | *UBC* | 15 | GTTGTTGTTGTTGTT GCGAAGACAG cttcacgaagatctgcattg |
| UBC-HBS15-3WJ-5′-19 | *UBC* | 15 | GTTGTTGTTGTTGTT GGTGATCTCG tgcgacccaaatcccggctg |
| UBC-HBS20-3WJ-5′-1 | *UBC* | 20 | TTGTTGTTGTTGTTGTTGTT AAAGGCGCGT gcagagtggactctttctgg |
| UBC-HBS20-3WJ-5′-2 | *UBC* | 20 | TTGTTGTTGTTGTTGTTGTT CGATTAAGTA gtttcccagcaaagatcaac |
| UBC-HBS20-3WJ-5′-3 | *UBC* | 20 | TTGTTGTTGTTGTTGTTGTT CCGAACAAAG cttccttatcttggatcttt |
| UBC-HBS20-3WJ-5′-4 | *UBC* | 20 | TTGTTGTTGTTGTTGTTGTT ACGCTCCGAG cttcacgaagatttgcatcc |
| UBC-HBS20-3WJ-5′-5 | *UBC* | 20 | TTGTTGTTGTTGTTGTTGTT GCCCGCAGTA ttgacattctcaatggtgtc |
| UBC-HBS20-3WJ-5′-6 | *UBC* | 20 | TTGTTGTTGTTGTTGTTGTT ACCGGAATTA gatggtcttaccagtcaggg |
| UBC-HBS20-3WJ-5′-7 | *UBC* | 20 | TTGTTGTTGTTGTTGTTGTT AGGGCCCGAT gatggtcttaccagtcaggg |
| UBC-HBS20-3WJ-5′-8 | *UBC* | 20 | TTGTTGTTGTTGTTGTTGTT GTCAGACAAA cacctctgagacggagcacc |
| UBC-HBS20-3WJ-5′-9 | *UBC* | 20 | TTGTTGTTGTTGTTGTTGTT CATTCTCACG agctgttttccggcaaagat |
| UBC-HBS20-3WJ-5′-10 | *UBC* | 20 | TTGTTGTTGTTGTTGTTGTT TATCGCGACG tgttgtagtcagacagggtg |
| UBC-HBS20-3WJ-5′-11 | *UBC* | 20 | TTGTTGTTGTTGTTGTTGTT TGATACGGCT ctctgctggtcaggaggaat |
| UBC-HBS20-3WJ-5′-12 | *UBC* | 20 | TTGTTGTTGTTGTTGTTGTT CTCTTGAAAC gctttgacgttctcgatagt |
| UBC-HBS20-3WJ-5′-13 | *UBC* | 20 | TTGTTGTTGTTGTTGTTGTT TTGCGACGGA tggtcttgccagtgagtgtc |
| UBC-HBS20-3WJ-5′-14 | *UBC* | 20 | TTGTTGTTGTTGTTGTTGTT GTAGTGGAAT ctctgagacggagtaccagg |
| UBC-HBS20-3WJ-5′-15 | *UBC* | 20 | TTGTTGTTGTTGTTGTTGTT GGCGTAGCGT gccttccttatcttggatct |
| UBC-HBS20-3WJ-5′-16 | *UBC* | 20 | TTGTTGTTGTTGTTGTTGTT GGATAACACT tttgacgttctcgatggtgt |
| UBC-HBS20-3WJ-5′-17 | *UBC* | 20 | TTGTTGTTGTTGTTGTTGTT CGCATTAAGG ctctgagacggagcaccagg |
| UBC-HBS20-3WJ-5′-18 | *UBC* | 20 | TTGTTGTTGTTGTTGTTGTT GCGAAGACAG cttcacgaagatctgcattg |
| UBC-HBS20-3WJ-5′-19 | *UBC* | 20 | TTGTTGTTGTTGTTGTTGTT GGTGATCTCG tgcgacccaaatcccggctg |

**Table S3. DNA sequences of UBC probes used for HBS-length optimization (8x amplification).**

UBC-targeting split probes with varying HBS lengths (5, 9, 15, and 20 nt) were used to optimize probe specificity prior to amplifier hybridization (Figure 4). Short HBS designs (5 and 9 nt) function as 3WJ-dependent split probes that require cooperative assembly, whereas longer HBS designs (15 and 20 nt) serve as high-affinity controls to assess nonspecific probe retention in the absence of productive 3WJ formation. All experiments in this table were performed using 8× primary amplifier with an extended docking domain (P-Amp-1b-HBSopt) (Table S1), in contrast to the standard 16× amplification used in other 3WJ-FISH experiments. Target-recognition sequences are shown in lowercase, and functional domains (HBS and JFS) are shown in uppercase. All sequences are listed in the 5′ to 3′ direction.

| Name | Probe Architecture | HBS type | Sequence (5’→3’) | Application (Figures) |
|---|---|---|---|---|
| H3-3WJ-3' | 3WJ | H-HBS | gtgaaatcgc TTCCCGCTTG CAACCTACA | DNA origami<br>(Fig. 1D (i), Fig. 2A-C, Fig. 3E-H, Fig. S2A-D, Fig. S4);<br><br>3WJ-IF<br>(Fig.5 B-E, Fig. 5F, Fig. 6A-B, Fig. 6 C-E, Fig. 2E-F, Fig.S2 E-F, Fig. S3B-C, Fig. S15, Fig. S16, Fig. S17, Fig. S18A-B, Fig. S19A); |
| H3-3WJ-5' | 3WJ | H-HBS | CTCAACCTT CAAGCGGGAA agcagcccac | |
| H3-CIR-3' | Circular | H-HBS | CAACCTACA AAAAAAAAAA gtgaaatcgc | DNA origami<br>(Fig. S4A-D)<br>3WJ-FISH (Fig. S6)<br>3WJ-IF (Fig. S15) |
| H3-CIR-5' | Circular | H-HBS | agcagcccac AAAAAAAAAA CTCAACCTT | |
| H3-Pi-3' | π-shaped | H-HBS | gtgaaatcgc Aaccgcaa CAACCTACA | DNA origami<br>(Fig. S4A-D) |
| H3-Pi-5' | π-shaped | H-HBS | CTCAACCTT Aagcggaa gcagcccaca | |
| H3-4C-3' | 4C | H-HBS | gtgaaatcgc AAACCGCAAA CAACCTACA | DNA origami (Fig. 2A-C, Fig. S4A-D) |
| H3-4C-5' | 4C | H-HBS | CTCAACCTT AAAGCGGAAA agcagcccac | |
| H3-0C-3' | 0C | H-HBS | gtgaaatcgc AAAAAAAAAA CAACCTACA | DNA origami (Fig. 2A-C, Fig. S4A-D) |
| H3-0C-5' | 0C | H-HBS | CTCAACCTT AAAAAAAAAA agcagcccac | |
| H1-3WJ-3' | 3WJ | H-HBS | aacgagacgg GTCCTCGCTC CACTCTACT | 3WJ-IF<br>(Fig.5 F, S17A-B; Fig. S20) |
| H1 -3WJ-5' | 3WJ | H-HBS | CCTTCATTC GAGCGAGGAC tcaatcgcgc | |
| H3-3WJ-N-3' | 3WJ | N-HBS | gtgaaatcgc TTAGGTCCGG GGAGACACA | 3WJ-IF<br>(Fig. S2E-F) |
| H3-3WJ-N-5' | 3WJ | N-HBS | GCATACTAT CCGGACCTAA agcagcccac | |

**Table S4. DNA sequences of 3WJ probe architectures used for origami and antibody-based detection.**

Probe designs include 3WJ, π-shaped, 4C, 0C, and circular architecture. The 3WJ designs contain junction-forming (JFS) domains, whereas the other architectures lack JFS and instead form distinct structures through alternative strand connectivity. Uppercase letters denote functional domains (HBS and, where applicable, JFS), while lowercase letters denote target-binding and spacer regions. The same target-binding sequences were used across DNA origami and antibody-based detection systems. Sequences used for antibody–DNA conjugation are provided in Table S9. In a separate comparison (Figure S2), HBS variants were constructed using either four nucleotides (N-HBS) or three nucleotides (A/T/C; H-HBS). An additional H1 probe is included only for 3WJ-IF. The “Application” column indicates the experimental systems in which each probe was used: DNA origami (Origami), antibody-based detection (IF), and RNA detection (FISH).

| Name | Target | Full sequence (5’→3’) | Application (Figures) |
|---|---|---|---|
| NEAT1-Z-5′-1 | *NEAT1* | gcctattcctcctgactcctcc TTTTTTTTTT TTCCAACTC | Probe architecture comparison (Fig. S6) |
| NEAT1-Z-3′-1 | *NEAT1* | gatcatttccagggctgctgcg TTTTTTTTTT ACATCCAAC | |
| NEAT1-Z-5′-2 | *NEAT1* | aaaaaaggggctgctggcat TTTTTTTTTT TTCCAACTC | |
| NEAT1-Z-3′-2 | *NEAT1* | ttcacagcccttggtctgga TTTTTTTTTT ACATCCAAC | |
| NEAT1-Z-5′-3 | *NEAT1* | ccgcaccagtcaccggtgag TTTTTTTTTT TTCCAACTC | |
| NEAT1-Z-3′-3 | *NEAT1* | cctgtcaaacatgctaggtg TTTTTTTTTT ACATCCAAC | |
| NEAT1-Z-5′-4 | *NEAT1* | aagcgttggtcaatgttgtc TTTTTTTTTT TTCCAACTC | |
| NEAT1-Z-3′-4 | *NEAT1* | gcactgccacctggaaaata TTTTTTTTTT ACATCCAAC | |
| NEAT1-Z-5′-5 | *NEAT1* | tagcgccaaacctagagaaaag TTTTTTTTTT TTCCAACTC | |
| NEAT1-Z-3′-5 | *NEAT1* | gagtgagctcacaagaagagtt TTTTTTTTTT ACATCCAAC | |
| NEAT1-Z-5′-6 | *NEAT1* | aactttcagctggatggctgc TTTTTTTTTT TTCCAACTC | |
| NEAT1-Z-3′-6 | *NEAT1* | cggtccatgaagcatttttgt TTTTTTTTTT ACATCCAAC | |
| NEAT1-Z-5′-7 | *NEAT1* | gaacttcctcctcctaagcc TTTTTTTTTT TTCCAACTC | |
| NEAT1-Z-3′-7 | *NEAT1* | cctggaaacagaacattgga TTTTTTTTTT ACATCCAAC | |
| NEAT1-Z-5′-8 | *NEAT1* | atgcagcatctgaaaaccttt TTTTTTTTTT TTCCAACTC | |
| NEAT1-Z-3′-8 | *NEAT1* | accggaggctcaatttagaag TTTTTTTTTT ACATCCAAC | |
| NEAT1-Z-5′-9 | *NEAT1* | gcaaggttccaagcacaaaact TTTTTTTTTT TTCCAACTC | |
| NEAT1-Z-3′-9 | *NEAT1* | acagcttagggatcttcttgaa TTTTTTTTTT ACATCCAAC | |
| NEAT1-Z-5′-10 | *NEAT1* | ttgtggcatcaacgttaaaatgt TTTTTTTTTT TTCCAACTC | |
| NEAT1-Z-3′-10 | *NEAT1* | tacaaggcatcaatctgcg TTTTTTTTTT ACATCCAAC | |
| NEAT1-Z-5′-11 | *NEAT1* | ctaagaacttctccgagaaacg TTTTTTTTTT TTCCAACTC | |
| NEAT1-Z-3′-11 | *NEAT1* | cgccccaagttatttcatcagg TTTTTTTTTT ACATCCAAC | |
| NEAT1-Z-5′-12 | *NEAT1* | ctcccagcgtttagcacaacacaatg TTTTTTTTTT TTCCAACTC | |
| NEAT1-Z-3′-12 | *NEAT1* | aggaatgaccaacttgtacc TTTTTTTTTT ACATCCAAC | |
| NEAT1-Z-5′-13 | *NEAT1* | cacaatgcccaaactagacctg TTTTTTTTTT TTCCAACTC | |
| NEAT1-Z-3′-13 | *NEAT1* | tcctagtaatctgcaatgcaat TTTTTTTTTT ACATCCAAC | |
| NEAT1-Z-5′-14 | *NEAT1* | ctctgtgtcacctgttttcagtt TTTTTTTTTT TTCCAACTC | |
| NEAT1-Z-3′-14 | *NEAT1* | tttggttctcggaaaactggtga TTTTTTTTTT ACATCCAAC | |
| NEAT1-Z-5′-15 | *NEAT1* | caggaggaagctggtaaagacattt TTTTTTTTTT TTCCAACTC | |
| NEAT1-Z-3′-15 | *NEAT1* | ctctgaaacaggctgtcttggccac TTTTTTTTTT ACATCCAAC | |
| NEAT1-Z-5′-16 | *NEAT1* | ctcgtcttaagtggtccctta TTTTTTTTTT TTCCAACTC | |
| NEAT1-Z-3′-16 | *NEAT1* | atccagaagagcccatctaat TTTTTTTTTT ACATCCAAC | |
| NEAT1-Z-5′-17 | *NEAT1* | acacctgtgacaaatgaggaac TTTTTTTTTT TTCCAACTC | |
| NEAT1-Z-3′-17 | *NEAT1* | gctcatgattttcaatcacaag TTTTTTTTTT ACATCCAAC | |
| NEAT1-Z-5′-18 | *NEAT1* | tggttgttacaaagactacaat TTTTTTTTTT TTCCAACTC | |
| NEAT1-Z-3′-18 | *NEAT1* | acagtgaccacaaaaggttatt TTTTTTTTTT ACATCCAAC | |
| NEAT1-Z-5′-19 | *NEAT1* | tcatccccaagtcattggttaaga TTTTTTTTTT TTCCAACTC | |
| NEAT1-Z-3′-19 | *NEAT1* | tcccaacgacagtaattgtttgca TTTTTTTTTT ACATCCAAC | |
| NEAT1-Z-5′-20 | *NEAT1* | tccccatacatgcgtgactaat TTTTTTTTTT TTCCAACTC | |
| NEAT1-Z-3′-20 | *NEAT1* | caacagcatacccgagactact TTTTTTTTTT ACATCCAAC | |

**Table S5. DNA sequences of Z-FISH probes targeting *NEAT1*.**

Z-FISH probes were designed using the same target recognition sequences (TRS) as the corresponding *NEAT1* 3WJ-FISH probes, but with distinct functional domains to generate the Z-shaped architecture illustrated in Fig. S6A. Lowercase letters indicate target recognition sequences, whereas uppercase letters indicate functional domains. All sequences are listed in the 5′ to 3′ direction.

| **Name** | **Target** | **Full sequence (5’→3’)** | **Application (Figures)** |
|---|---|---|---|
| NEAT1-CIR-5′-1 | *NEAT1* | gcctattcctcctgactcctcc TTTTTTTTTTT ctcaaccTt | Probe architecture comparison (Fig. S6) |
| NEAT1-CIR-3′-1 | *NEAT1* | caaccTAcA TTTTTTTTTTT gatcatttccagggctgctgcg | |
| NEAT1-CIR-5′-2 | *NEAT1* | aaaaaaggggctgctggcat TTTTTTTTTTT ctcaaccTt | |
| NEAT1-CIR-3′-2 | *NEAT1* | caaccTAcA TTTTTTTTTTT ttcacagcccttggtctgga | |
| NEAT1-CIR-5′-3 | *NEAT1* | ccgcaccagtcaccggtgag TTTTTTTTTTT ctcaaccTt | |
| NEAT1-CIR-3′-3 | *NEAT1* | caaccTAcA TTTTTTTTTTT cctgtcaaacatgctaggtg | |
| NEAT1-CIR-5′-4 | *NEAT1* | aagcgttggtcaatgttgtc TTTTTTTTTTT ctcaaccTt | |
| NEAT1-CIR-3′-4 | *NEAT1* | caaccTAcA TTTTTTTTTTT gcactgccacctggaaaata | |
| NEAT1-CIR-5′-5 | *NEAT1* | tagcgccaaacctagagaaaag TTTTTTTTTTT ctcaaccTt | |
| NEAT1-CIR-3′-5 | *NEAT1* | caaccTAcA TTTTTTTTTTT gagtgagctcacaagaagagtt | |
| NEAT1-CIR-5′-6 | *NEAT1* | aactttcagctggatggctgc TTTTTTTTTTT ctcaaccTt | |
| NEAT1-CIR-3′-6 | *NEAT1* | caaccTAcA TTTTTTTTTTT cggtccatgaagcatttttgt | |
| NEAT1-CIR-5′-7 | *NEAT1* | gaacttcctcctcctaagcc TTTTTTTTTTT ctcaaccTt | |
| NEAT1-CIR-3′-7 | *NEAT1* | caaccTAcA TTTTTTTTTTT cctggaaacagaacattgga | |
| NEAT1-CIR-5′-8 | *NEAT1* | atgcagcatctgaaaaccttt TTTTTTTTTTT ctcaaccTt | |
| NEAT1-CIR-3′-8 | *NEAT1* | caaccTAcA TTTTTTTTTTT accggaggctcaatttagaag | |
| NEAT1-CIR-5′-9 | *NEAT1* | gcaaggttccaagcacaaaact TTTTTTTTTTT ctcaaccTt | |
| NEAT1-CIR-3′-9 | *NEAT1* | caaccTAcA TTTTTTTTTTT acagcttagggatcttcttgaa | |
| NEAT1-CIR-5′-10 | *NEAT1* | ttgtggcatcaacgttaaaatgt TTTTTTTTTTT ctcaaccTt | |
| NEAT1-CIR-3′-10 | *NEAT1* | caaccTAcA TTTTTTTTTTT tacaaggcatcaatctgcg | |
| NEAT1-CIR-5′-11 | *NEAT1* | ctaagaacttctccgagaaacg TTTTTTTTTTT ctcaaccTt | |
| NEAT1-CIR-3′-11 | *NEAT1* | caaccTAcA TTTTTTTTTTT cgccccaagttatttcatcagg | |
| NEAT1-CIR-5′-12 | *NEAT1* | ctcccagcgtttagcacaacacaatg TTTTTTTTTTT ctcaaccTt | |
| NEAT1-CIR-3′-12 | *NEAT1* | caaccTAcA TTTTTTTTTTT aggaatgaccaacttgtacc | |
| NEAT1-CIR-5′-13 | *NEAT1* | cacaatgcccaaactagacctg TTTTTTTTTTT ctcaaccTt | |
| NEAT1-CIR-3′-13 | *NEAT1* | caaccTAcA TTTTTTTTTTT tcctagtaatctgcaatgcaat | |
| NEAT1-CIR-5′-14 | *NEAT1* | ctctgtgtcacctgttttcagtt TTTTTTTTTTT ctcaaccTt | |
| NEAT1-CIR-3′-14 | *NEAT1* | caaccTAcA TTTTTTTTTTT tttggttctcggaaaactggtga | |
| NEAT1-CIR-5′-15 | *NEAT1* | caggaggaagctggtaaagacattt TTTTTTTTTTT ctcaaccTt | |
| NEAT1-CIR-3′-15 | *NEAT1* | caaccTAcA TTTTTTTTTTT ctctgaaacaggctgtcttggccac | |
| NEAT1-CIR-5′-16 | *NEAT1* | ctcgtcttaagtggtcccttа TTTTTTTTTTT ctcaaccTt | |
| NEAT1-CIR-3′-16 | *NEAT1* | caaccTAcA TTTTTTTTTTT atccagaagagcccatctaat | |
| NEAT1-CIR-5′-17 | *NEAT1* | acacctgtgacaaatgaggaac TTTTTTTTTTT ctcaaccTt | |
| NEAT1-CIR-3′-17 | *NEAT1* | caaccTAcA TTTTTTTTTTT gctcatgattttcaatcacaag | |
| NEAT1-CIR-5′-18 | *NEAT1* | tggttgttacaaagactacaat TTTTTTTTTTT ctcaaccTt | |
| NEAT1-CIR-3′-18 | *NEAT1* | caaccTAcA TTTTTTTTTTT acagtgaccacaaaaggttatt | |
| NEAT1-CIR-5′-19 | *NEAT1* | tcatccccaagtcattggttaaga TTTTTTTTTTT ctcaaccTt | |
| NEAT1-CIR-3′-19 | *NEAT1* | caaccTAcA TTTTTTTTTTT tcccaacgacagtaattgtttgca | |
| NEAT1-CIR-5′-20 | *NEAT1* | tccccatacatgcgtgactaat TTTTTTTTTTT ctcaaccTt | |
| NEAT1-CIR-3′-20 | *NEAT1* | caaccTAcA TTTTTTTTTTT caacagcatacccgagactact | |

**Table S6. DNA sequences of CIR-FISH probes targeting *NEAT1*.**

CIR-FISH probes were designed using the same target recognition sequences (TRS) as the corresponding *NEAT1* 3WJ-FISH probes, but with distinct functional domains to generate the circular architecture illustrated in Fig. S6A. Lowercase letters indicate target recognition sequences, whereas uppercase letters indicate functional domains. All sequences are listed in the 5′ to 3′ direction.

| Name | 6HB staple position | Displayed handle | Sequence (5′→3′) |
|---|---|---|---|
| H3 -ext-1 | 0[125]4[112] | H3 | ACAAACAATTCGACCACCGGAACCGCCTGACAGAATCAAGTT ttgtgggctgctgcgatttcac |
| H3 -ext-2 | 0[629]4[616] | H3 | ATGGAAATACCTACCAGAACCGCCACCCGGCATAGTAAGATA ttgtgggctgctgcgatttcac |
| H3 -ext-3 | 0[1133]4[1120] | H3 | GATGGTGGTTCCGATCAGCAGCGAAAGAGAGATTTGTATCAT ttgtgggctgctgcgatttcac |
| H3 -ext-4 | 1[112]3[125] | H3 | ATTTTGCGGAACAATCCTAATTTACGAGTCCTTATCATTCCA ttgtgggctgctgcgatttcac |
| H3 -ext-5 | 1[616]3[629] | H3 | AGCTTAGATTAAGAATCATAGGTCTGAGAAGCAACACTATCA ttgtgggctgctgcgatttcac |
| H3 -ext-6 | 1[1120]3[1133] | H3 | CTGGGGTGCCTAATAAATTTTTAGAACCTTGATTCCCAATTC ttgtgggctgctgcgatttcac |
| H3 -ext-7 | 2[139]0[126] | H3 | AAAATAATATCCCAAGAAACCACCAGAAAAGTATTAGACTTT ttgtgggctgctgcgatttcac |
| H3 -ext-8 | 2[643]0[630] | H3 | GTGAATTTATCAAACGCTGAGAAGAGTCCAGGAAAAACGCTC ttgtgggctgctgcgatttcac |
| H3 -ext-9 | 2[1147]0[1134] | H3 | CAACGCAAGGATAAGAGTGAGCTAACTCCGAAAATCCTGTTT ttgtgggctgctgcgatttcac |
| H3 -ext-10 | 3[126]5[139] | H3 | AGAACGGGTATTAACCGTAATCAGTAGCCCCTCAGAGCCGCC ttgtgggctgctgcgatttcac |
| H3 -ext-11 | 3[630]5[643] | H3 | TAACCCTCGTTTACAAAAGGAATTACGATCAGAACCGCCACC ttgtgggctgctgcgatttcac |
| H3 -ext-12 | 3[1134]5[1147] | H3 | TGCGAACGAGTAGAAACAAAGTACAACGCAGCATCGGAACGA ttgtgggctgctgcgatttcac |
| H3 -ext-13 | 4[111]2[98] H3 | H3 | TGCCTTTAGCGTCATAATCGGCTGTCTTCATGTAGAAACCAA ttgtgggctgctgcgatttcac |
| H3 -ext-14 | 4[615]2[602] | H3 | ATAAGAGCAAGAAAATTGAGTTAAGCCCAGACTACCTTTTTA ttgtgggctgctgcgatttcac |
| H3 -ext-15 | 4[1119]2[1106] | H3 | CGCCTGATAAATTGATTCCATATAACAGCTCATATATTTTAA ttgtgggctgctgcgatttcac |
| H3 -ext-16 | 5[98]1[111] | H3 | GGAACCAGAGCCACAACTCGTATTAAATTGAGTAACATTATC ttgtgggctgctgcgatttcac |
| H3 -ext-17 | 5[602]1[615] | H3 | AGTACCGCCACCCTATTTTGACGCTCAAGAAAACATAGCGAT ttgtgggctgctgcgatttcac |
| H3 -ext-18 | 5[1106]1[1119] | H3 | CGGGATCGTCACCCAATCGGCAAAATCCATAAAGTGTAAAGC ttgtgggctgctgcgatttcac |
| H3 -ext-19 | 3[126]5[139] | H3 | ACAAACAATTCGACCACCGGAACCGCCTGACAGAATCAAGTT ttgtgggctgctgcgatttcac |

**Table S7. Modified 6HB extension staple sequences used for DNA origami assembly**

The 6HB DNA origami design was adapted from a previously reported architecture (*29*). Core scaffold and staple sequences were used as described previously and are not repeated here. Only modified extension staples used in this study are listed. These staples contain the original 6HB staple-binding region (uppercase) fused to the H3 target-binding sequence (lowercase), enabling display of single-stranded H3 extensions on the assembled origami for subsequent detection experiments. All sequences are listed in the 5′ to 3′ direction.

| Name | Target | Full sequence (5'→3') | Application (Figures) |
|---|---|---|---|
| UBC-5'H3-smFISH-1 | *UBC* | GTGGGCTGCTGCGATTTCACTTT gcagagtggactctttctgg | Sensitivity validation (Fig. 3A-C); RNA-seq correlation (Fig. S7); smFISH vs 3WJ-FISH (Fig. S9) |
| UBC-5'H3-smFISH-2 | *UBC* | GTGGGCTGCTGCGATTTCACTTT gtttcccagcaaagatcaac | |
| UBC-5'H3-smFISH-3 | *UBC* | GTGGGCTGCTGCGATTTCACTTT cttccttatcttggatcttt | |
| UBC-5'H3-smFISH-4 | *UBC* | GTGGGCTGCTGCGATTTCACTTT cttcacgaagatttgcatcc | |
| UBC-5'H3-smFISH-5 | *UBC* | GTGGGCTGCTGCGATTTCACTTT ttgacattctcaatggtgtc | |
| UBC-5'H3-smFISH-6 | *UBC* | GTGGGCTGCTGCGATTTCACTTT gatggtcttaccagtcaggg | |
| UBC-5'H3-smFISH-7 | *UBC* | GTGGGCTGCTGCGATTTCACTTT gatggtcttaccagtcaggg | |
| UBC-5'H3-smFISH-8 | *UBC* | GTGGGCTGCTGCGATTTCACTTT cacctctgagacggagcacc | |
| UBC-5'H3-smFISH-9 | *UBC* | GTGGGCTGCTGCGATTTCACTTT agctgttttccggcaaagat | |
| UBC-5'H3-smFISH-10 | *UBC* | GTGGGCTGCTGCGATTTCACTTT tgttgtagtcagacagggtg | |
| UBC-5'H3-smFISH-11 | *UBC* | GTGGGCTGCTGCGATTTCACTTT ctctgctggtcaggaggaat | |
| UBC-5'H3-smFISH-12 | *UBC* | GTGGGCTGCTGCGATTTCACTTT gctttgacgttctcgatagt | |
| UBC-5'H3-smFISH-13 | *UBC* | GTGGGCTGCTGCGATTTCACTTT tggtcttgccagtgagtgtc | |
| UBC-5'H3-smFISH-14 | *UBC* | GTGGGCTGCTGCGATTTCACTTT ctctgagacggagtaccagg | |
| UBC-5'H3-smFISH-15 | *UBC* | GTGGGCTGCTGCGATTTCACTTT gccttccttatcttggatct | |
| UBC-5'H3-smFISH-16 | *UBC* | GTGGGCTGCTGCGATTTCACTTT tttgacgttctcgatggtgt | |
| UBC-5'H3-smFISH-17 | *UBC* | GTGGGCTGCTGCGATTTCACTTT ctctgagacggagcaccagg | |
| UBC-5'H3-smFISH-18 | *UBC* | GTGGGCTGCTGCGATTTCACTTT cttcacgaagatctgcattg | |
| UBC-5'H3-smFISH-19 | *UBC* | GTGGGCTGCTGCGATTTCACTTT tgcgacccaaatcccggctg | |
| EPIC1-5'H3-smFISH-1 | *EPIC1* | GTGGGCTGCTGCGATTTCACTTT gcaggagctctgagggataa | smFISH vs 3WJ-FISH (Fig. S8) |
| EPIC1-5'H3-smFISH-2 | *EPIC1* | GTGGGCTGCTGCGATTTCACTTT tgaggccacacttcacgttt | |
| EPIC1-5'H3-smFISH-3 | *EPIC1* | GTGGGCTGCTGCGATTTCACTTT ggcaggagtcgacgctacag | |
| EPIC1-5'H3-smFISH-4 | *EPIC1* | GTGGGCTGCTGCGATTTCACTTT gcctcagctgcgattgtgga | |
| EPIC1-5'H3-smFISH-5 | *EPIC1* | GTGGGCTGCTGCGATTTCACTTT aagggcaaatccatgatctc | |
| EPIC1-5'H3-smFISH-6 | *EPIC1* | GTGGGCTGCTGCGATTTCACTTT aatacagacgctgtgggcag | |
| EPIC1-5'H3-smFISH-7 | *EPIC1* | GTGGGCTGCTGCGATTTCACTTT cctctgtttgaagatctcag | |

| | | | |
|---|---|---|---|
| EPIC1-5'H3-smFISH-8 | *EPIC1* | GTGGGCTGCTGCGATTTCACTTT ttcagagtccagggctcggg | |
| EPIC1-5'H3-smFISH-9 | *EPIC1* | GTGGGCTGCTGCGATTTCACTTT gtgggatctgaaacccggag | |
| EPIC1-5'H3-smFISH-10 | *EPIC1* | GTGGGCTGCTGCGATTTCACTTT gaagaggcaccgatgctcat | |
| EPIC1-5'H3-smFISH-11 | *EPIC1* | GTGGGCTGCTGCGATTTCACTTT ccatgtaaggcatctcagag | |
| EPIC1-5'H3-smFISH-12 | *EPIC1* | GTGGGCTGCTGCGATTTCACTTT cataacactttacatacgat | |
| EPIC1-5'H3-smFISH-13 | *EPIC1* | GTGGGCTGCTGCGATTTCACTTT tgtctttccctctggaatgc | |
| EPIC1-5'H3-smFISH-14 | *EPIC1* | GTGGGCTGCTGCGATTTCACTTT cagccagtcagatttggtca | |
| EPIC1-5'H3-smFISH-15 | *EPIC1* | GTGGGCTGCTGCGATTTCACTTT gccatttattaagggccgat | |
| EPIC1-5'H3-smFISH-16 | *EPIC1* | GTGGGCTGCTGCGATTTCACTTT aatcaagcccagcttctgaa | |
| EPIC1-5'H3-smFISH-17 | *EPIC1* | GTGGGCTGCTGCGATTTCACTTT ctctgtcgcccaagggttta | |
| EPIC1-5'H3-smFISH-18 | *EPIC1* | GTGGGCTGCTGCGATTTCACTTT gcaatgaagatagcatcgaa | |
| EPIC1-5'H3-smFISH-19 | *EPIC1* | GTGGGCTGCTGCGATTTCACTTT cctgagaggctatccgtctg | |
| EPIC1-5'H3-smFISH-20 | *EPIC1* | GTGGGCTGCTGCGATTTCACTTT gagaagacaactcagtccgc | |
| EPIC1-5'H3-smFISH-21 | *EPIC1* | GTGGGCTGCTGCGATTTCACTTT actctcccactggggatgag | |
| EPIC1-5'H3-smFISH-22 | *EPIC1* | GTGGGCTGCTGCGATTTCACTTT gctcatctaagcttaatcac | |
| EPIC1-5'H3-smFISH-23 | *EPIC1* | GTGGGCTGCTGCGATTTCACTTT tcagtgtccctctttaccag | |
| EPIC1-5'H3-smFISH-24 | *EPIC1* | GTGGGCTGCTGCGATTTCACTTT tctcattgtgtcctcacacg | |
| EPIC1-5'H3-smFISH-25 | *EPIC1* | GTGGGCTGCTGCGATTTCACTTT aaggccccgtagggttcagt | |
| EPIC1-5'H3-smFISH-26 | *EPIC1* | GTGGGCTGCTGCGATTTCACTTT acatttattcctagccattc | |
| EPIC1-5'H3-smFISH-27 | *EPIC1* | GTGGGCTGCTGCGATTTCACTTT ctggtgctgccataacaaag | |
| EPIC1-5'H3-smFISH-28 | *EPIC1* | GTGGGCTGCTGCGATTTCACTTT tccaggtggaacaagagcca | |
| EPIC1-5'H3-smFISH-29 | *EPIC1* | GTGGGCTGCTGCGATTTCACTTT acaatgaggtaagggggtga | |
| EPIC1-5'H3-smFISH-30 | *EPIC1* | GTGGGCTGCTGCGATTTCACTTT tctgggtccagggaggcaga | |
| TOP2A-5'H3-smFISH-1 | *TOP2A* | GTGGGCTGCTGCGATTTCACTTT ccttggatttcttgcttgtg | RNA-seq correlation (Fig. S7); smFISH vs 3WJ-FISH (Fig. S10) |
| TOP2A-5'H3-smFISH-2 | *TOP2A* | GTGGGCTGCTGCGATTTCACTTT caccagaattcaaagctgga | |
| TOP2A-5'H3-smFISH-3 | *TOP2A* | GTGGGCTGCTGCGATTTCACTTT tctgaatcagaccagggatt | |
| TOP2A-5'H3-smFISH-4 | *TOP2A* | GTGGGCTGCTGCGATTTCACTTT ttgaccacgcggagaaggca | |

| | | |
|---|---|---|
| TOP2A-5'H3-smFISH-5 | *TOP2A* | GTGGGCTGCTGCGATTTCACTTT tcatatcccctctgaatcag |
| TOP2A-5'H3-smFISH-6 | *TOP2A* | GTGGGCTGCTGCGATTTCACTTT agcagattcagcaccaagca |
| TOP2A-5'H3-smFISH-7 | *TOP2A* | GTGGGCTGCTGCGATTTCACTTT gttatgagaggaggtgtctt |
| TOP2A-5'H3-smFISH-8 | *TOP2A* | GTGGGCTGCTGCGATTTCACTTT aagttcttgtaacttggaag |
| TOP2A-5'H3-smFISH-9 | *TOP2A* | GTGGGCTGCTGCGATTTCACTTT acgcctgatgttatttacaa |
| TOP2A-5'H3-smFISH-10 | *TOP2A* | GTGGGCTGCTGCGATTTCACTTT gctggttgtcatcatataaa |
| TOP2A-5'H3-smFISH-11 | *TOP2A* | GTGGGCTGCTGCGATTTCACTTT tagcagaatccttgccacca |
| TOP2A-5'H3-smFISH-12 | *TOP2A* | GTGGGCTGCTGCGATTTCACTTT acctttacttctcgcttgtc |
| TOP2A-5'H3-smFISH-13 | *TOP2A* | GTGGGCTGCTGCGATTTCACTTT attcctttcgatcatctatc |
| TOP2A-5'H3-smFISH-14 | *TOP2A* | GTGGGCTGCTGCGATTTCACTTT aaacgatgtcgcagaagaga |
| TOP2A-5'H3-smFISH-15 | *TOP2A* | GTGGGCTGCTGCGATTTCACTTT atctgtcataatcattatct |
| TOP2A-5'H3-smFISH-16 | *TOP2A* | GTGGGCTGCTGCGATTTCACTTT gacccacaatcttgatgata |
| TOP2A-5'H3-smFISH-17 | *TOP2A* | GTGGGCTGCTGCGATTTCACTTT tttgtctctcccaaccacac |
| TOP2A-5'H3-smFISH-18 | *TOP2A* | GTGGGCTGCTGCGATTTCACTTT tacactcagtggagtttcgg |
| TOP2A-5'H3-smFISH-19 | *TOP2A* | GTGGGCTGCTGCGATTTCACTTT ttgggaattcccttgattct |
| TOP2A-5'H3-smFISH-20 | *TOP2A* | GTGGGCTGCTGCGATTTCACTTT cttaaacttcacccagttta |
| TOP2A-5'H3-smFISH-21 | *TOP2A* | GTGGGCTGCTGCGATTTCACTTT agctcttgggttgtaaagtc |
| TOP2A-5'H3-smFISH-22 | *TOP2A* | GTGGGCTGCTGCGATTTCACTTT gcagtttatttccattaaga |
| TOP2A-5'H3-smFISH-23 | *TOP2A* | GTGGGCTGCTGCGATTTCACTTT gctcttctgaccattagtgc |
| TOP2A-5'H3-smFISH-24 | *TOP2A* | GTGGGCTGCTGCGATTTCACTTT agctgtccaaatatgagagc |
| TOP2A-5'H3-smFISH-25 | *TOP2A* | GTGGGCTGCTGCGATTTCACTTT attattccatatactaatta |
| TOP2A-5'H3-smFISH-26 | *TOP2A* | GTGGGCTGCTGCGATTTCACTTT aaacccacatttgctgggtc |
| TOP2A-5'H3-smFISH-27 | *TOP2A* | GTGGGCTGCTGCGATTTCACTTT ggagcaaaatatgttccaat |
| TOP2A-5'H3-smFISH-28 | *TOP2A* | GTGGGCTGCTGCGATTTCACTTT ccatttcgaccacctgtcac |
| TOP2A-5'H3-smFISH-29 | *TOP2A* | GTGGGCTGCTGCGATTTCACTTT cccatattatccatccatgt |
| TOP2A-5'H3-smFISH-30 | *TOP2A* | GTGGGCTGCTGCGATTTCACTTT aatctgatcagctacataat |
| TOP2A-5'H3-smFISH-31 | *TOP2A* | GTGGGCTGCTGCGATTTCACTTT gcttctcgaacattgagtat |

| | | |
|---|---|---|
| TOP2A-5'H3-smFISH-32 | *TOP2A* | GTGGGCTGCTGCGATTTCACTTT aggaaggctgtaaaatgcca |
| TOP2A-5'H3-smFISH-33 | *TOP2A* | GTGGGCTGCTGCGATTTCACTTT atgtgctggtgcccaaacct |
| TOP2A-5'H3-smFISH-34 | *TOP2A* | GTGGGCTGCTGCGATTTCACTTT tatttgaactggatacgatg |
| TOP2A-5'H3-smFISH-35 | *TOP2A* | GTGGGCTGCTGCGATTTCACTTT gtccatacaagtaatcctca |
| TOP2A-5'H3-smFISH-36 | *TOP2A* | GTGGGCTGCTGCGATTTCACTTT gagaacaagataagttcctt |
| TOP2A-5'H3-smFISH-37 | *TOP2A* | GTGGGCTGCTGCGATTTCACTTT ccatccaccatagaagggat |
| TOP2A-5'H3-smFISH-38 | *TOP2A* | GTGGGCTGCTGCGATTTCACTTT caccatgatgataagaagac |
| TOP2A-5'H3-smFISH-39 | *TOP2A* | GTGGGCTGCTGCGATTTCACTTT tgaagactttgtgtagtcca |
| TOP2A-5'H3-smFISH-40 | *TOP2A* | GTGGGCTGCTGCGATTTCACTTT ttctcgtggagggacatcaa |
| TOP2A-5'H3-smFISH-41 | *TOP2A* | GTGGGCTGCTGCGATTTCACTTT tcaaggtctgacacgacact |
| TOP2A-5'H3-smFISH-42 | *TOP2A* | GTGGGCTGCTGCGATTTCACTTT attttgcccgaggagccaca |
| TOP2A-5'H3-smFISH-43 | *TOP2A* | GTGGGCTGCTGCGATTTCACTTT taattacttaaaataatcgc |
| TOP2A-5'H3-smFISH-44 | *TOP2A* | GTGGGCTGCTGCGATTTCACTTT actatttagacagtattata |
| TOP2A-5'H3-smFISH-45 | *TOP2A* | GTGGGCTGCTGCGATTTCACTTT caaactgctctagttttagc |
| TOP2A-5'H3-smFISH-46 | *TOP2A* | GTGGGCTGCTGCGATTTCACTTT atgctctatctcatatctac |
| TOP2A-5'H3-smFISH-47 | *TOP2A* | GTGGGCTGCTGCGATTTCACTTT aaagttctatctgatggtaa |
| TOP2A-5'H3-smFISH-48 | *TOP2A* | GTGGGCTGCTGCGATTTCACTTT agatgaagaacatgagcaat |
| TOP2A-5'H3-smFISH-49 | *TOP2A* | GTGGGCTGCTGCGATTTCACTTT ttgactgtgtcaagttatag |
| TOP2A-5'H3-smFISH-50 | *TOP2A* | GTGGGCTGCTGCGATTTCACTTT atgttgtccccgagtcttct |

**Table S8. DNA sequences of gene-specific probes used for conventional smFISH experiments.**

Each probe consists of a readout sequence (uppercase) fused to a Target-recognition sequence (TRS; lowercase) targeting the mRNAs *UBC* and *TOP2A* or the long noncoding RNA (lncRNA) *EPIC1*. All sequences are listed in the 5′ to 3′ direction.

| Name | Sequence (5’→3’) | Modification | Conjugation chemistry | Target |
| --- | --- | --- | --- | --- |
| H1 handle | TTGCGCGATTGACCGTCTCGTT | 5′-Azide | Click chemistry | Anti-rabbit IgG |
| H3 handle | TTGTGGGCTGCTGCGATTTCAC | 5′-Azide | Click chemistry | Anti-mouse IgG |

**Table S9. DNA sequences used for antibody–DNA conjugation.**

DNA oligonucleotides bearing 5′-azide modifications were conjugated to secondary antibodies via copper-free click chemistry. The H3 sequence listed here is identical to the sequence used for origami display (Table S7) and serves as the target sequence for detection. All sequences are listed in the 5′ to 3′ direction.

| Antibody | Antibody type | Host species | Clonality | Supplier | Catalog number | Conjugation | Dilution |
|---|---|---|---|---|---|---|---|
| α- Tubulin | Primary | Rabbit | Monoclonal | Abcam | ab52866 | None | 1:200 |
| α- Tubulin | Primary | Mouse | Monoclonal | Cell Signaling Technology | 3873T | None | 1:200 |
| TNFα | Primary | Mouse | Monoclonal | Novus Biologicals | NBP2-45331 | None | 1:100 |
| αSMA | Primary | Rabbit | Monoclonal | Cell Signaling Technology | 19245T | None | 1:200 |
| EZH2 | Primary | Rabbit | Monoclonal | Cell Signaling Technology | 16098S | None | 1:400 |
| EZH2 | Primary | Mouse | Monoclonal | Proteintech | 66476-1-Ig | None | 1:200 |
| RNAPII | Primary | Mouse | Monoclonal | Abcam | ab5408 | None | 1:400 |
| Anti-rabbit IgG | Secondary | Goat | Polyclonal | Jackson ImmunoResearch Laboratories | 111-005-144 | DNA oligo (3WJ-IF); Alexa Fluor 647 (Conv- IF) | 1:200 |
| Anti-mouse IgG | Secondary | Goat | Polyclonal | Jackson ImmunoResearch Laboratories | 115-005-166 | DNA oligo (3WJ-IF); Alexa Fluor 647 (Conv- IF) | 1:200 |

**Table S10. Antibodies used in this study**

Primary antibodies and secondary antibodies used for immunofluorescence (IF) and 3WJ-based IF experiments. Secondary antibodies were either conjugated to DNA oligonucleotides for 3WJ amplification or labeled with fluorophores for conventional IF. Dilutions correspond to those used for imaging experiments.

| Figure | Imaging modality | Microscope | Laser lines (nm) | Frames | Pixel size (nm) |
|---|---|---|---|---|---|
| Figure 1D (*i*) | STORM (super-resolution) | Custom-built STORM microscope | 642 (excitation); 405 (activation) | ~20,000 | 13 |
| Figure 1D (*ii*), 5F, S17A-B | Mesoscopic large-FOV | Custom-built mesoscope | 405/ 530/ 650 | _ | 320 |
| Figure 1D (*iii*), S13A-B | Widefield (fluorescence) | Custom-built STORM microscope | 405/ 561/ 642 | _ | 130 |
| Figure 1D (*iv*), 6C, S4B (right), S17A-B | STORM (super-resolution) | Custom-built STORM microscope | 561/ 642 (excitation); 405 (activation) | ~40,000 | 13 |
| Figure 2A, 3E, S5C | STORM (super-resolution) | Custom-built STORM microscope | 642 (excitation); 405 (activation) | ~20,000 | 13 |
| Figure 2E, 3A, 4A, 4H | Widefield (fluorescence) | Custom-built STORM microscope | 405/ 642 | _ | 130 |
| Figure 5B, 5C | Mesoscopic large-FOV | Custom-built mesoscope | 405/ 650 | _ | 320 |
| Figure 5F | Mesoscopic large-FOV | Custom-built mesoscope | 405/ 530/ 650 | _ | 320 |
| Figure 6A | Widefield (fluorescence) | Custom-built STORM microscope | 405/ 642 | _ | 130 |
| Figure S2B, S2E, S4B (left), S3B | Widefield (fluorescence) | Custom-built STORM microscope | 405/ 642 | _ | 130 |
| Figure S4B (right), S14A-B | STORM (super-resolution) | Custom-built STORM microscope | 642 (excitation); 405 (activation) | ~20,000 | 13 |
| Figure S6; S8; S9; S10; S11; S12A-K | Widefield (fluorescence) | Custom-built STORM microscope | 405/ 642 | _ | 130 |
| Figure S15, S20C-D | Widefield (fluorescence) | Custom-built STORM microscope | 642 | _ | 130 |
| Figure S16C | Widefield (fluorescence) | Custom-built STORM microscope | 405/ 642 | _ | 130 |
| Figure S18A, S19A | Widefield (fluorescence) | Custom-built STORM microscope | 561/ 642 | _ | 130 |

**Table S11. Imaging acquisition conditions**

A custom-built high-resolution (NA = 1.42) wide-field fluorescence microscope was used for both super-resolution (STORM) imaging and widefield fluorescence imaging, with pixel sizes of 13 nm and 130 nm, respectively. Mesoscopic ultra-large-field-of-view (FOV) imaging was performed using a previously described mesoscopic large-FOV fluorescence imaging system (objective NA = 0.5) with a pixel size of 320 nm. Image acquisition was controlled using LabVIEW, and image reconstruction and processing were performed using custom MATLAB-based pipelines as described in the Methods and corresponding references.

| Figure | Comparison | Test | n | Mean ± SD | p-value |
|---|---|---|---|---|---|
| **Figure 2B** | Mean WF intensity per structure (3WJ vs 4C vs 0C) | Welch ANOVA (Dunnett T3) | 226, 117, 71 | 268.1 ± 123.6 vs 133.7 ± 58.9 vs 66.9 ± 53.7 | <0.0001 |
| **Figure 2C** | Mean localizations per structure (3WJ vs 4C vs 0C) | Welch ANOVA (Dunnett T3) | 226, 117, 71 | 262.2 ± 124.7 vs 107.5 ± 49.32 vs 66.71 ± 53.70 | <0.0001 |
| **Figure 2F** | Mean signal intensity (A vs S) | Welch's t-test (two-tailed) | 60, 31 | 357.7 ± 90.32 vs 181.4 ± 25.08 | <0.0001 |
| **Figure 2G** | Mean spots (A vs S) | Welch's t-test (two-tailed) | 60, 30 | 394.0 ± 119.3 vs 270.8 ± 103.6 | <0.0001 |
| **Figure 3B** | Mean signal intensity (smFISH vs 3WJ-FISH) | Mann–Whitney U test (two-tailed) | 127, 131 | 64.52 ± 16.06 vs 142.0 ± 35.80 | <0.0001 |
| **Figure 3C** | Mean spots (smFISH vs 3WJ-FISH) | Mann–Whitney U test (two-tailed) | 127, 131 | 264.2 ± 100.5 vs 344.6 ± 100.7 | <0.0001 |
| **Figure 3F** | Mean WF intensity per structure (1× vs 8× vs 16×) | Welch ANOVA (Dunnett's T3) | 171, 89, 71 | 34.94 ± 14.23 vs 63.37 ± 33.18 vs 137.9 ± 71.62 | <0.0001 |
| **Figure 3G** | Mean localizations per structure (1× vs 8× vs 16×) | Welch ANOVA (Dunnett's T3) | 171, 89, 71 | 36.47 ± 18.98 vs 81.30 ± 35.95 vs 127.7 ± 49.80 | <0.0001 |
| **Figure 3H** | Mean cluster size (1× vs 8× vs 16×) | Welch ANOVA (Dunnett's T3) | 133, 188, 201 | 38.41 ± 5.56 vs 49.20 ± 5.73 vs 55.23 ± 8.70 | <0.0001 |
| **Figure 4B** | Total background (5 nt vs 9 nt vs 15 nt vs 20 nt) | Kruskal–Wallis test (Dunn's multiple comparisons) | 641, 5537, 12213, 14187 | 4.023 ± 3.633 vs 11.54 ± 14.92 vs 84.64 ± 79.68 vs 106.9 ± 91.22 | <0.0001 |
| **Figure 4C** | Mean background (5 nt vs 9 nt vs 15 nt vs 20 nt) | Kruskal–Wallis test (Dunn's multiple comparisons) | 52, 44, 46, 41 | 0.00 ± 0.00 vs 0.00 ± 0.00 vs 219.8 ± 20.0 vs 224.2 ± 15.66 | <0.0001 |
| **Figure 4D** | Background spots per cell (5, 9, 15, and 20 nt) | Kruskal–Wallis test (Dunn's multiple comparisons) | 52, 52, 46, 41 | 0.00 ± 0.00 vs 0.19 ± 0.53 vs 45.59 ± 29.33 vs 90.78 ± 52.54 | <0.0001 |
| **Figure 4E** | Mean signal intensity (5 nt vs 9 nt) | Welch's t-test (two-tailed) | 62, 44 | 260.0 ± 48.08 vs 241.1 ± 19.77 | 0.0155 |
| **Figure 4F** | Mean spots (5 nt vs 9 nt) | Welch's t-test (two-tailed) | 65, 43 | 11.14 ± 5.81 vs 115.1 ± 54.77 | <0.0001 |
| **Figure 4G** | Signal intensity (Pos, Neg5, Neg3, Un5, Un3, Un) | Welch ANOVA (Dunnett's T3 multiple comparisons) | 20, 18, 49, 22, 18, 21 | 1630 ± 467.2 vs 253.1 ± 48.7 vs 279.4 ± 43.6 vs 256.3 ± 35.1 vs 458.3 ± 131.8 vs 497.8 ± 100.0 | <0.0001 |
| **Figure 5D** | α-tubulin fluorescence intensity per cell (3WJ-IF vs Conv IF) | Welch's t-test (two-tailed) | 7595, 4880 | 24152 ± 6078 vs 5775 ± 1600 | <0.0001 |
| **Figure 5E** | TNFα fluorescence intensity per TNFα-positive object (3WJ-IF vs Conv IF) | Mann–Whitney test (two-tailed) | 2577, 8233 | 21660 ± 6212 vs 4469 ± 1500 | <0.0001 |
| **Figure 6B** | Mean EZH2 signal intensity (IF vs IF+FISH vs 3WJ-IF+FISH) | Welch ANOVA (Dunnett's T3 multiple comparisons) | 34, 28, 28 | 608.8 ± 135.5 vs 199.9 ± 12.25 vs 11844 ± 2446 | <0.0001 |
| **Figure 6D** | EZH2 cluster density (IF+FISH vs 3WJ-IF+FISH) | Welch's t-test (two-tailed) | 15, 38 | 0.6030 ± 0.3623 vs 3.610 ± 2.893 | <0.0001 |

| | | | | | |
|---|---|---|---|---|---|
| **Figure 6E** | EZH2 localization per cell within Xi (IF+FISH vs 3WJ-IF+FISH) | Welch's t-test (two-tailed) | 15, 38 | 70.67 ± 21.82 vs 89.84 ± 25.17 | 0.0100 |
| **Figure S2C** | Mean spots (N-HBS vs H-HBS) | Welch's t-test (two-tailed) | 5, 45 | 377.0 ± 149.3 vs 745.5 ± 271.9 | 0.0047 |
| **Figure S2D** | Mean signal (N-HBS vs H-HBS) | Welch's t-test (two-tailed) | 5, 45 | 37.70 ± 19.01 vs 95.76 ± 17.65 | 0.0015 |
| **Figure S2F** | Mean signal (N-HBS vs H-HBS) | Welch's t-test (two-tailed) | 66, 87 | 2211 ± 402.6 vs 5225 ± 748.1 | <0.0001 |
| **Figure S3C** | Mean RNAPII signal intensity per cell (+LNA vs −LNA) | Welch's t-test (two-tailed) | 18, 20 | 17492 ± 2763 vs 2452 ± 452.0 | <0.0001 |
| **Figure S4C** | Mean WF intensity per structure (3WJ vs 4C vs 0C vs π vs CIR) | Welch ANOVA (Dunnett's T3 multiple comparisons) | 226, 117, 71, 43, 72 | 268.1 ± 123.6 vs 133.7 ± 58.96 vs 66.86 ± 53.70 vs 49.32 ± 22.93 vs 75.59 ± 33.77 | <0.0001 |
| **Figure S4D** | Mean localizations per structure (3WJ vs 4C vs 0C vs π vs CIR) | Welch ANOVA (Dunnett's T3 multiple comparisons) | 226, 117, 71, 43, 72 | 262.2 ± 124.7 vs 107.5 ± 49.32 vs 66.71 ± 53.70 vs 132.6 ± 42.41 vs 113.1 ± 43.37 | <0.0001 |
| **Figure S5D** | Distances between adjacent localization clusters (d1 and d2) | Descriptive statistics | 28 | 152 ± 10 nm (d1); 153 ± 11 nm (d2) | N/A |
| **Figure S6B** | *NEAT1* spots per nucleus (Z vs CIR vs 3WJ) | Kruskal–Wallis test with Dunn's multiple comparisons | 38, 28, 32 | 0.00 ± 0.00 vs 1.50 ± 1.75 vs 7.88 ± 3.80 | <0.0001 (3WJ vs CIR; 3WJ vs Z) |
| **Figure S6C** | *NEAT1* spot intensity per nucleus (Z vs CIR vs 3WJ) | Kruskal–Wallis test with Dunn's multiple comparisons | 1, 18, 32 | 0.00 ± N/A vs 263.1 ± 54.39 vs 436.4 ± 96.16 | <0.0001 (3WJ vs CIR); 0.0513 (3WJ vs Z) |
| **Fig. S10B** | Mean *TOP2A* signal intensity per cell (smFISH vs 3WJ-FISH) | Mann–Whitney U test (two-tailed) | 24, 146 | 14.46 ± 6.29 vs 165.4 ± 23.90 | <0.0001 |
| **Fig. S10C** | Mean *TOP2A* spots per cell (smFISH vs 3WJ-FISH) | Mann–Whitney U test (two-tailed) | 24, 146 | 16.25 ± 9.16 vs 47.36 ± 42.44 | <0.0001 |
| **Fig. S11B** | Mean *TOP2A* signal intensity per cell (smFISH vs 3WJ-FISH) | Mann–Whitney U test (two-tailed) | 15, 103 | 19.71 ± 3.45 vs 50.65 ± 13.93 | <0.0001 |
| **Fig. S11C** | Mean *TOP2A* spots per cell (smFISH vs 3WJ-FISH) | Mann–Whitney U test (two-tailed) | 15, 103 | 66.73 ± 26.38 vs 100.80 ± 54.52 | 0.0169 |
| **Fig. S15B** | Mean nuclear EZH2 intensity per cell (16× CIR-IF vs 16× 3WJ-IF) | Welch's t-test (two-tailed) | 55, 55 | 465.8 ± 80.40 vs 1374 ± 262.6 | <0.0001 |
| **Figure S16D** | Mean signal intensity (3WJ-IF vs Neg5 vs Neg3) | Welch ANOVA (Dunnett's T3 multiple comparisons) | 26, 37, 28 | 5405 ± 1297 vs 785.2 ± 113.9 vs 216.4 ± 20.49 | <0.0001 |
| **Figure S18B** | Mean RNAPII signal intensity per cell (Conv IF vs 3WJ-IF) | Welch's t-test (two-tailed) | 70, 60 | 571.0 ± 141.0 vs 10911 ± 3934 | <0.0001 |

**Table S12. Summary of statistical analyses for all quantified data in this study.**

Statistical comparisons, sample sizes (n), mean values ± standard deviation (SD), and corresponding P values are reported for each quantified dataset. For comparisons involving more than two groups, significance was assessed using Welch's ANOVA followed by Dunnett's T3 multiple comparisons test. For pairwise comparisons, two-tailed unpaired Welch's t-tests were used. Non-parametric tests (Kruskal–Wallis with Dunn's test, Mann–Whitney U test) were used where indicated (Figs. S6, S10, S11). All tests were performed as described in the Methods. Exact P values or thresholds are indicated for each comparison.

| Method | Gene name | Cell line | Mean spots/cell | SD | Median | Number of cells (n) | RNA-seq expression (TPM) | Ensembl ID |
|---|---|---|---|---|---|---|---|---|
| 3WJ-FISH | *EPIC1* | MCF7 | 1.77 | 0.89 | 2.00 | 52 | 1.39 | ENSG00000224271 |
| | *TOP2A* | MCF7 | 47.36 | 42.44 | 36.00 | 146 | 57.21 | ENSG00000131747 |
| | *TOP2A* | HCT116 | 100.80 | 54.52 | 93.00 | 103 | 89.87 | ENSG00000131747 |
| | *TOP2A* | HeLa | 156.00 | 56.42 | 148.00 | 88 | 157.54 | ENSG00000131747 |
| | *TOP2A* | U2OS | 240.10 | 116.20 | 225.50 | 106 | 279.03 | ENSG00000131747 |
| | *UBC* | HeLa | 344.60 | 100.70 | 331.00 | 131 | 416.20 | ENSG00000150991 |
| smFISH | *TOP2A* | MCF7 | 16.25 | 9.16 | 14.50 | 24 | 57.21 | ENSG00000131747 |
| | *TOP2A* | HCT116 | 66.73 | 26.38 | 62.00 | 15 | 89.87 | ENSG00000131747 |
| | *UBC* | HeLa | 264.20 | 100.50 | 247.00 | 127 | 416.20 | ENSG00000150991 |

**Table S13. Descriptive statistics of RNA spots per cell detected by 3WJ-FISH and conventional smFISH, with corresponding RNA-seq expression values used for correlation analyses in Figure 3D and Fig. S7.**

RNA spots per cell were quantified from 3WJ-FISH and conventional smFISH images for the indicated genes across multiple cell lines. Values are reported as mean spots per cell, standard deviation (SD), and median, with n indicating the number of cells analyzed. Corresponding RNA-seq expression values were obtained from the Cancer Cell Line Encyclopedia (CCLE; DepMap Public 22Q1) and are reported in transcripts per million (TPM). Ensembl gene IDs are provided for each target. The 3WJ-FISH and smFISH measurements were used for the correlation analyses shown in Fig. 3D and Fig. S7, respectively.

| Category | Reagent/Material | Catalog# | Vendor |
| --- | --- | --- | --- |
| Buffers and solutions | IHC Antigen Retrieval Solution, 10X Low pH | 00-4955-58 | Invitrogen (Thermo Fisher Scientific) |
| | PBS - Phosphate-Buffered Saline (10X) pH 7.4, RNase-free | AM9625 | Invitrogen (Thermo Fisher Scientific) |
| | SSC (20X), RNase-free | AM9763 | Invitrogen (Thermo Fisher Scientific) |
| | TAE (50X), TRIS + acetate + EDTA | J63931.K3 | Thermo Scientific |
| | Tris (1 M), pH 8.0, RNase-free | AM9856 | Invitrogen (Thermo Fisher Scientific) |
| | UltraPure™ Distilled Water, RNase-and DNase-free | 10977-015 | Invitrogen (Thermo Fisher Scientific) |
| Chemicals | 16% Formaldehyde solution | 28908 | Thermo Scientific |
| | Ammonium chloride | 213330 | Sigma-Aldrich |
| | Bovine serum albumin, Ultrapure | 9048-46-8 | American Bio |
| | Chrome Alum-Gelatin Adhesive | 1033B | Newcomer Supply |
| | Chrome Alum-Gelatin Adhesive, 250ml | NC1692262 | Newcomer Supply |
| | Citric acid | C0759-100G | Sigma-Aldrich |
| | Dimethylsulfoxide (DMSO; anhydrous) | D12345 | Invitrogen (Thermo Fisher Scientific) |
| | Dulbecco's modified Eagle's medium | 11885084 | Gibco |
| | Ethanol for molecular biology, RNase-and DNase-free | 3916EA | Decon Labs |
| | Ethylenediaminetetraacetic acid (EDTA), BioUltra, anhydrous | EDS-100G | Sigma-Aldrich |
| | Fetal bovine serum | | Gibco, Invitrogen |
| | Formamide, Deionized | S4117 | Millipore-Sigma |
| | Glucose | G7021-1KG | Sigma-Aldrich |
| | Glycerol for molecular biology | G5516 | Sigma-Aldrich |
| | Guanidinum hydrochloride | G3272 | Sigma-Aldrich |
| | Hydrochloric acid, 36.5 to 38.0% | A144-500 | Fisher Chemical |
| | Hydrogen peroxide solution, 30% stabilized | JT2189-1 | VWR (J.T.Baker) |
| | Magnesium chloride ($MgCl_2$) | M8266-100G | Sigma-Aldrich |
| | N, N-Dimethylformamide (DMF) solvent | D4551-250ML | Sigma-Aldrich |
| | Penicillin-Streptomycin | 15140148 | Gibco |
| | Poly-D-lysine hydrobromide, Mol wt 30,000-70,000 | P7886-10MG | Sigma-Aldrich |
| | RNaseZap™ RNase Decontamination Solution | AM9780 | Thermo Fisher Scientific |
| | Sodium bicarbonate (NaHCO3) | S5761 | Sigma-Aldrich |
| | Sodium chloride (NaCl) | S9888-500G | Sigma-Aldrich |
| | Sodium Hydroxide (Pallets) | 7708 | MallincKrodt AR ® |
| | Triton X-100 | T8787-100ml | Sigma-Alrich |
| | Tween® 20 | P1379 | Millipore-Sigma |
| | Xylenes | 534056 | Sigma-Aldrich |
| Chromatography and purification | Amicon Ultra 0.5ml filters with 50KDa molecular cut-off membranes | UCF505096 | Merck/EMD Millipore |
| | Amicon® Ultra centrifuge filter (50KDa MWCO) | UFC505024 | Sigma-Aldrich |
| | Cytiva NAP™ columns NAP-5 | 45-000-151 | Fisher scientific |
| | NAP-5 size-exclusion columns | 17–0853-02 | GE Healthcare |
| | Zeba™ Spin Desalting columns, 7K MWCO | 89877 | Thermo Fisher Scientific |
| Conjugation reagents | biotinylated bovine albumin | A4503 | Sigma-Aldrich |
| | DBCO-sulfo-NHS esters | CLK-A124-10 | Jena Bioscience |
| | Streptavidin | S888 | Thermo Fisher Scientific |
| Consumables | Square cover glasses #1.5 (22mmx22mm) | 12-541-016 | Thermo Fisher Scientific |

| | | | |
|---|---|---|---|
| | Advanced PAP pen | Z672548 | Millipore-Sigma |
| | SecureSeal™ hybridization chambers (SA50) | 621502 | Grace Bio-Labs |
| | SecureSeal™ hybridization chambers (SA8R-0.5) | 621505 | Grace Bio-Labs |
| Enzymes and inhibitors | Catalase | C3155-50MG | Millipore-Sigma |
| | Glucose oxidase | G7141-50KU | Millipore-Sigma |
| | RNase inhibitor, Murine | M0314L | New England Biolabs (NEB) |
| | Ribonucleoside Vanadyl Complexes (RVC) | S1402S | New England Biolabs (NEB) |
| Fluorophores and dyes | 4',6-diamidino-2-phenylindole (DAPI) | D3571 | Invitrogen (Thermo Fisher Scientific) |
| | Alexa 647 carboxylic acid succinimidyl ester | A20006 | Thermo Fisher Scientific |
| | FluoSpheres™ carboxylate-modified microspheres (0.1 µm, yellow–green) | F8803 | Invitrogen (Thermo Fisher Scientific) |
| Instruments | Hybridization oven Hybaid HS9360 | HS9360 | Thermo Fisher Scientific |
| | NanoDrop™ 2000 spectrophotometers | ND2000 | Fisher Scientific |
| | Retriever 2100 (2100 Antigen Retriever) | 62700-10 | Electron Microscopy Sciences (EMS) |
| | Thermocycler | T100 | Bio-Rad |
| Nucleic acid reagents | Denhardt's Solution (50X) | 750018 | Thermo Fisher Scientific |
| | Dextran sulfate sodium salt from Leuconostoc spp. | D8906 | Sigma-Aldrich |
| | Heparin sodium salt from porcine intestinal mucosa | H3393-10KU | Millipore-Sigma |
| | M13mp18 ssDNA scaffold | N4040s | New England Biolabs (NEB) |
| | Yeast tRNA (10 mg/mL) | AM7119 | Invitrogen (Thermo Fisher Scientific) |

**Table S14. Reagents and materials used in this study.**
